# Spatial organization of PI3K-PI(3,4,5)P3-AKT signaling by focal adhesions

**DOI:** 10.1101/2024.07.05.602013

**Authors:** Jing Wang, Zhengyang An, Zhongsheng Wu, Wei Zhou, Pengyu Sun, Piyu Wu, Rui Xue, Song Dang, Xue Bai, Wenxu Wang, Rongmei Chen, Yongtao Du, Pei Huang, Sin Man Lam, Youwei Ai, Suling Liu, Guanghou Shui, Zhe Zhang, Zheng Liu, Jianyong Huang, Xiaohong Fang, Kangmin He

## Abstract

The class I PI3K-AKT signaling pathway is the master regulator of cell survival, growth, and proliferation, and among the most frequently mutated pathways in cancer. However, where and how the PI3K-AKT signaling is spatially activated and organized in mammalian cells remain poorly understood. Here, we identified focal adhesions (FAs) as the subcellular signaling hubs organizing the activation of PI3K-PI(3,4,5)P_3_-AKT signaling in mammalian cells. We found that class IA PI3Ks are preferentially and dynamically recruited to FAs for activation, resulting in localized production of the critical signaling lipid PI(3,4,5)P_3_ around FAs. As the effector protein of PI(3,4,5)P_3_, AKT molecules are dynamically recruited around FAs for activation. Mechanistically, the spatial recruitment/activation of PI3K-PI(3,4,5)P_3_-AKT cascade are regulated by the activated FAK. Furthermore, combined inhibition of class I PI3K and FAK results in a more potent inhibitory effect on cancer cells. Thus, our results unveil a growth-factor independent, compartmentalized organization mechanism for PI3K-PI(3,4,5)P_3_-AKT signaling.

## INTRODUCTION

The PI3K-AKT signaling pathway is one of the most critical and extensively investigated cell signaling pathways. It is the central regulator of various cellular processes including cell growth, proliferation, metabolism, and survival. Hyperactivation of PI3K-AKT signaling is highly related to a significant number of human diseases especially cancers (Fruman et al., 2017; Riehle et al., 2013). Notably, *PIK3CA*, which encodes the catalytic subunit p110α of class I PI3K, is the most frequently mutated oncogene across human cancers (Bailey et al., 2018; Lawrence et al., 2014; Vasan et al., 2019). Thus, the PI3K-AKT pathway has have been the focus of extensive anticancer drug discovery efforts over decades (Castel et al., 2021).

Class I PI3Ks are lipid kinases composed of a p110 catalytic subunit (p110α, p110β or p110δ for class IA, and p110γ for class IB) and a regulatory subunit. The catalytic subunits of class IA PI3K are stabilized and inactivated by the p85-like regulatory subunits in the cytosol (Bilanges et al., 2019; Burke, 2018; Rathinaswamy and Burke, 2020; Vadas et al., 2011). The canonical view, derived from extensive structural and biochemical studies, suggests that class I PI3Ks are mainly activated downstream of receptor tyrosine kinases (RTKs) or G-protein-coupled receptors (GPCRs) (Bilanges et al., 2019; Burke, 2018; Liu et al., 2021; Rathinaswamy and Burke, 2020; Vadas et al., 2011). The spatial recruitment and activation of class I PI3K at specific cellular membranes determines where and how PI3K-AKT signaling is activated and regulated (Ebner et al., 2017a; Thapa et al., 2020). Surprisingly, as the direct visualization of the dynamic spatial targeting of endogenous class I PI3K in live mammalian cells has not been achieved, the precise subcellular localization and regulatory mechanism of class I PI3K, especially p110α with distinct hotspot activating mutations, have remained elusive. This has impeded the mechanistic understanding of how the enzyme activity of class I PI3K is regulated in normal cells and how oncogenic hotspot mutations hyperactive class I PI3K-ATK signaling in cancer cells.

*PIK3CA*, the most frequently mutated oncogene, and *PTEN*, the second most frequently mutated tumor suppressor gene after *TP53* (Bailey et al., 2018; Lawrence et al., 2014), respectively regulate the generation and degradation of the critical signaling lipid PI(3,4,5)P_3_. Activating mutations in PI3K or loss-of-function mutations in PTEN result in an aberrant elevation of PI(3,4,5)P_3_, promoting cancer cell survival, proliferation, and driving tumorigenesis (Isakoff et al., 2005; Kang et al., 2005; Riehle et al., 2013; Samuels et al., 2005). Thus, PI(3,4,5)P_3_ is rigorously controlled at an extremely low level in normal cells under basal conditions (Balla, 2013; Riehle et al., 2013). Nevertheless, PI(3,4,5)P_3_ plays critical signaling roles by recruiting and activating various effector proteins, including the serine/threonine protein kinase AKT, at cellular membranes (Balla, 2013; Riehle et al., 2013). Binding to PI(3,4,5)P_3_ triggers the allosteric activation of AKT, which subsequently phosphorylates over 100 substates to regulate a myriad of essential cellular functions (Ebner et al., 2017b; Guo et al., 2015; Lucic et al., 2018; Manning and Toker, 2017). By hyperactivating cells with growth factors, previous studies have revealed that the recruitment of AKT to the plasma membrane or intracellular membranes is crucial for AKT signaling activation (Gao et al., 2011; Manning and Toker, 2017; Sugiyama et al., 2019). However, the precise subcellular compartment where class I PI3K is recruited, PI(3,4,5)P_3_ is generated, and thus AKT is recruited/activated in cells under basal or physiological conditions, remains largely unknown. Furthermore, it remains unclear how the kinase activity of PI3K and the phosphatase activity of PTEN are spatially and temporally coordinated to sustain a low level of PI(3,4,5)P_3_ and AKT activation for cell survival and growth.

Therefore, even though the PI3K-PI(3,4,5)P_3_-AKT pathway is among the most critical and most extensively studied signaling pathways, unraveling the dynamic molecular signaling events during PI3K-PI(3,4,5)P_3_-AKT signaling activation within live contexts remains highly challenging. In this study, through the development of new tools and the utilization of innovative methodologies, including lipid sensor design, genome editing, single-molecule imaging, and quantitative imaging analysis, we unexpectedly discovered that FAs are the long-sought-after subcellular compartment organizing the recruitment and activation of class IA PI3K, generation of PI(3,4,5)P_3_, and activation of AKT signaling in mammalian cells.

## RESULTS

### Class IA PI3Ks are dynamically and preferentially recruited to FAs

The surprising lack of clarity regarding the dynamic subcellular recruitment and localization of class I PI3K has impeded our understanding of the spatial organization of class I PI3K-AKT signaling in live mammalian cells. Previous studies, which included the overexpression of GFP-fused p110α and p110β, microinjection of dye-labeled p110α/p85α complex, or immunostaining of p110α and p85α with antibodies in fixed cells, have been unsuccessful in uncovering the precise dynamic subcellular localization of class I PI3K (Layton et al., 2014; Salamon and Backer, 2013; Singh et al., 2016; Thapa et al., 2020; Yip et al., 2004). To tackle this challenge, we chose to label the endogenous catalytic p110α subunit with the bright fluorescent protein mNeonGreen (Shaner et al., 2013) in human breast cancer SUM159 cells using the CRIPSR-Cas9 genome editing technique (Figure S1A). To capture the lipid kinase-catalyzed enzymatic processes that involve a very small number of enzyme molecules, we employed single-molecule quantitative imaging. By imaging the endogenous p110α molecules in gene-edited cells expressing p110α-mNeonGreen^+/-^ at a fast image rate (10 Hz) using total internal reflection fluorescence (TIRF) microscopy with single-molecule sensitivity, we unexpectedly uncovered that p110α-mNeonGreen molecules were recruited to the plasma membrane as individual, dispersed spots (Figure 1A; Video S1). Each fluorescent spot represents a single p110α molecule (Figure 1A). These spots appeared abruptly and then quickly dissociated from the plasma membrane. By creating a maximum-intensity projection of these p110α-mNeonGreen molecules over a time series, we observed well-structured patterns at the plasma membrane (Figure 1A). These structured patterns remained evident when overlaying the identified individual p110α molecules throughout the time series (Figure 1A; Video S1).

**Figure 1.**
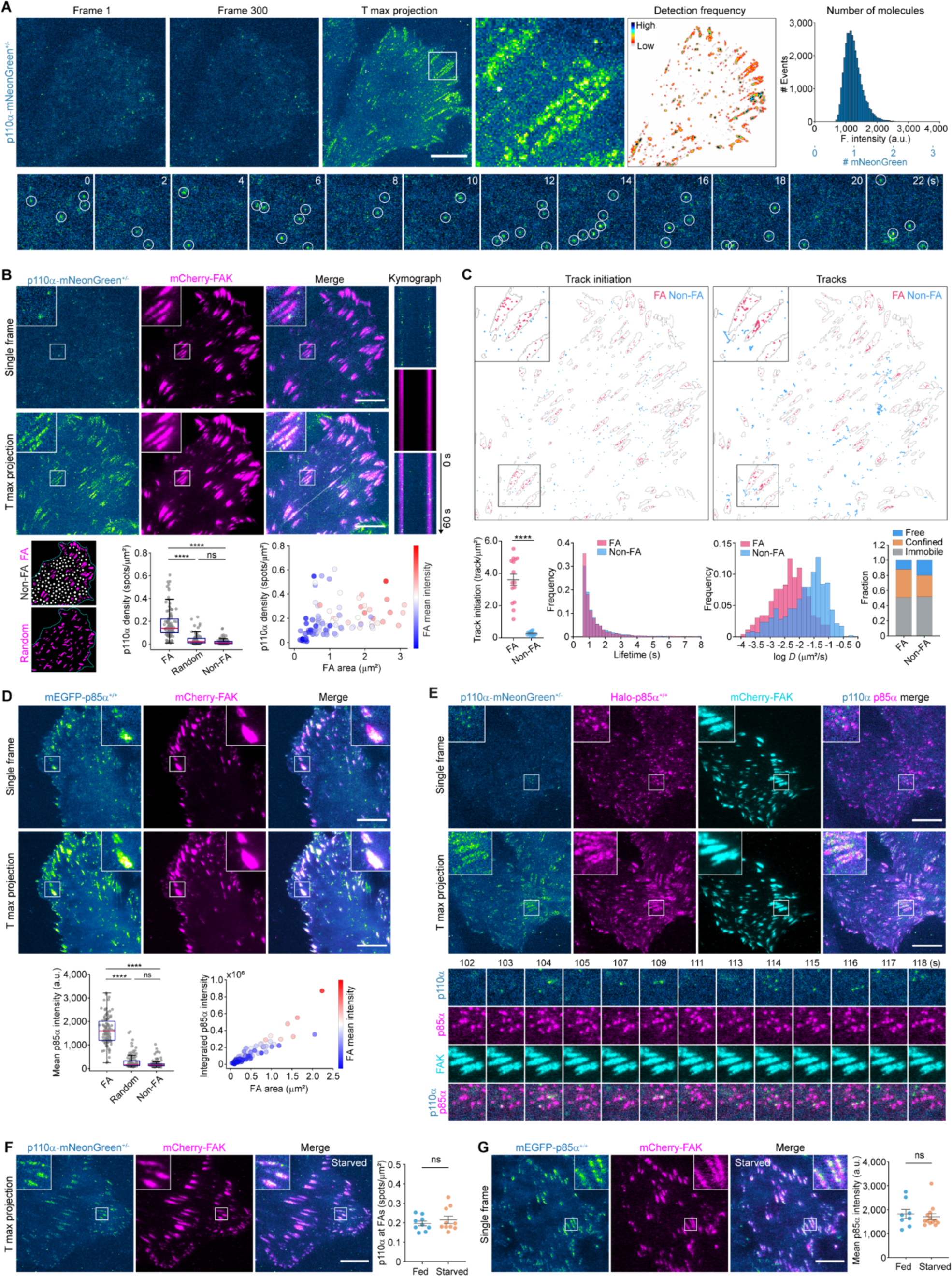
Endogenous p110α and p85α are dynamically and preferentially recruited to FAs. (A) Bottom surfaces of gene-edited p110α-mNeonGreen^+/-^ cells were imaged at 0.1-s intervals for 601 frames by TIRF microscopy with single mNeonGreen sensitivity. From left to right: Images of the first and the 300th frames of a representative time series; the maximum-intensity projection of the time series (T max projection); the detection frequency of individual p110α-mNeonGreen molecules in the time series; the distribution of intensity and number of detected p110α-mNeonGreen molecules from 12 cells. The montage shows the indicated frames of the boxed region. SUM159 cells were used unless otherwise noted. (B) p110α-mNeonGreen^+/-^ cells transiently expressing mCherry-FAK were imaged at 0.2-s intervals for 601 frames. Top left: Images of a single frame and the maximum-intensity projection of a representative time series. Top right: Kymographs generated along the line on the merged maximum-intensity projection image show the dynamic recruitment of p110α-mNeonGreen to FAs. From left to right in the bottom panels: Images showing FAs identified (FA), random areas outside of FAs (Non-FA), and areas where FAs were distributed randomly on the plasma membrane (Random); box plots showing the relative density of p110α-mNeonGreen molecules recruited to the different regions (box: median with the 25th and 75th percentiles; whiskers: 1.5-fold the interquartile range); scatter plots showing the relationship between the area/intensity of mCherry-FAK and the recruitment density of p110α-mNeonGreen on each FA. (C) Individual p110α-mNeonGreen molecules in the time series from (B) were detected and tracked. Top left: The initiation sites of each track (duration ≥ 3 frames) in FA and Non-FA regions. Top right: Tracks in FA and Non-FA regions. From left to right in the bottom panels: The density of tracks appearing in FA and Non-FA regions; distributions of lifetimes of tracks; distributions of diffusion coefficients (*D*) of tracks; fraction of tracks with different diffusion status. The last 301 frames of each time series were used for tracking analysis (n = 15 cells). (D) Gene-edited mEGFP-p85α^+/+^ cells transiently expressing mCherry-FAK were imaged at 0.2-s intervals for 601 frames. Top and middle: Images of a single frame and maximum-intensity projection of a representative time series. Bottom: Box plots showing the mean intensity of mEGFP-p85α recruited to FA, Random, and Non-FA regions; scatter plots showing the relationship between the intensity of mCherry-FAK and the integrated intensity of mEGFP-p85α on each FA in the time series. (E) Dual gene-edited p110α-mNeonGreen^+/-^ and Halo-p85α^+/+^ cells transiently expressing mCherry-FAK were imaged at 1-s intervals for 301 frames. Top and middle: Images of a single frame and maximum-intensity projection of a representative time series. Bottom: Montage showing the indicated frames of the enlarged boxed regions. (F and G) p110α-mNeonGreen^+/-^ cells (F) or mEGFP-p85α^+/+^ cells (G) transiently expressing mCherry-FAK were cultured overnight in complete medium (Fed) or starvation medium (Starved), and then imaged at 0.2-s intervals for 601 frames. Left: Images of the maximum-intensity projection of a representative time series. Right: Plots showing the density of p110α-mNeonGreen spots or the mean intensity of mEGFP-p85α recruited to FAs (mean ± SEM; n = 9 Fed and 10 Starved cells in F; n = 8 Fed and 13 Starved cells in G). Cells were imaged at the bottom surface by TIRF microscopy in (A-G). Statistical analysis was performed using the ordinary one-way ANOVA with Tukey’s multiple comparisons test; \*\*\*\**P* < 0.0001; ns, not significant. Scale bars, 10 μm.

This unexpected observation prompted us to explore the precise subcellular locations of these structured patterns. FAs are specialized plasma membrane-associated large macromolecular assemblies that mediate cell-extracellular matrix (ECM) interactions (Case et al., 2015; Geiger et al., 2009). FAs serve as a signaling hub to sense and transduce extracellular biochemical and physical cues to regulate cell adhesion and migration (Case et al., 2015; Kuo, 2014). Given the resemblance between the patterns created by p110α molecules and those of FAs, we transiently expressed the FA marker mCherry-FAK (focal adhesion kinase) in p110α-mNeonGreen^+/-^ cells and conducted single-molecule imaging. Remarkably, individual p110α molecules constantly and transiently visited FAs, with the clustered structures formed by p110α molecules showing nice colocalization with FAK-labeled FAs (Figure 1B; Video S2). This spatial and temporal correlation was further confirmed by imaging p110α-mNeonGreen^+/-^ cells transiently expressing two other FA markers (Figure S1B). By comparing the membrane areas occupied by FAs with other random or non-FA areas, we further confirmed the preferential recruitment of p110α molecules to FAs (Figure 1B). The cumulative number of p110α molecules recruited over a given period was proportional to the size of the FA structure, while the relative frequency was largely unrelated (Figure 1B). These results suggest that p110α molecules are primarily recruited to FAs, and larger FA structures tend to recruit more p110α molecules due to their larger membrane areas.

Additionally, by performing single-molecule tracking and analysis of individual p110α molecules, we found that the frequency of tracks appearing in FAs was more than ten times higher than in non-FA regions (Figure 1C). While p110α molecules displayed short residence times within both FA and non-FA regions, the tracks appearing in FAs predominantly exhibited confined or immobile behavior (Figure 1C). This suggests that p110α binds to relatively stationary partners within FAs. For further validation of the preferential recruitment of other class I PI3K enzymes to FAs, we generated a gene-edited cell line by fusing mNeonGreen to endogenous p110β (p110β-mNeonGreen^+/+^) (Figure S1C). Similar to p110α, the endogenous p110β was preferentially and dynamically recruited to FAs (Figure S1D). Furthermore, when we transiently expressed mNeonGreen-tagged p110α, p110β, p110δ, or p110γ in SUM159 cells at expression levels similar to those of endogenous p110α and p110β, we found that, in addition to p110α and p110β, the class IA catalytic subunit p110δ also exhibited dynamic recruitment to FAs (Figure S1E). However, the class IB catalytic subunit p110γ did not display such behavior (Figure S1E). The preferential recruitment of p110α and p110β to FAs was further verified in multiple other cell lines (Figures S3A-S3D). Thus, it appears that FAs are the preferential and primary membrane binding sites for the catalytic subunits of class IA PI3K in mammalian cells under basal conditions.

Structural and biochemical studies have revealed that the catalytic and regulatory subunits of class I PI3K form a complex (Aytenfisu et al., 2022; Huang et al., 2007; Vadas et al., 2011). We found that the removal of p110α’s N-terminal adaptor binding domain, which interacts with the iSH2 domain of p85 (Huang et al., 2007), prevented the recruitment of p110α to the plasma membrane (Figure S1F). This indicates that p85 is required for the effective recruitment of p110α to FAs at the plasma membrane. To explore the subcellular localization of the endogenous regulatory p85 subunit, we generated a gene-edited cell line expressing mEGFP-p85α^+/+^ (Figure S2A). Live-cell imaging revealed that mEGFP-p85α^+/+^ was specifically recruited and highly concentrated in FAs (Figures 1D and S2B; Video S3). The transiently expressed p85α displayed similar preferential enrichment in FAs in several other cell lines (Figures S3A-S3D). Furthermore, the endogenous regulatory p85β subunit in the gene-edited cells expressing mEGFP-p85β^+/+^ also exhibited preferential recruitment to FAs (Figures S2C and S2D).

To investigate the subcellular localization and relationship of the endogenous catalytic and regulatory subunits in live cells, we generated a gene-edited cell line expressing both p110α-mNeonGreen^+/-^ and Halo-p85α^+/+^ (Figure S2E). Both p110α and p85α molecules exhibited preferential recruitment to FAs, with p85α molecules showing a more stable association and greater enrichment with FAs (Figure 1E). The simultaneous co-appearance of p110α and p85α molecules was also observed (Figure 1E), indicating the membrane recruitment of the p110α-p85α complex from the cytosol to FAs. Furthermore, we noticed that overnight starvation of the cells did not affect the dynamic recruitment of endogenous p110α and p85α molecules to FAs at the plasma membrane (Figures 1F and 1G).

### Dynamic compartmentalized enrichment of PI(3,4,5)P_3_ at the plasma membrane

Activated class I PI3K catalyzes the generation of PI(3,4,5)P_3_. The preferential spatial targeting of class IA PI3K to FAs prompted us to speculate whether PI(3,4,5)P_3_ is spatially generated in FAs in cells under basal conditions. In previous studies, the PH-TH module of Btk has been used to monitor PI(3,4,5)P_3_ increases in cells stimulated by agonists such as EGF, PDGF, or insulin (Manna et al., 2007; Varnai et al., 1999). Nevertheless, due to the extremely low levels of PI(3,4,5)P_3_ under basal conditions and the limited sensitivity of the single PH-TH module, detecting the subcellular distribution and dynamics of trace amounts of PI(3,4,5)P_3_ in mammalian cells has been technically challenging. It was reported recently that the lipid-binding PH-TH module of Btk operates as a dimer, and the PH-TH dimer is ultrasensitive to the density of PI(3,4,5)P_3_ at the membrane surface of supported lipid bilayers (Chung et al., 2019; Wang et al., 2019). Therefore, to enhance the sensitivity of the commonly used PH-TH module for PI(3,4,5)P_3_ detection in cells, we generated a tandem dimer of the PH-TH module of human Btk (residues 2-177) (Varnai et al., 1999) fused with a fluorescent protein, similar to a recently reported approach (Walpole et al., 2022). Interestingly, in comparison to the mEGFP-tagged single PH-TH module, which exhibited some weak local enrichment but mostly diffuse distribution, the mEGFP-tagged tandem dimer displayed a distinct and uneven distribution at the plasma membrane (Figure S4A). Specifically, the tandem dimer exhibited regional enrichment clustered around the cell periphery, whereas the PI(4,5)P_2_ sensor in the same cell showed nearly uniform distribution (Figure S4A).

To reduce the size and minimize the possibility of non-specific binding (Okoh and Vihinen, 1999), we trimmed the C-terminus of the PH-TH module and determined that the minimal essential residues required for effective tandem dimer recruitment were 2-166 (Figure S4B). Consistent with the original tandem dimer (residues 2-177), the trimmed tandem dimer (residues 2-166) displayed a similar uneven regional enrichment around the cell periphery (Figure 2A). Point mutations interfering with PI(3,4,5)P_3_ binding (R28C) or dimer formation (F98V) (Wang et al., 2019) prevented recruitment of mEGFP-2xBtk(2-166) to the plasma membrane (Figures 2B and 2D). Acute depletion or generation of PI(3,4,5)P_3_ at the plasma membrane using rapamycin-induced recruitment of the inositol 3-phosphatase PTEN or the iSH2 domain of p85α with the FKBP–FRB system resulted in the rapid dissociation or recruitment, respectively, of mEGFP-2xBtk(2-166) (Figures S4C and S4D). Similarly, inhibiting PI(3,4,5)P_3_ generation with alpelisib (a selective inhibitor of p110α used in clinical practice) or inducing PI(3,4,5)P_3_ generation with EGF also caused the rapid dissociation or recruitment of mEGFP-2xBtk(2-166) (Figures S4E and S4F). Consistent with the notion that the plasma membrane PI(3,4)P_2_ is primarily the product of PI(3,4,5)P_3_ (Goulden et al., 2019), the PI(3,4)P_2_ sensor exhibited partial regional overlap with the 2xBtk(2-166) sensor (Figure S4H). Acute depletion of PI(3,4)P_2_ through the recruitment of the PI(3,4)P_2_-specific phosphatase INPP4B reduced the binding of the PI(3,4)P_2_ sensor at the plasma membrane (Goulden et al., 2019), while the regional enrichment of mEGFP-2xBtk(2-166) remained unaffected (Figure S4I). Notably, compared to the monomeric form, the dimeric PH-TH module exhibited more pronounced local enrichment (Figures 2A-2D and S4G) and displayed a much stronger dynamic response to PI(3,4,5)P_3_ elimination or induction (Figures S4E and S4F), highlighting a higher avidity of 2xBtk(2-166) for PI(3,4,5)P_3_. Together, the above results demonstrate that 2xBtk(2-166) fused with a fluorescent tag serves as a highly selective and sensitive genetically encoded biosensor for PI(3,4,5)P_3_, hereinafter referred to as the PI(3,4,5)P_3_ sensor.

**Figure 2.**
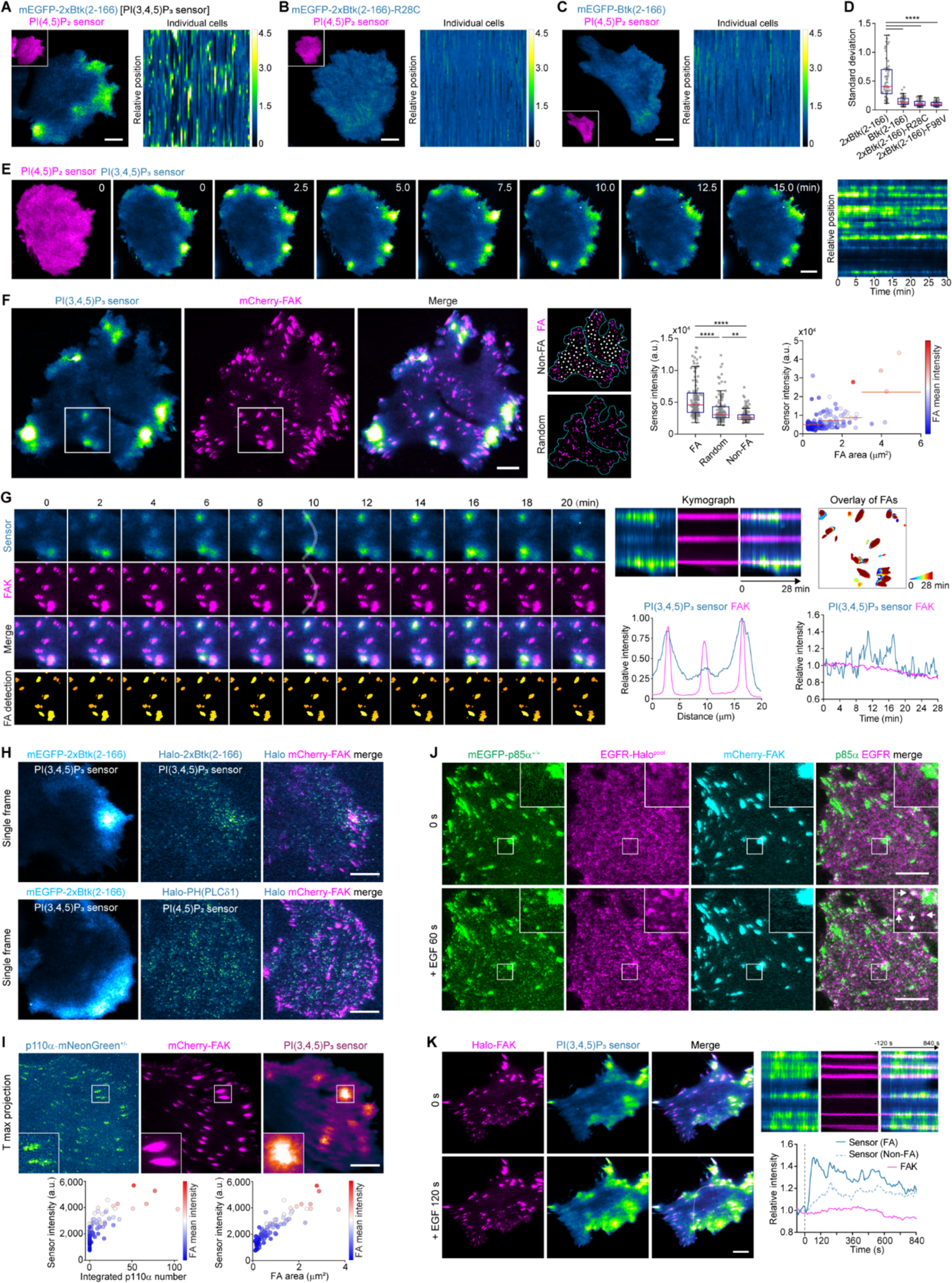
PI(3,4,5)P_3_ is dynamically generated and enriched around FAs. (A-C) Cells transiently expressing the PI(4,5)P_2_ sensor mScarlet-I-PH(PLC81) along with low levels of mEGFP-tagged 2xBtk(2-166) (A), 2xBtk(2-166)-R28C (B), or Btk(2-166) (C) were imaged at the bottom surfaces by TIRF microscopy. Left: Representative images of the mEGFP-tagged Btk sensor and the PI(4,5)P_2_ sensor (inserts). Right: The intensity profile heatmap of the mEGFP-tagged Btk sensor around the cell periphery from multiple cells (n = 60 cells each). (D) Box plots showing the standard deviation of mEGFP-tagged sensor distribution at the plasma membrane (box: median with the 25th and 75th percentiles; whiskers: 1.5-fold the interquartile range; n = 62, 62, 63, and 60 cells). (E) Cells transiently expressing the PI(3,4,5)P_3_ sensor mEGFP-2xBtk(2-166) and the PI(4,5)P_2_ sensor mScarlet-I-PH(PLC81) were imaged at 10-s intervals for 30 min. Representative montage shows PI(3,4,5)P_3_ and PI(4,5)P_2_ sensors at the indicated times. The fluorescence intensity profile of the PI(3,4,5)P_3_ sensor around the cell periphery over time is shown in the right panel. (F) Cells transiently expressing the PI(3,4,5)P_3_ sensor mEGFP-2xBtk(2-166) and mCherry-FAK were imaged at the bottom surfaces at 10-s intervals by TIRF microscopy. Left: Images of PI(3,4,5)P_3_ sensor and mCherry-FAK from a single frame of a representative time series. Middle: Images showing FA, Non-FA, and Random regions of the cells. Right: Box plots showing the mean fluorescence intensity of the PI(3,4,5)P_3_ sensor on each FA, Non-FA, or Random regions; scatter plots showing the relationship between the fluorescence intensity of the PI(3,4,5)P_3_ sensor and the size/intensity of each FA (red lines represent the average intensity within the indicated area ranges). (G) The montage shows selected frames of the boxed regions in (F) (FAs detected are plotted at the bottom). Kymographs were generated along the white line (on the 10-min frame) for the time series. The plots below the kymographs show the relative intensity profile along the line on the single frame. Top right: FAs detected in the time series are overlaid, with each frame represented by a different color. Bottom right: Plots showing the relative intensity of the PI(3,4,5)P_3_ sensor and mCherry-FAK on FAs over time. (H) Top: Images of a cell transiently expressing the PI(3,4,5)P_3_ sensor Halo-2xBtk(2-166) (labeled with JFX_650_-HaloTag ligand), mEGFP-2xBtk(2-166), and mCherry-FAK. Bottom: Images of a cell transiently expressing the PI(4,5)P_2_ sensor Halo-PH(PLC81) (labeled with JFX_650_-HaloTag ligand), the PI(3,4,5)P_3_ sensor mEGFP-2xBtk(2-166), and mCherry-FAK. (I) Top: Images of the maximum-intensity projection of a representative time series. Bottom: Scatter plots showing the relationship between the intensity of the PI(3,4,5)P_3_ sensor and the integrated number of p110α-mNeonGreen molecules (left) or the area/intensity of mCherry-FAK (right) on each FA in the time series. (J) Dual gene-edited mEGFP-p85α^+/+^ and EGFR-Halo (pool, labeled with JFX_650_-HaloTag ligand) cells transiently expressing mCherry-FAK were imaged at 10-s intervals, with EGF added (set as 0 s) during continuous imaging. Representative images show the recruitment of mEGFP-p85α to the plasma membrane and colocalization with EGFR (arrows) after EGF treatment. (K) Cells transiently expressing the PI(3,4,5)P_3_ sensor mEGFP-2xBtk(2-166) and Halo-FAK (labeled with JFX_650_-HaloTag ligand) were imaged at 10-s intervals with EGF added (set as 0 s) during continuous imaging. Images from a representative time series at 0 s and 120 s after EGF treatment are shown. Kymographs were generated along the white line for the time series. Plots show the relative intensity of the PI(3,4,5)P_3_ sensor at FA and Non-FA regions, as well as Halo-FAK at FAs over time. Cells were imaged at the bottom surface by TIRF microscopy in (A-K). Statistical analysis in (D) and (F) was performed using ordinary one-way ANOVA with Tukey’s multiple comparisons test; \*\**P <* 0.001; \*\*\*\**P* < 0.0001. Scale bars, 10 μm.

With this novel PI(3,4,5)P_3_ sensor, we observed dynamic and active oscillations of PI(3,4,5)P_3_ at multiple subcellular regions of the plasma membrane in SUM159 cells through live-cell TIRF imaging (Figure 2E; Video S4). By imaging the cells in 3D using spinning-disk confocal microscopy, we further verified that the PI(3,4,5)P_3_ sensor was highly enriched at several local regions on the basal plasma membrane as well as the leading edge in the middle plane of the cells (Figure S4J). Similar to SUM159 cells (with a heterozygous *PIK3CA^H1047L^* mutation), breast cancer T47D cells (with a heterozygous *PIK3CA^H1047R^* mutation) also displayed strong uneven regional enrichment and active oscillation of the PI(3,4,5)P_3_ sensor at the plasma membrane (Figure S5A; Video S5). Other cancer cell lines such as HCT116 (with a heterozygous *PIK3CA^H1047R^* mutation) and HeLa also exhibited local enrichment and oscillation of the PI(3,4,5)P_3_ sensor at the plasma membrane in a subset of cells (Figure S5A; Videos S6 and S7). Cells that exhibited active random migration during the imaging process, such as the fibrosarcoma cell line HT-1080, tended to recruit the PI(3,4,5)P_3_ sensor to the growing edge of the migrating cells (Figure S5A and Video S8).

### PI(3,4,5)P_3_ is generated and enriched around FAs

To explore whether the subcellular regions with local enrichment of the PI(3,4,5)P_3_ sensor are related to FAs, we imaged cells co-expressing the PI(3,4,5)P_3_ sensor and mCherry-FAK. Excitingly, we noted a close spatial and temporal correlation between the locally enriched PI(3,4,5)P_3_ sensor and FAK-marked FAs in SUM159 cells (Figures 2F and 2G; Video S9), T47D cells (Figure S5B; Video S10), and several other cell lines (Figure S5C). This spatial correlation was further verified in cells transiently expressing other makers for FAs (Figure S6A), or in gene-edited cells with fluorescently tagged endogenous FAK or paxillin (Figure S6B). At the single-molecule level, JFX dye-labeled individual PI(3,4,5)P_3_ sensor molecules diffused and enriched in regions with clusters of FAs (Figure 2H). Conversely, JFX dye-labeled individual PI(4,5)P_2_ sensor molecules exhibited non-selective diffusion throughout the plasma membrane (Figure 2H).

Moreover, we noted that local generation, enrichment, and fluctuation of the PI(3,4,5)P_3_ sensor signal were closely related to different types of adhesion structures at the plasma membrane. In the case of relatively stationary cells during imaging (e.g., SUM159 and T47D cells), the locally enriched PI(3,4,5)P_3_ sensor signals primarily associated with larger or clusters of stable FAs (Figures 2F and S5B; Videos S9 and S10). On a subset of these relatively stationary FAs, the enriched PI(3,4,5)P_3_ sensor exhibited active local fluctuations (Figures 2G and S5B). In cells forming active lamellipodia or membrane protrusions during their migration, the PI(3,4,5)P_3_ sensor signal was highly enriched within nascent adhesions located at the growing membrane protrusions (Figure S6C; Video S11). The generation and clearance of the PI(3,4,5)P_3_ sensor signal closely correlated with the extension and retraction of surface protrusions. Inhibition of PI(3,4,5)P_3_ generation resulted in the disassembly of nascent adhesions, retraction of lamellipodial protrusions, and impaired formation of new nascent adhesions and surface protrusions (Figure S6D). It has been observed that PI(3,4,5)P_3_ (detected by the PH domain of AKT) can propagate with F-actin waves on the basal surface of human cancer cells, thus forming the excitable Ras/PI3K/ERK network (Zhan et al., 2020). By imaging the basal surface of cells co-expressing the PI(3,4,5)P_3_ sensor, F-actin marker, and Halo-FAK, we showed that the PI(3,4,5)P_3_ wave propagated concurrently with the FAK wave and slightly lagged behind the F-actin wave on the cell surface (Figure S6E; Video S12). Thus, in all these scenarios, the PI(3,4,5)P_3_ sensor signal is highly correlated with distinct adhesion structures at the plasma membrane.

To further analyze the spatiotemporal relationship between the recruitment of class IA PI3K and PI(3,4,5)P_3_ generation, we monitored the real-time recruitment of endogenous p110α and the PI(3,4,5)P_3_ sensor to individual FAs using triple-color TIRF microscopy. Intriguingly, while nearly all FAs displayed recruitment of p110α, PI(3,4,5)P_3_ was preferentially enriched in a subset of FAs (Figure 2I). Typically, these PI(3,4,5)P_3_ sensor-enriched FAs exhibited larger areas or clustered together, subsequently recruiting more catalytic class IA PI3K enzymes to catalyze local PI(3,4,5)P_3_ generation (Figure 2I).

### Recruitment of class I PI3K and generation of PI(3,4,5)P_3_ in EGF-stimulated cells

Consistent with the observation that the recruitment of class IA PI3K to FAs was not affected by serum depletion (Figures 1F and 1G), the compartmentalized distribution of the PI(3,4,5)P_3_ sensor at the plasma membrane was similarly observed in cells subjected to overnight starvation (Figure S6F). Since the prevailing view suggests that the membrane recruitment of class I PI3K is mediated by agonist-stimulated RTKs or GPCRs (Aytenfisu et al., 2022; Backer, 2010; Balla, 2013; Batrouni and Baskin, 2020; Bilanges et al., 2019; Burke, 2018; Burke and Williams, 2015; Fruman et al., 2017; Salamon and Backer, 2013; Vadas et al., 2011), we then speculated whether the dynamics and mechanisms of class IA PI3K recruitment and activation differ between cells exposed to agonist stimulation and those under basal conditions.

To monitor the recruitment dynamics of p110α and p85α molecules in live cells during agonist stimulation, we generated gene-edited cell lines in which the endogenous epidermal growth factor receptor (EGFR) of the RTK family was fused with the HaloTag. Within one minute of EGF stimulation, both p85α and p110α molecules were rapidly and dynamically recruited to the plasma membrane (Figures 2J and S2F). Notably, these EGF-induced newly recruited p85α and p110α molecules localized predominantly outside of FAs and exhibited colocalization with clustered EGFR spots on the cell membrane (Figures 2J and S2F). Interestingly, in FA regions, EGF stimulation caused a brief dissociation (∼30-60 s), followed by the re-accumulation of p85α (Figure S2G). This is consistent with the continuous PI(3,4,5)P_3_ generation in FAs and the transient PI(3,4,5)P_3_ generation across the rest of the plasma membrane by activated EGFR (Figure 2K). Together, these findings confirm that the mechanisms governing the spatial targeting and activation of class IA PI3K indeed vary significantly between cells under basal and stimulated conditions. p110α and p85α are primarily recruited to FAs for activation under basal conditions. However, with external agonist stimulation, p110α and p85α are transiently recruited to the activated receptors, in addition to FAs, for activation.

### AKT1 is dynamically recruited around FAs for activation

PI(3,4,5)P_3_ recruits and activates effector proteins such as the PH domain-containing protein kinase AKT (Ebner et al., 2017a; Riehle et al., 2013). Thus, the compartmented enrichment of PI(3,4,5)P_3_ around FAs prompted speculation about whether AKT recruitment and activation are also organized around FAs. Studies over the years have shown that in response to growth factor stimulation, AKT is recruited from the cytosol to the plasma membrane for activation (Ebner et al., 2017b; Liu et al., 2018; Norris et al., 2017; Watton and Downward, 1999). However, it remains unclear whether FAs serves as the subcellular compartment organizing the dynamic recruitment and activation of AKT. To explore the subcellular distribution and more importantly, the recruitment dynamics of AKT in live cells, we created genome-edited SUM159 cells in which endogenous AKT1 was tagged with mNeonGreen at the N-terminus (Figure S7A). By single-molecule imaging, we noticed that endogenous AKT1 molecules showed dynamic recruitment and uneven distribution at the plasma membrane (Figure 3A; Video S13). By creating a maximum-intensity projection of these AKT molecules over a time series, we observed that the well-structured AKT1 patterns were spatially close to FAs (Figure 3A). Individual AKT1 molecules were mainly detected in membrane regions close to FAs (Figure 3B). By tracking the dynamic movement of individual AKT1 molecules, it was evident that AKT1 molecules were preferentially recruited to and concentrated around FAs, as shown by the overlay of tracks (Figure 3C) and track density map (Figure 3D). Interestingly, of these AKT1 molecules recruited dynamically to the plasma membrane, the AKT1 molecules recruited close to FAs showed longer residence time than those recruited to non-FA areas (Figure 3C). AKT1 molecules exhibited rapid diffusion at the plasma membrane (diffusion coefficients of ∼0.3-0.4 μm^2^/s), with those recruited around FAs diffusing slightly slower than those not in proximity to FAs (Figure 3D). Thus, by tracking the single-molecule dynamics of endogenous AKT1, we uncovered that AKT1 molecules are recruited preferentially around FAs at the plasma membrane.

**Figure 3.**
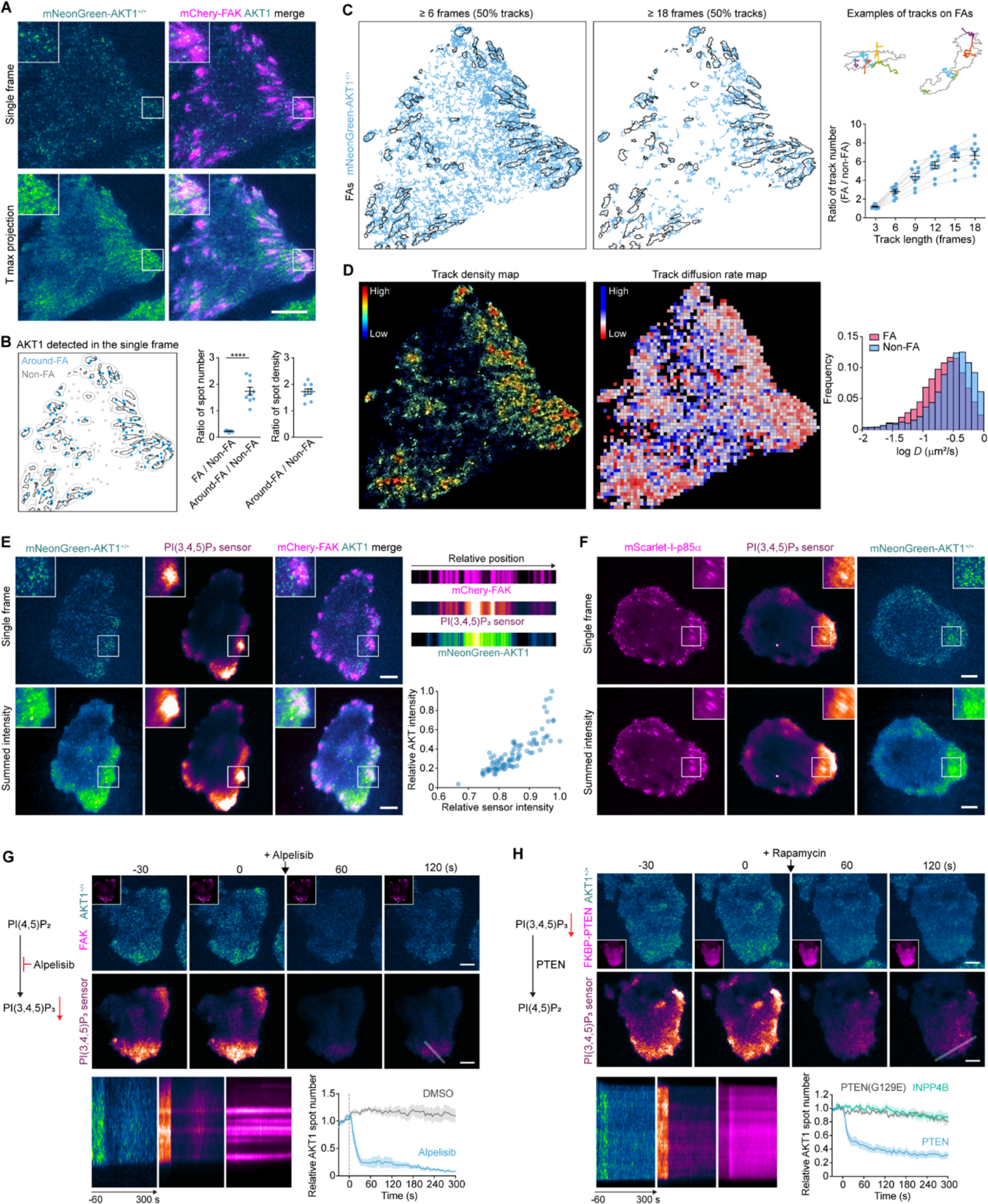
Compartmentalized recruitment of AKT1 around FAs by PI(3,4,5)P_3_. (A) Bottom surfaces of gene-edited mNeonGreen-AKT1^+/+^ cells transiently expressing mCherry-FAK were firstly imaged at 0.1-s intervals for 10 frames, and then imaged at 0.05-s intervals in the mNeonGreen-AKT1^+/+^ channel for 601 frames by TIRF microscopy. Images show a single frame and maximum-intensity projection of a representative time series. (B) AKT1 molecules detected in FAs, areas around FAs (Around-FA), and other regions of the single frame in (A). The plots in the middle show the ratio of AKT1 molecules detected in FA and Non-FA regions, as well as molecules detected in Around-FA and Non-FA regions. The plot on the right shows the ratio of spot densities in Around-FA and Non-FA regions (n = 9 cells). (C) Individual mNeonGreen-AKT1 molecules in the time series from (A) were tracked, and tracks with lifetime ≥ 6 frames or 18 frames were plotted (50% of random tracks are shown). Top right: Enlarged views of two regions with example tracks in different colors. Bottom right: Plots showing the ratio of track number in FAs and Non-FA regions for tracks with lifetime ≥ 3, 6, 9, 12, 15 and 18 frames (n = 9 cells). (D) Left and middle: Track spatial density map (left) and track diffusion map (middle) of mNeonGreen-AKT1 molecules in the time series from (A). Right: Distributions of diffusion coefficients (*D*) of tracks in FA and Non-FA regions (n = 9 cells). (E) mNeonGreen-AKT1^+/+^ cells transiently expressing mCherry-FAK and the PI(3,4,5)P_3_ sensor Halo-2xBtk(2-166) (labeled with JFX_650_-HaloTag ligand) were imaged at 1-s intervals. Left: Images of a single frame and summed intensity of a representative time series (121 frames). Top right: Intensity profiles of the three channels along the cell boundary (40 pixels width, starting from the top of the cell). Bottom right: Scatter plots showing relative intensity of mNeonGreen-AKT1 and PI(3,4,5)P_3_ sensor on each FA of the cell (89 FAs). (F) mNeonGreen-AKT1^+/+^ cells transiently expressing mScarlet-I-p85α and the PI(3,4,5)P_3_ sensor Halo-2xBtk(2-166) were imaged at 1-s intervals. Shown are images of a single frame and summed intensity of a representative time series (121 frames). (G) mNeonGreen-AKT1^+/+^ cells transiently expressing mCherry-FAK and the PI(3,4,5)P_3_ sensor Halo-2xBtk(2-166) were imaged at 5-s intervals. p110α inhibitor alpelisib was added (set as 0 s) during continuous imaging. Top: Montages showing mNeonGreen-AKT1, mCherry-FAK (inserts) and PI(3,4,5)P_3_ sensor before and after alpelisib treatment. Bottom: Kymographs generated along the line showing the membrane recruitment of mNeonGreen-AKT1 and Halo-2xBtk(2-166) before and after alpelisib treatment. Plots show the relative number of mNeonGreen-AKT spots at the plasma membrane before and after DSMO or alpelisib treatment (n = 7 and 6 cells). (H) mNeonGreen-AKT1^+/+^ cells transiently co-expressing the PI(3,4,5)P_3_ sensor Halo-2xBtk(2-166) and LYN11-FRB-ECFP, together with either mCherry-FKBP-PTEN, mCherry-FKBP-PTEN(G129E) or mCherry-FKBP-INPP4B, were imaged at 5-s intervals. Rapamycin was added (set as 0 s) during continuous imaging. Top: Montages showing the removal of mNeonGreen-AKT1 and PI(3,4,5)P_3_ sensor from the plasma membrane upon recruitment of mCherry-FKBP-PTEN (inserts) from the cytosol to the plasma membrane. Kymographs were generated along the line. Plots show the number of mNeonGreen-AKT spots before and after rapamycin treatment (n = 7, 6, 7 cells for PTEN, PTEN(G129E), and INPP4B). Cells were imaged at the bottom surface by TIRF microscopy in (A-H). \*\*\*\**P* < 0.0001 by two-sided Student’s t test. Data are shown as mean ± SEM in (B), (C), (G) and (H). Scale bars, 10 μm.

Moreover, by monitoring AKT1 recruitment and PI(3,4,5)P_3_ generation in the same cells, we observed the preferential recruitment of AKT1 molecules to these PI(3,4,5)P_3_-enriched FAs (Figure 3E; Video S14). The amount of AKT1 molecules recruited to individual FAs was highly related with the amount of PI(3,4,5)P_3_. Live-cell tracking of p85α subunit recruitment, PI(3,4,5)P_3_ generation, and AKT1 recruitment further showed that although class IA PI3Ks were recruited to almost all FAs, PI(3,4,5)P_3_ and AKT1 molecules were enriched only around a subset of larger FAs or clusters of FAs (Figure 3F). The preferential recruitment of AKT1 molecules to PI(3,4,5)P_3_-enriched FAs was also observed in several different cell lines (Figure S7B).

To further demonstrate that the compartmentalized AKT recruitment around FAs is indeed mediated by class I PI3K, we treated cells with the PI3K inhibitor alpelisib during continuous imaging. Alpelisib treatment led to the immediate dissociation of the PI(3,4,5)P_3_ sensor from the plasma membrane, accompanied by the reduced membrane recruitment of AKT1 (Figure 3G). Given that the PH domain of AKT1 can bind to both PI(3,4,5)P_3_ and PI(3,4)P_2_ *in vitro* (Franke et al., 1997; Liu et al., 2018), we then determined whether the dynamic membrane recruitment of ATK1 is indeed mediated by PI(3,4,5)P_3_. By recruiting the inositol 3-phosphatase PTEN or the PI(3,4)P_2_-specific phosphatase INPP4B to the plasma membrane, we found that acute depletion of PI(3,4)P_2_ by INPP4B effectively reduced the membrane binding of the PI(3,4)P_2_ sensor but did not affect the membrane recruitment of AKT1 (Figures 3H and S7C). However, PTEN-mediated PI(3,4,5)P_3_ depletion led to the immediate disappearance of both the PI(3,4,5)P_3_ sensor and AKT1 from the plasma membrane (Figures 3H and S7D). Furthermore, activation of class I PI3K by EGF led to the transient generation and enrichment of PI(3,4,5)P_3_ mostly around clusters of FAs at the plasma membrane, which led to the increased membrane recruitment of AKT1 molecules (Figure S7E). Together, the above results demonstrate that the preferential recruitment of class IA PI3Ks to FAs results in the localized generation of PI(3,4,5)P_3_, which further mediates and determines the compartmentalized recruitment of AKT1 around FAs.

### The recruitment of class IA PI3Ks, generation of PI(3,4,5)P_3_, and recruitment/activation of AKT are regulated by the activated FAK

The next question is how FAs regulate the local recruitment of class IA PI3K and thus the compartmentalized PI(3,4,5)P_3_ generation and AKT activation. The nonreceptor tyrosine kinase FAK is a central signaling mediator of integrin-mediated signaling from FAs (Luo and Guan, 2010; Zhao and Guan, 2011). FAK is activated in response to integrin–ECM engagement during cell attachment and various extracellular stimuli (Luo and Guan, 2010). Interestingly, treating cells with defactinib (also known as VS-6063), a second-generation FAK inhibitor used for anticancer combination therapies (Dawson et al., 2021), effectively reduced the PI(3,4,5)P_3_ sensor signal from the plasma membrane of SUM159 cells (Figures 4A and 4B; Video S15). However, the FA structures remained largely unchanged immediately following defactinib treatment (Figure 4A). The immediate inhibitory impact of defactinib on PI(3,4,5)P_3_ generation was also observed in two additional cell lines (Figures S8A and S8B).

**Figure 4.**
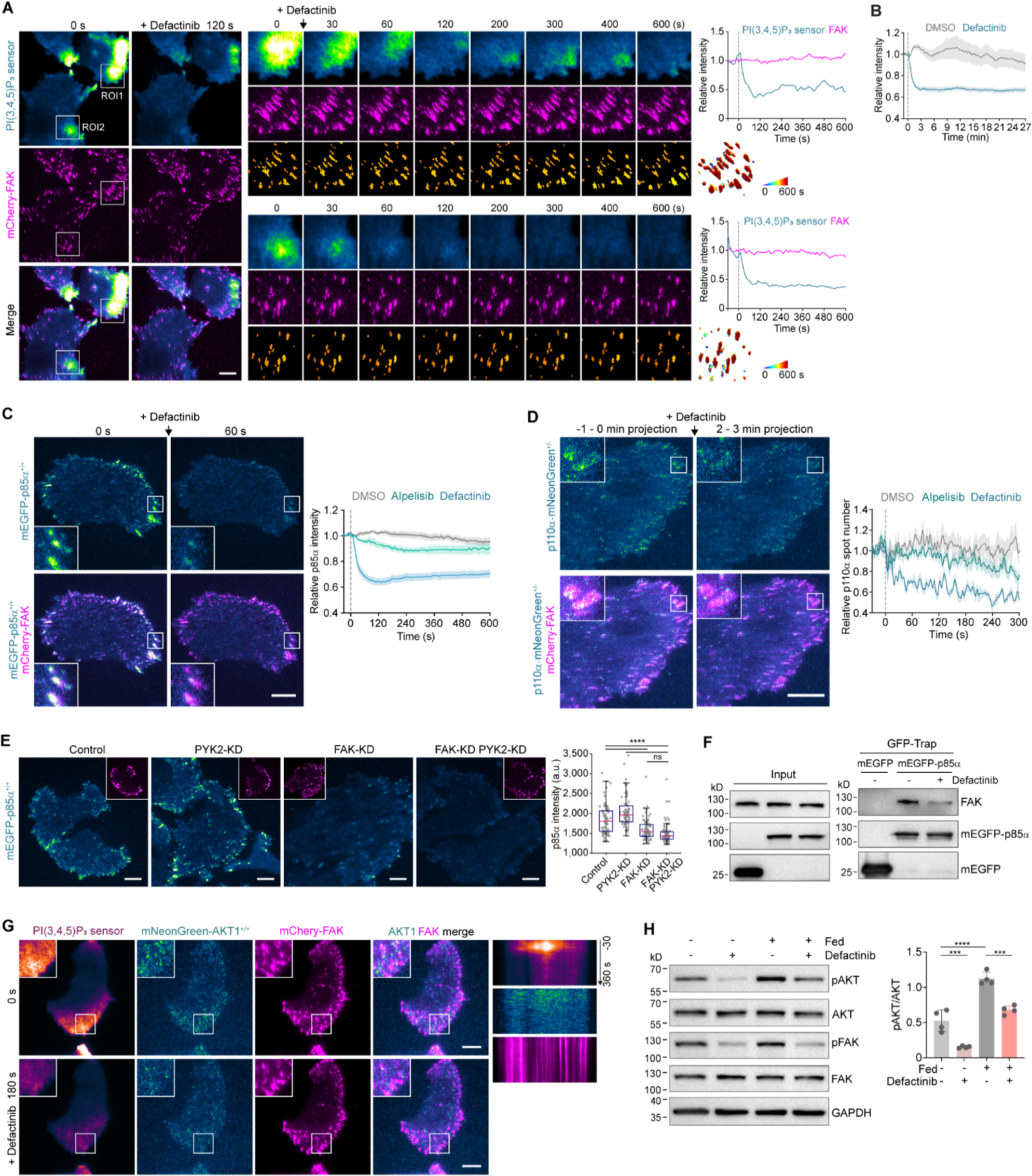
FAK regulates class IA PI3K recruitment, PI(3,4,5)P_3_ generation, and AKT activation. (A) Cells transiently expressing the PI(3,4,5)P_3_ sensor mEGFP-2xBtk(2-166) and mCherry-FAK were imaged at 10-s intervals, with FAK inhibitor defactinib added (set as 0 s) during continuous imaging. Left: Images at 0 s and 120 s after defactinib treatment in a representative time series. Middle: Montages showing the indicated frames of ROI1 and ROI2, with detected FAs plotted at the bottom. Right: Plots showing the relative intensity of PI(3,4,5)P_3_ sensor and mCherry-FAK on FAs in ROI1 and ROI2 over time. (B) Plots showing the average intensity of the PI(3,4,5)P_3_ sensor mEGFP-2xBtk(2-166) at the plasma membrane from cells treated with DMSO or defactinib (n = 13 and 25 cells). (C) mEGFP-p85α^+/+^ cells transiently expressing mCherry-FAK were imaged at 10-s intervals, with defactinib or p110α inhibitor alpelisib added (set as 0 s) during continuous imaging. Left: Images at 0 s and 120 s after defactinib treatment from a representative time series. Right: Plots showing the average intensity of mEGFP-p85α on FAs in cells treated with DMSO, defactinib, or alpelisib (n = 8, 11, and 7 cells). (D) p110α-mNeonGreen^+/-^ cells transiently expressing mCherry-FAK and the PI(3,4,5)P_3_ sensor Halo-2xBtk(2-166) were imaged at 1-s intervals, with DMSO, defactinib, or alpelisib added (set as 0 s) during continuous imaging. Left: Maximum-intensity projections of the 60 frames before and 1 min after defactinib treatment from a representative time series. Right: Plots showing the relative numbers of p110α-mNeonGreen molecules recruited to the plasma membrane of cells treated with DMSO, defactinib, or alpelisib (n = 5, 5, and 6 cells). (E) mEGFP-p85α^+/+^ cells were treated with control shRNA or shRNA targeting FAK, PYK2, or both FAK and PYK2. Left: Images showing the distribution of mEGFP-p85α and transiently expressed mCherry-paxillin (inserts) at the bottom surfaces of the shRNA-treated cells. Right: Box plots showing the average intensity of mEGFP-p85α at the plasma membrane of shRNA-treated cells (box: median with the 25^th^ and 75^th^ percentiles; whiskers: 1.5-fold the interquartile range; n = 66, 69, 65, and 69 cells). (F) Cells transiently expressing mEGFP or mEGFP-p85α were treated with DMSO or defactinib. Compared to the DMSO-treated cells, the level of endogenous FAK co-precipitated by mEGFP-p85α was reduced. (G) mNeonGreen-AKT1^+/+^ cells transiently expressing mCherry-FAK and the PI(3,4,5)P_3_ sensor Halo-2xBtk(2-166) were imaged at 5-s intervals. Defactinib was added (set as 0 s) during continuous imaging. Left: Images at 0 s and 180 s after defactinib treatment from representative time series. Right: Kymographs showing the reduced membrane recruitment of Halo-2xBtk(2-166) and mNeonGreen-AKT1 after defactinib treatment. (H) After culturing in complete medium (Fed) or starvation medium for 24 hours, cells were treated with DMSO or defactinib for 10 min. Total and phosphorylated AKT and FAK were analyzed by western blot with the indicated antibodies. The ratios of phosphorylated AKT (pAKT) to total AKT are shown for four independent experiments. Data are shown as mean ± SEM in (B-D) and mean ± SD in (H). Statistical analysis was performed using ordinary one-way ANOVA with Tukey’s multiple comparisons test; \*\*\**P* < 0.001; \*\*\*\**P* < 0.0001; ns, not significant. Scale bars, 10 μm.

Why does inhibiting FAK activation result in a significant reduction in PI(3,4,5)P_3_ generation? By examining the recruitment dynamics of endogenous p85α at the plasma membrane of cells treated with defactinib, we noticed an immediate dissociation of endogenous p85α from still-intact FAs (Figure 4C). Interestingly, though the PI3K inhibitor alpelisib also led to the immediate decrease of the PI(3,4,5)P_3_ signal, it only slightly reduced the localization of p85α on the plasma membrane (Figure 4C). Furthermore, FAK but not PI3K inhibitor also decreased the recruitment of endogenous p110α to the cell membrane (Figure 4D). In addition to SUM159 cells, the reduced recruitment of p110α and p85α to FAs was also verified in other defactinib-treated cell lines (Figures S8C-S8F). Additionally, defactinib treatment compromised the membrane recruitment of p85β (Figure S8G) and p110β (Figure S8H).

Proline-rich tyrosine kinase 2 (PYK2), another member of the FAK family, is a close paralog of FAK and has redundant roles with FAK under specific circumstances (Dawson et al., 2021). Given that defactinib can also inhibit the kinase activity of PYK2 (Dawson et al., 2021), we then investigated whether PYK2 was also involved in regulating PI(3,4,5)P_3_ generation. We knocked down the expression of either PYK2 or FAK, or both PYK2 and FAK, in gene-edited mEGFP-p85α^+/+^ cells (Figures 4E and S8I). Intriguingly, depleting FAK expression, but not PYK2, significantly reduced the recruitment of endogenous p85α to FAs (Figure 4E). These findings indicate that p85α recruitment to FAs is related primarily to FAK. We further verified the interaction between p85α and FAK, which was effectively reduced by defactinib treatment (Figure 4F). Moreover, treating cells with 14z, a recently developed highly potent FAK inhibitor (Su et al., 2019), also led to the immediate reduction in the recruitment and localization of PI(3,4,5)P_3_ sensor (Figure S9A), p85α (Figure S9B), and p110α (Figure S9C) at the plasma membrane.

The autophosphorylation of FAK at Y397 upon integrin-mediated adhesion triggers the binding of Src, which in turn phosphorylates other tyrosine residues on FAK to induce its full catalytic activation (Lietha et al., 2007). Inhibition of Src reduced the recruitment of p85α and p110α to FAs and attenuated the PI(3,4,5)P_3_ sensor signal at the plasma membrane (Figures S9E and S9F), further demonstrating that the full activation of FAK is crucial for class IA PI3K recruitment and PI(3,4,5)P_3_ generation in FAs. During mitosis, FAs are disassembled and FAK is inactivated as the cells round up and detach from the ECMs (Ma et al., 2001; Yamakita et al., 1999). Through live-cell tracking of the entire mitotic process, we found that PI(3,4,5)P_3_ but not PI(4,5)P_2_ disappeared alongside the disassembly and disappearance of FAs as the cell entered mitosis (Figure S9G). After cytokinesis, the PI(3,4,5)P_3_ sensor signal started to appear and re-enrich in the re-assembled FAs with activated FAK (Figure S9G). Thus, observation of this physiological cellular process further shows that the generation and local enrichment of PI(3,4,5)P_3_ are tightly correlated with FAs and the activated FAK.

In addition to reduced generation of PI(3,4,5)P_3_, inhibition of FAK activity also led to reduced recruitment of AKT1 molecules to the plasma membrane (Figure 4G). In line with this diminished recruitment, the phosphorylation of AKT was significantly attenuated in cells treated with FAK inhibitors (Figures 4H and S9D). Notably, the inhibitory effect on AKT phosphorylation was more pronounced in serum-starved cells (Figures 4H and S9D). Together, these results demonstrate that the recruitment and activation of AKT at the plasma membrane of cells under basal conditions are regulated by the activated FAK.

### Distinct subcellular distributions of PI3K and PTEN determine the spatial distribution of PI(3,4,5)P_3_ and AKT at the plasma membrane

Although class IA PI3Ks were recruited to almost all FA structures, the PI(3,4,5)P_3_ sensor and AKT1 were enriched only in a subset of FAs (Figures 2I and 3E). This discrepancy raises the critical question of how localized PI(3,4,5)P_3_ and AKT enrichment is established and sustained. SUM159 cells carry a heterozygous H1047L mutation in the p110α catalytic subunit (*PIK3CA*^WT/H1047L^). His1047 in the kinase domain of p110α is a mutation hotspot in cancer (Samuels et al., 2004). The oncogenic H1047L or H1047R mutation causes aberrant activation of PI3K-AKT signaling and promotes tumor growth (Vasan et al., 2019). To explore the impact of the oncogenic mutation on the generation and subcellular distribution of PI(3,4,5)P_3_, we generated two gene-edited SUM159 cell lines: one homozygous for H1047L (*PIK3CA*^H1047L/H1047L^) and another homozygous for H1047 (*PIK3CA*^WT/WT^) (Figure 5A). As expected, the phosphorylation level of AKT in the *PIK3CA*^WT/WT^ cells was much lower than that in cells with heterozygous or homozygous H1047L mutation (Figure 5B). By employing mass spectrometry to measure PI(3,4,5)P_3_ levels in whole cells, we verified that the *PIK3CA^H1047L/H1047L^*and *PIK3CA^WT/H1047L^* cells exhibited elevated PI(3,4,5)P_3_ levels compared to the *PIK3CA^WT/WT^* cells (Figure 5C). Interestingly, at the single-cell level, the *PIK3CA*^WT/WT^ cells exhibited a markedly reduced localized enrichment of the PI(3,4,5)P_3_ sensor at the plasma membrane compared to the *PIK3CA*^WT/H1047L^ cells (Figure 5D). The *PIK3CA*^H1047L/H1047L^ cells displayed similar or slightly enhanced localized PI(3,4,5)P_3_ enrichment at the plasma membrane, as observed using TIRF microscopy (Figure 5D) or spinning-disk confocal microscopy in 3D (Figure S10A). By single-molecule imaging, we found that like the wild-type p110α, p110α molecules carrying the H1047L mutation were dynamically and preferentially recruited to FAs in SUM159 cells (Figure 5E) and several other cancer cell lines (Figures S3E-S3G). Therefore, it seems that the catalytic p110α molecules carrying the H1047L oncogenic mutation are still dynamically recruited to FAs, where they promote both the localized production and compartmentalized enrichment of PI(3,4,5)P_3_.

**Figure 5.**
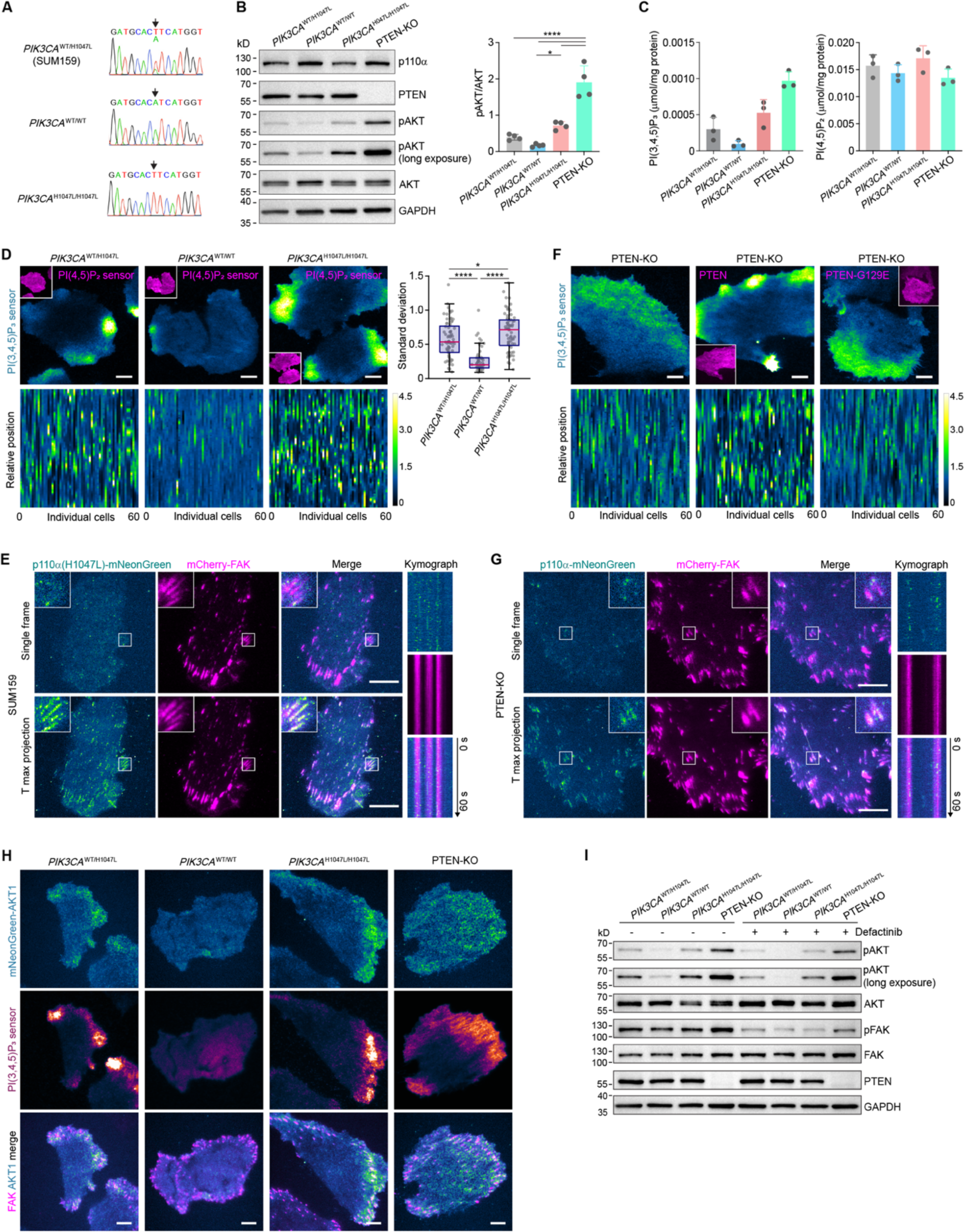
*PIK3CA* and *PTEN* mutation differently regulate the subcellular distributions of PI(3,4,5)P_3_ and AKT1. (A) Sequencing results of genomic DNA from the parental SUM159 cells (*PIK3CA^WT/H1047L^*) and gene-edited SUM159 cells homozygous for H1047 (*PIK3CA^WT/WT^*) or H1047L (*PIK3CA^H1047L/H1047L^*) of *PIK3CA*. (B) *PIK3CA^WT/H1047L^*, *PIK3CA^WT/WT^*, *PIK3CA^H1047L/H1047L^*, or PTEN-KO SUM159 cells were serum-starved overnight and then subjected to western blot analysis using the indicated antibodies. The ratios of phosphorylated AKT to total AKT are shown for four independent experiments. (C) Quantification of PI(3,4,5)P_3_ and PI(4,5)P_2_ levels in the four cell lines by mass spectrometry. (D) *PIK3CA^WT/H1047L^*, *PIK3CA^WT/WT^*, or *PIK3CA^H1047L/H1047L^*cells were transiently transfected with the PI(3,4,5)P_3_ sensor mEGFP-2xBtk(2-166) and the PI(4,5)P_2_ sensor mScarlet-I-PH(PLC81). Shown are representative images and intensity profile heatmaps of the PI(3,4,5)P_3_ sensor in the three cell lines (n = 60 cells each). Box plots show the standard deviation of PI(3,4,5)P_3_ sensor distribution at the plasma membrane (box: median with the 25th and 75th percentiles; whiskers: 1.5-fold the interquartile range; n = 69, 61, and 67 cells). (E) SUM159 cells transiently co-expressing mCherry-FAK with p110α(H1047L)-mNeonGreen were imaged at 0.2-s intervals for 301 frames. Left: Images of a single frame and the maximum-intensity projection of a representative time series are shown. Right: Kymographs generated along the line showing recruitment of p110α(H1047L) to FAs. (F) PTEN-KO SUM159 cells were transiently expressed with the PI(3,4,5)P_3_ sensor mEGFP-2xBtk(2-166), or mEGFP-2xBtk(2-166) together with mScarlet-I-PTEN or mScarlet-I-PTEN-G129E (inserts). Shown are representative images and intensity profile heatmaps of the PI(3,4,5)P_3_ sensor (n = 60 cells each). (G) PTEN-KO SUM159 cells transiently co-expressing mCherry-FAK and p110α-mNeonGreen were imaged at 0.2-s intervals for 301 frames. Left: Images of a single frame and the maximum-intensity projection of a representative time series are shown. Right: Kymographs were generated along the line. (H) *PIK3CA^WT/H1047L^*, *PIK3CA^WT/WT^*, *PIK3CA^H1047L/H1047L^*, or PTEN-KO cells transiently expressing mNeonGreen-AKT1, mCherry-FAK and the PI(3,4,5)P_3_ sensor Halo-2xBtk(2-166) were imaged at 0.3-s intervals. Images of a single frame of a representative time series are shown. (I) *PIK3CA^WT/H1047L^*, *PIK3CA^WT/WT^*, *PIK3CA^H1047L/H1047L^*, or PTEN-KO cells were treated with or without defactinib for 10 min. The total and phosphorylated AKT and FAK levels were analyzed by western blot. Statistical analysis was performed using ordinary one-way ANOVA with Tukey’s multiple comparisons test; \**P* < 0.05; \*\*\*\**P* < 0.0001. Scale bars, 10 μm.

The dephosphorylation of PI(3,4,5)P_3_ can occur through either the 3-phosphatase PTEN or the 5-phosphatase SHIP1/2 (Balla, 2013). As SHIP1 is expressed specifically in hematopoietic cells (Gilby et al., 2007), we generated a SHIP2 knockout (KO) cell line in SUM159 cells (Figure S10B). SHIP2 depletion did not affect the distribution of PI(3,4,5)P_3_ at the plasma membrane (Figure S10C), consistent with SHIP2’s primary localization in clathrin-coated structures at the plasma membrane (Nakatsu et al., 2010). PTEN loss or mutations, frequently observed in breast and lung cancers (Lee et al., 2018), lead to elevated levels of PI(3,4,5)P_3_ in cells (Sun et al., 1999). Interestingly, the PI(3,4,5)P_3_ sensor exhibited a more uniform distribution at the plasma membrane of PTEN-KO SUM159 cells (Figures 5F and S10A). Re-expression of the wild-type PTEN, but not the lipid phosphatase-dead G129E mutant, restored the localized distribution of the PI(3,4,5)P_3_ sensor at the plasma membrane (Figure 5F). Moreover, through mass spectrometry, we confirmed that PTEN knockout strongly increased the cellular level of PI(3,4,5)P_3_ (Figure 5C). Breast cancer SUM149 cells, which have wild-type *PIK3CA* but lack PTEN expression (Hollestelle et al., 2007) (Figure S10D), also exhibited a relatively even distribution of the highly enriched PI(3,4,5)P_3_ sensor at the plasma membrane (Figure S10E). In the PTEN-KO SUM159 cells and PTEN-deficient SUM149 cells, p110α molecules were still dynamically recruited to FAs (Figures 5G and S10F). PTEN was diffusely distributed around the plasma membrane (Figure S10G) and was also found in cytoplasmic vesicles in mammalian cells (Naguib et al., 2015). Thus, while PI(3,4,5)P_3_ is constantly generated by class IA PI3K locally in FAs, PTEN loss results in increased PI(3,4,5)P_3_ levels across the entire cell surface while weakening its compartmentalized distribution.

In accordance with the differential distribution of PI(3,4,5)P_3_ in the *PIK3CA* mutated cells, the dynamic recruitment and compartmentalized distribution of AKT1 molecules around FAs were markedly decreased in SUM159 cells homozygous for H1047 (Figure 5H). Similar to the PI(3,4,5)P_3_ sensor, AKT1 molecules also showed a relatively more uniform distribution at the plasma membrane of PTEN-KO cells (Figure 5H). In the *PIK3CA* or *PTEN* mutated cells, the level of AKT phosphorylation examined by western blotting (Figures 5B and 5I) was quite consistent with the above-described imaging results. Therefore, by manipulating *PIK3CA* and *PTEN* genetically, we found that the spatial generation and compartmentalized distribution of PI(3,4,5)P_3_, and thus AKT recruitment and signaling activation, are collectively controlled by the enzymatic phosphorylation and dephosphorylation metabolic reactions at different subcellular locations. These processes are substantially influenced by oncogenic *PIK3CA* mutations and PTEN loss.

### Combined inhibition of p110α and FAK exerts a stronger inhibitory effect on PI3K-AKT signaling and cancer cell aggressiveness

Given that p110α molecules carrying oncogenic hotspot mutations were also actively recruited to FAs (Figure 5E), we proceeded to explore whether the dynamic recruitment of p110α with oncogenic mutations is also regulated by FAK. Similar to wild-type p110α molecules, the inhibition of FAK but not PI3K effectively diminished the membrane recruitment of p110α molecules with H1047L mutation (Figures 6A). FAK inhibitor treatment also effectively decreased the PI(3,4,5)P_3_ sensor signal at the plasma membrane of SUM159 cells with heterozygous or homozygous *PIK3CA*^H1047L^ mutations (Figure S10H). Therefore, the membrane recruitment and activation of p110α molecules with oncogenic hotspot mutations is also regulated by the activated FAK. Accordingly, FAK inhibition reduced AKT signaling in SUM159 cells with heterozygous or homozygous *PIK3CA*^H1047L^ mutations (Figure 5I).

**Figure 6.**
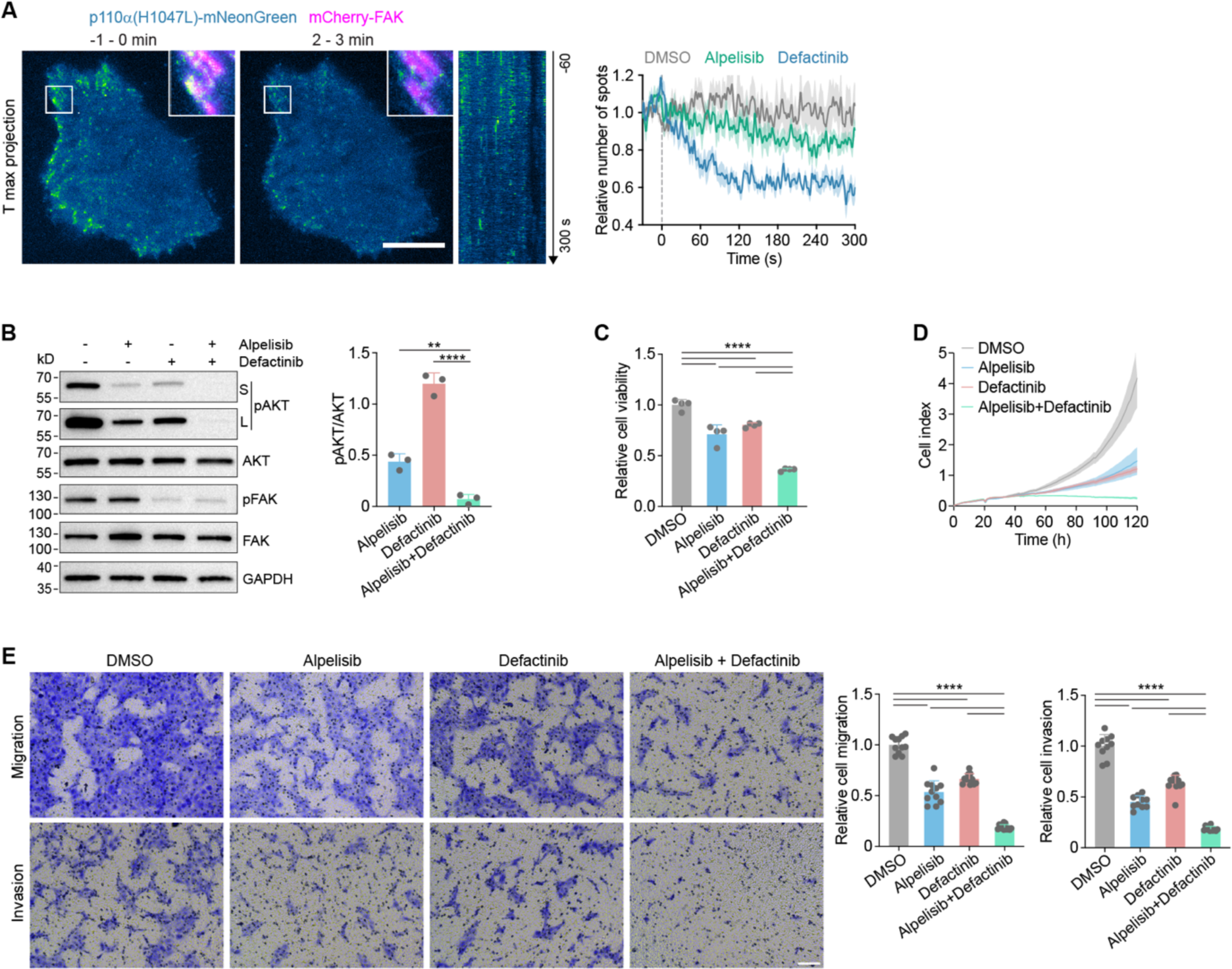
Combined inhibition of p110α and FAK exerts a stronger inhibitory effect on PI3K-AKT signaling and cancer cell aggressiveness. (A) Cells transiently expressing mCherry-FAK with p110α(H1047L)-mNeonGreen were imaged at 1-s intervals, with DMSO, alpelisib, or defactinib added (set as 0 s) during continuous imaging. The maximum-intensity projections of the 60 frames before and 1 minute after defactinib treatment in the time series are shown. Inserts show enlarged boxed regions with the p110α and FAK images overlaid. Plots show the relative numbers of p110α molecules recruited to the plasma membrane (n = 7, 7, and 7 cells). (B) Cells were treated with DMSO, alpelisib, defactinib, or both alpelisib and defactinib for 24 hours. Total and phosphorylated AKT and FAK levels were analyzed by western blot with the indicated antibodies (S: short exposure; L: long exposure). The ratios of phosphorylated AKT to total AKT are shown for three independent experiments. (C) Cells were treated with DMSO, alpelisib, defactinib, or both alpelisib and defactinib for 72 hours, and cell viability was measured by CCK-8 assay. (D) Cells were treated with DMSO, alpelisib, defactinib, or both alpelisib and defactinib, and cell growth curves were generated by real-time cell analysis from one representative experiment. (E) Cells were treated with DMSO, alpelisib, defactinib, or both alpelisib and defactinib for 48 hours, and then subjected to transwell migration and invasion assays. Representative images of cells with different treatments are shown. The migrated and invaded cells were measured in 10-12 random fields per well from one representative experiment. Data are shown as mean ± SEM in (A), mean ± SD in (B-E). Statistical analysis was performed using ordinary one-way ANOVA with Tukey’s multiple comparisons test; \*\**P* < 0.01; \*\*\*\**P* < 0.0001. Scale bars, 10 μm in (A); 100 μm in (E).

FAK inhibitors have been widely used in combination with other agents to enhance efficiency and overcome resistance in the treatments of solid tumors (Dawson et al., 2021). The inhibitory effect of FAK inhibitors on class IA PI3K recruitment/activation, PI(3,4,5)P_3_ generation and AKT recruitment/activation prompted us to explore whether the combination of FAK and PI3K inhibitors, currently used in clinical practice or trials, could synergistically enhance their inhibitory effects on PI3K-AKT signaling and aggressiveness of cancer cells. Encouragingly, compared with the inhibition of p110α or FAK alone, the combined inhibition of FAK and p110α led to a significantly greater reduction in AKT phosphorylation (Figures 6B and S10I). Moreover, the combination of p110α and FAK inhibitors significantly impaired the viability and proliferation (Figures 6C and 6D) and reduced the migration and invasion abilities (Figure 6E) of human breast cancer SUM159 cells. Thus, the combined inhibition of FAK and class I PI3K exerted a stronger inhibitory effect on AKT signaling, as well as on cancer cell growth and aggressiveness.

## DISCUSSION

Contrary to the canonical view that class I PI3K-AKT signaling is mainly activated by agonist-stimulated RTK or GPCRs, our study demonstrates that, under basal conditions, PI3K-AKT signaling activation takes place predominantly in FAs and is regulated by the activated FAK (Figure 7). Mediation of environmental stimuli through ECM-integrin-FAK signaling may represent the primary mechanism underlying the cell-intrinsic regulation of class I PI3K-AKT signaling under physiological conditions. However, in cells treated with ligands for RTKs, the recruitment/activation of class I PI3K is likely dominated by the activated receptors located at the plasma membrane or endosomes (Bilanges et al., 2019; Burke, 2018; Rathinaswamy and Burke, 2020; Vadas et al., 2011). Nevertheless, FAs are still involved in the recruitment and activation of PI3K-AKT signaling in EGF-stimulated cells. Thus, our study suggests that mammalian cells may have evolved different mechanisms for the spatial organization and regulation of class I PI3K-AKT signaling under different environmental biochemical and mechanical cues.

**Figure 7.**
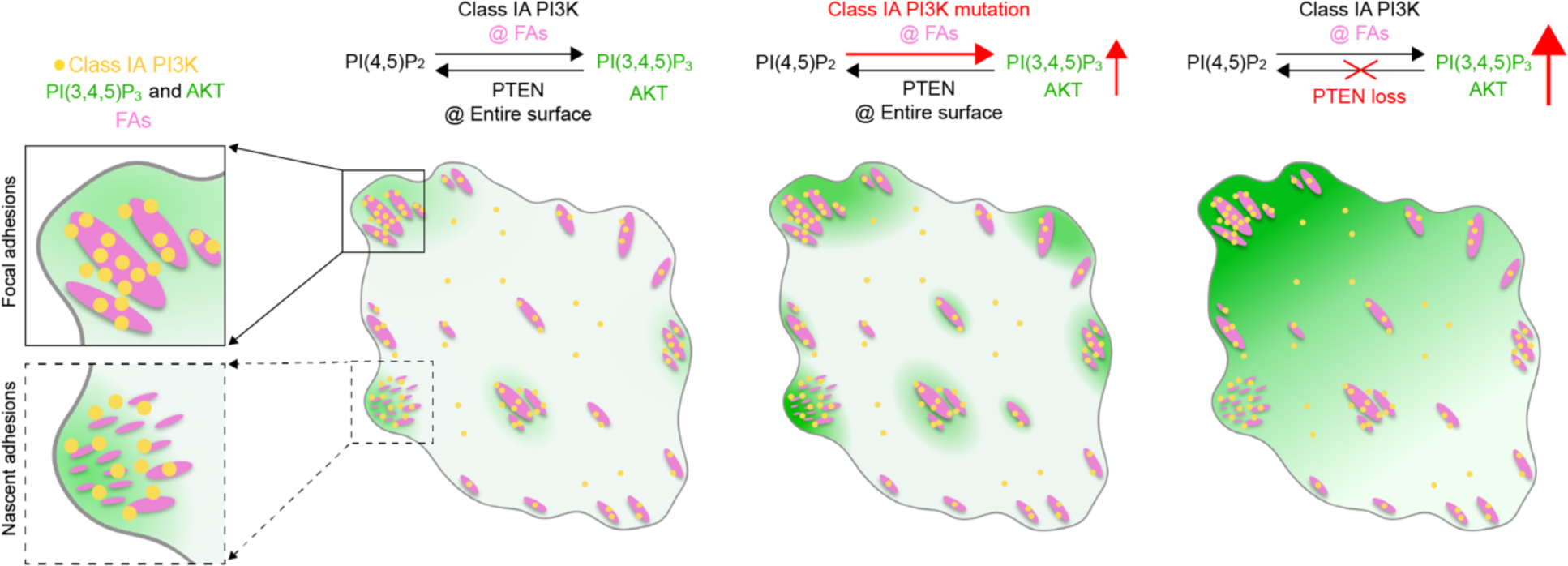
Model of compartmentalized activation of PI3K-PI(3,4,5)P_3_-AKT signaling around focal adhesions. Cell-ECM interaction induces the activation of FAK, which recruits the regulatory p85 subunit and the catalytic p110 subunit of class IA PI3K to nascent and mature FAs. This leads to the local generation of PI(3,4,5)P_3_ in FAs. PI(3,4,5)P_3_ diffusion and elimination by PTEN establish a PI(3,4,5)P_3_ gradient, leading to enrichment centering around clusters of nascent adhesions or larger FAs. Activating mutations of *PIK3CA* further boost PI(3,4,5)P_3_ generation and accumulation around FA structures, whereas the loss of PTEN leads to the accumulation of PI(3,4,5)P_3_ across the entire surface of cells. The local generation and enrichment of PI(3,4,5)P_3_ induces the dynamic recruitment of AKT around FAs and thus leads to the compartmentalized activation of AKT signaling.

Despite decades of extensive structural and biochemical studies, the spatial organization, activation, and regulation of class I PI3K signaling in mammalian cells remain unclear. Most of what we know about the dynamic spatial localization of class I PI3K was obtained from studies in the model organism *Dictyostelium discoideum* during chemotaxis (Comer and Parent, 2002; Funamoto et al., 2002). Previous studies on the static subcellar localization class I PI3K in mammalian cells were primarily carried out by stimulating cells with high concentrations of growth factors such as EGF, platelet-derived growth factor (PDGF), or insulin (Salamon and Backer, 2013; Thapa et al., 2020). However, in the context of serum, tissues, or tumor microenvironments, the concentrations of agonists or growth factors are typically low (usually less than 1 ng/ml) (Pinilla-Macua et al., 2017; Rich et al., 2017). The longstanding critical question is how mammalian cells sense the surrounding mechanical or subtle chemical cues to activate the class I PI3K signaling pathway under basal or physiological conditions. FAs serve as a signaling hub to sense and transduce extracellular biochemical and physical cues to regulate cell adhesion and migration in mammalian cells (Case et al., 2015; Kuo, 2014). As the central mediator of integrin-mediated signaling from FAs, the nonreceptor tyrosine kinase FAK is activated and phosphorylated in response to integrin–ECM engagement during cell attachment (Luo and Guan, 2010; Zhao and Guan, 2011). As shown by us and others (Chen et al., 1996; Chen and Guan, 1994), the p85 subunit of PI3K can interact with the tyrosine phosphorylated FAK. Nevertheless, the p85 regulatory subunit exists in excess over the p110 catalytic subunits in cells and the monomeric p85 executes other essential cellular functions independent of the PI3K complex (Fox et al., 2020; Luo et al., 2005). Whether the association of p85 subunit with phosphorylated FAK leads to the recruitment and activation of class I PI3K complex in FAs has been unclear. Through quantitative single-molecule imaging of genome-edited cells, our most intriguing finding is that both the catalytic and regulatory subunits of class IA PI3K are preferentially and dynamically recruited to FAs for activation in mammalian cells. Meanwhile, phosphatidylinositol phosphate kinase type 1γ (PIPKIγ), a PIP 5-kinase that generates PI(4,5)P_2_ from PI(4)P, is also targeted to FAs (Di Paolo et al., 2002; Ling et al., 2002). In the absence of external agonists, the physical interactions of cells with ECMs stimulate FAK activation, which can lead to the recruitment and activation of class I PI3K in FAs and place it close to its physiological substrate PI(4,5)P_2_. Thus, our findings suggest that the ECM-integrin-FAK mechanical signal acts as a growth factor-independent mechanism for class I PI3K recruitment and activation.

Compartmentalized organization of lipid metabolism at the plasma membrane offers distinct advantages in the precise and efficient spatial control of specificity, intensity, and duration of lipid signaling (Sugiyama et al., 2019). Similar to PI(4,5)P_2_ and other phospholipids, PI(3,4,5)P_3_ diffuses rapidly in the inner leaflet of plasma membranes (∼ 0.1 to 1 μm^2^/s) (Hammond et al., 2009). Thus, PI(3,4,5)P_3_ generated locally in FAs by class IA PI3K would rapidly diffuse to the rest of the plasma membrane. At the same time, these PI(3,4,5)P_3_ molecules would be efficiently depleted by PTEN across the entire inner surface of the cells. Thus, the distinct subcellular distribution of the competing lipid kinase and phosphatase helps maintain a low overall level of PI(3,4,5)P_3_ at the cell membrane while retaining a relatively high level of PI(3,4,5)P_3_ in the nascent adhesions or larger FAs (Figure 7). This compartmentalized enrichment of PI(3,4,5)P_3_ around FAs is also expected to recruit various PI(3,4,5)P_3_ effector proteins around FAs for activation and signaling. Indeed, by taking advantage of genome editing and single-molecule imaging, we uncovered the preferential dynamic recruitment of endogenous AKT molecules around PI(3,4,5)P_3_-enriched FAs. Oncogenic activation mutations of class I PI3Ks further boost the local generation and enrichment of PI(3,4,5)P_3_ and thus AKT activation around FAs. Loss of PTEN resulted in the accumulation of PI(3,4,5)P_3_ at the entire surface of the cells while weakening the compartmentalized distribution of PI(3,4,5)P_3_ and AKT activation around FAs. On the other hand, it has been shown in *Dictyostelium discoideum* that the asymmetric distribution of class I PI3K at the leading edge and PTEN along the lateral sides and rear of the cell helps to establish a polarized accumulation of PI(3,4,5)P_3_ at the anterior side of *Dictyostelium* cells during chemotaxis (Comer and Parent, 2002; Funamoto et al., 2002; Iijima and Devreotes, 2002). Therefore, the distinct spatial distribution of PI(3,4,5)P_3_ and PTEN in mammalian cells is quite different from that in *Dictyostelium* cells during chemotaxis. This is consistent with the notion that *Dictyostelium* cells lack integrin-containing FAs (Mijanovic and Weber, 2022). Thus, it seems that mammalian cells have evolved a unique compartmentalized regulatory mechanism underlying class IA PI3K recruitment, PI(3,4,5)P_3_ generation and AKT signaling activation. Whether the local recruitment of AKT leads to the subcellular enrichment and phosphorylation of various AKT substrates around FAs requires further studies.

Seminal structural or biochemical studies have shown that the oncogenic hotspot mutations can hyperactivate p110α by either weakening its inhibitory interaction with p85α or promoting its binding to membranes (Burke, 2018; Huang et al., 2007; Liu et al., 2022; Vasan et al., 2019). However, direct measurement of the membrane-binding kinetics of wild-type and mutated p110α molecules in live cells has not yet been achieved. One of the most intriguing findings from our study is that p110α molecules with H1047L mutation are dynamically and primarily recruited to FAs for activation under basal conditions. Thus, the single-molecule imaging and analysis method employed in this study offers an ideal system for systematic and comparative analysis of the spatiotemporal kinetics of p110α with different oncogenic mutations in live cells. Meanwhile, FAK is overexpressed and activated in various human cancers and has been considered a promising anticancer drug target, especially in combination therapies (Dawson et al., 2021). Excitingly, we found that inhibition of FAK activity impairs the recruitment of both wild-type and mutant p110α molecules to the plasma membrane. Consistent with this observation, the combined inhibition of FAK and p110α exhibits a stronger inhibitory effect on the viability, proliferation, migration, and invasion of SUM159 cells with H1047L mutation. Therefore, considering the challenges posed by drug tolerance and resistance during PI3K inhibition (Castel et al., 2021; Vanhaesebroeck et al., 2021), the combined inhibition of FAK and class I PI3K may potentially offer a new therapeutic strategy for cancers.

## ACKNOWLEDGMENTS

We thank Dr. Luke D. Lavis for the generous gift of the JFX_650_-HaloTag ligand. We thank Dr. Joan Brugge for generously providing SUM159 cells. We thank Dr. Tamas Balla for the generous gifts of plasmids PH-Btk-GFP and PH-Btk(R28C)-GFP. This research was funded by the National Key R&D Program of China (2022YFA1304500), the National Natural Science Foundation of China (92354305, 32321004, 91957106), and the State Key Laboratory of Molecular Developmental Biology of China.

## RESOURCE AVAILABILITY

### Lead contact

Further information and requests for resources and reagents should be directed to and will be fulfilled by the lead contact, Kangmin He.

### Materials availability

Plasmids and other reagents generated in this study will be available upon request, but a completed Materials Transfer Agreement may be required if there is potential for commercial application.

### Data and code availability

The download links for published MATLAB codes used for imaging analysis were provided in the Method section. Other custom MATLAB routines are available upon request from the lead contact. Any additional information required to reanalyze the data reported in this paper is available from the lead contact upon request.

## EXPERIMENTAL MODEL AND SUBJECT DETAILS

### Cell culture

SUM159 and SUM149 cells were cultured at 37°C and 5% CO_2_ in DMEM/F12 (containing L-glutamine and 15 mM HEPES, Corning), supplemented with 5% FBS (Gibco), 100 U/mL penicillin and streptomycin (Corning), 1 μg/mL hydrocortisone (Sigma-Aldrich), 5 μg/mL insulin (Sigma-Aldrich), pH 7.4. HeLa, T47D, HCT116, and HT-1080 cells were cultured at 37°C and 5% CO_2_ in DMEM (Corning), supplemented with 10% FBS (Gibco), and 100 U/mL penicillin and streptomycin (Corning). Cells were routinely verified to be mycoplasma-free using the TransDetect PCR Mycoplasma Detection Kit (TransGen Biotech).

### Chemicals and treatment

Alpelisib (MedChemExpress, HY-15244, 3 μM), dasatinib (Targetmol, T1448, 50 nM), defactinib (Targetmol, T1996, 5 μM), TGX-221 (MedChemExpress, HY-10114, 3 μM), and 14z (MedChemExpress, HY-128580, 0.5 μM), rapamycin (Merck Millipore, 553210-100UG, 1 μM), or EGF (PeproTech, AF-100-15, 100 ng/mL) were added to cells during continuous imaging using a syringe pump (Harvard Apparatus). In the western blot analysis, 1 μM of alpelisib was used.

### Plasmids

The DNA sequences encoding the residues 2-177 and 2-166 of human Btk were amplified by PCR from PH-Btk-GFP (a kind gift from Dr. Tamas Balla) (Varnai et al., 1999) and inserted into a vector containing mEGFP to generate the plasmids mEGFP-Btk(2-177) and mEGFP-Btk(2-166) using the Gibson assembly method (NEBuilder HiFi DNA Assembly Master Mix, New England Biolabs; pEASY-Uni Seamless Cloning and Assembly Kit, TransGen Biotech). Two DNA fragments containing residues 2-166 were amplified by PCR from PH-Btk-GFP and inserted into a vector containing mEGFP to create the plasmid mEGFP-2xBtk(2-166) using the Gibson assembly method. A flexible (GGS)_5_ linker (5’-GGAGGATCCGGTGGATCTGGAGGTTCTGGTGGTTCTGGTGGTTCC-3’) was inserted between mEGFP and the first Btk(2-166) fragment. A second linker (5’-GGGGGATCGGGTGGTGTCGAC-3’) was placed between the first and second Btk(2-166) fragments. Similar methods were used to construct mEGFP-tagged 2xBtk(2-177), 2xBtk(2-170), 2xBtk(2-168), 2xBtk(2-166), 2xBtk(2-165), and 2xBtk(2-163). The DNA sequences encoding the residues 2-177 and 2-166 of human Btk with R28C mutation were amplified by PCR from PH-Btk(R28C)-GFP (Varnai et al., 1999) and inserted into a vector containing mEGFP to generate the plasmids mEGFP-2xBtk(2-177)-R28C and mEGFP-2xBtk(2-166)-R28C by the Gibson assembly method. Halo-2xBtk(2-166) and mScarlet-I-2xBtk(2-166) were generated by replacing the coding sequence of mEGFP in plasmid mEGFP-2xBtk(2-166) with the coding sequence of HaloTag or mScarlet-I.

The DNA sequences encoding the PH domain of human PLCδ1 were amplified by PCR from EGFP-PH(PLCδ1) and inserted into a vector containing mScarlet-I or HaloTag to create the plasmids mScarlet-I-PH(PLCδ1) and Halo-PH(PLCδ1). The DNA sequences encoding the full-length human p110α, p110β, p110δ, and p110γ were amplified by PCR and fused at the N-terminus of the coding sequence of mNeonGreen (Shaner et al., 2013) to generate the plasmids p110α-mNeonGreen, p110β-mNeonGreen, p110δ-mNeonGreen, and p110γ-mNeonGreen. DNA sequences encoding the full-length human p85⍺, p85β, or AKT1 were amplified by PCR and fused at the C-terminus of the coding sequence of mNeonGreen or mEGFP to generate the plasmids mNeonGreen-p85⍺, mNeonGreen-p85β, mEGFP-p85⍺, and mNeonGreen-AKT1. The DNA sequence encoding the full-length human FAK was amplified by PCR and fused at the C-terminus of the coding sequence of HaloTag to generate the plasmid Halo-FAK. The DNA sequences encoding human PTEN or PTEN-G129E were amplified by PCR and fused at the C-terminus of the coding sequence of mScarlet-I to generate the plasmids mScarlet-I-PTEN or mScarlet-I-PTEN-G129E. A flexible linker was inserted between mScarlet-I/HaloTag/mNeonGreen/mEGFP/mCherry and the DNA fragments.

LYN11-FRB-ECFP (#38003), mCherry-FKBP-INPP4B (#116864), NES-EGFP-cPHx3 (#116855), mCherry-FKBP-PTEN (#116866), mCherry-CRY2-iSH2 (#66839), mCherry-α-tubulin (#21043), mCherry-paxillin (#50526), mCherry-FAK (#55122), mCherry-vinculin (#55159) were obtained from Addgene. Plasmid transfection was performed using Lipofectamine 3000 (Invitrogen) according to the manufacturer’s instructions.

### Generation of fluorescent tag knock-in cell lines using the CRISPR/Cas9 approach

SUM159 cells were gene-edited to incorporate mNeonGreen at the C-terminus of p110α or p110β, mEGFP/HaloTag at the N-terminus of p85α or p85β, mNeonGreen at the N-terminus of AKT1, mEGFP at the N-terminus of FAK, and EGFP at the C-terminus of paxillin using the CRISPR/Cas9 approach as described (Ran et al., 2013). A single-guide RNA (sgRNA) targeting human *PIK3CA* (5’-ATGCATGCTGTTTAATTGTG-3’), human *PIK3CB* (5’-AACAAATACATTAGGAGCGA-3’), human *PIK3R1* (5’-ATTTGCAAACATGAGTGCTG-3’), human *PIK3R2* (5’-CCCGCCATGGCCGCGTGAGT-3’), human *AKT1* (5’-CTCGGGCACCATGAGCGACG-3’), human *FAK* (5’-CCTAGCATCTAGCAAAATAA-3’), or human *PXN* (5’-CAAGCTCTTCTGCTAGGTGC-3’) was cloned into pSpCas9(BB)-2A-Puro (PX459) (Addgene). Donor constructs used for homologous recombination were generated by cloning into the pUC19 vector with two ∼600-800-nucleotide fragments of genomic DNA upstream and downstream of the stop codon of human *PIK3CA* and *PIK3CB*, or the start codon of human *PIK3R1*, *PIK3R2*, *AKT1*, and *FAK*, and the open reading frame of mNeonGreen/mEGFP/Halo using the pEASY-Uni Seamless Cloning and Assembly Kit (TransGen Biotech). A flexible (GGS)_3_ or (GGS)_5_ linker was inserted between the start or stop codon of the gene and the open reading frame of mNeonGreen/mEGFP/Halo. The donor plasmid paxillin-EGFP was obtained from Addgene (#87420).

SUM159 cells were transfected with the donor plasmid and PX459 plasmid containing the sgRNA targeting sequence using Lipofectamine 3000 (Invitrogen). Cells expressing mNeonGreen, mEGFP, or HaloTag (labeled with the JF_549_-HaloTag ligand, Promega) were enriched by fluorescence-activated cell sorting (FACS) 5-7 days after transfection. The sorted cells were expanded and then subjected to single-cell sorting into 96-well plates (FACSAria II, BD Biosciences). The gene-edited clonal SUM159 cells were confirmed by imaging, PCR, and western blot analysis. Gene-edited cells expressing EGFR-Halo were generated as described (Bai et al., 2023).

### Mutation of *PIK3CA* in SUM159 cells using the CRISPR/Cas9 approach

SUM159 cells (*PIK3CA*^WT/H1047L^) were gene-edited to either correct the H1047L mutation (*PIK3CA*^WT/WT^) or introduce the H1047L mutation in the other allele (*PIK3CA*^H1047L/H1047L^) using the CRISPR/Cas9 approach as described above. The sgRNA sequence 5’-ATGAATGATGCACaTCATGG-3’ or 5’-ATGAATGATGCACtTCATGG-3’ was cloned into pSpCas9(BB)-2A-GFP (PX458) (Addgene) to generate the SUM159 cell lines with *PIK3CA*^H1047L/H1047L^ or *PIK3CA*^WT/WT^, respectively. The donor constructs, with or without H1047L mutation, used for homologous recombination were generated by cloning into the pUC19 vector with two ∼600-nucleotide fragments of genomic DNA upstream and downstream of H1047 of human *PIK3CA* using the pEASY-Uni Seamless Cloning and Assembly Kit (TransGen Biotech). SUM159 cells were transfected with the donor plasmid and PX458 plasmid containing the sgRNA targeting sequence using Lipofectamine 3000 (Invitrogen). Cells expressing GFP were sorted into 96-well plates 24 to 36 hours after transfection using single-cell sorting (FACSAria II, BD Biosciences). The monoclonal cell populations without or with H1047L mutations in both alleles were identified by sequencing.

### Knockout of PTEN or SHIP2 in SUM159 cells

Knockout of PTEN or SHIP2 was performed using the CRISPR/Cas9 approach as described (Ran et al., 2013). The sgRNA targeting human *PTEN* (5’-CCAGGGAGTAACTATTCCCA-3’) or human *SHIP2* (5’-GATGGTCTTGGCCTTACGTG-3’) were cloned into pSpCas9(BB)-2A-GFP (PX458) (Addgene, #48138). After 24-36 hours of transfection, SUM159 cells expressing GFP were subjected to single-cell sorting into 96-well plates (FACSAria II, BD Biosciences). Two to three weeks later, the monoclonal cell populations with mutations in both alleles were identified by sequencing, and the loss of PTEN or SHIP2 protein expression was confirmed by western blot analysis.

### Generation of stable cell lines

SUM159 cells stably expressing the PI(3,4,5)P_3_ sensor mNeonGreen-2xBtk(2-166) were generated by transduction using a lentiviral vector encoding mNeonGreen-2xBtk(2-166). Pools of cells expressing mNeonGreen-2xBtk(2-166) were identified by FACS 5-7 days after transduction, followed by two additional bulk FACS sorts to obtain a pool of cells stably expressing relatively low levels of the sensor.

### Knockdown of FAK or PYK2 using shRNA

Lentivirus shRNA containing the target sequence 5’-GATGTTGGTTTAAAGCGATTT-3’ or 5’-CGTATCCTCAAGGTCTGCTTC-3’ was utilized to knock down the expression of FAK or PYK2. The shRNA was generated by cloning the target sequence into the pLKO.1 - TRC cloning vector (Addgene, #10878). The scramble shRNA expressing 5’-CCTAAGGTTAAGTCGCCCTCG-3’ (Addgene, #1864) was used as a control. shRNA-expressing lentivirus was produced in HEK293FT cells. The supernatant containing lentivirus was collected 48 hours after transfection and added to SUM159 cells. After 24 hours, the cells were replaced with fresh medium containing 2 μg/mL puromycin (InvivoGen). After another 96 hours, the cells were subjected to live-cell imaging and western blot analysis.

### Live-cell imaging of PI(3,4,5)P_3_ distribution and dynamics at the plasma membrane

The bottom surfaces of cells were imaged using TIRF microscopy built on either a Nikon Ti2-E microscope or a Nikon TIE microscope. The Nikon Ti2-E microscope was equipped with a manual TIRF Illuminator Unit (Nikon), a CFI Apochromat TIRF 100X objective (1.49 NA, Nikon), a Perfect Focus Unit (Nikon), an UNO Stage Top Incubator (Okolab), OBIS CellX lasers (405, 488, 561 and 637 nm, Coherent), the W-VIEW GEMINI-2C Image Splitting Optics (Hamamatsu), and two EMCCD cameras (Evolve 512 Delta, Photometrics). Images were acquired using Micro-Manager 1.4 (Edelstein et al., 2010). The Nikon TiE microscope was equipped with a motorized TIRF Illuminator Unit (Nikon), a CFI Apochromat TIRF 100X objective (1.49 NA, Nikon), a Perfect Focus Unit (Nikon), a Motorized XY stage (Prior Scientific), a fully enclosed and environmentally controlled cage incubator (Okolab), OBIS 488, 561 and 647 nm lasers (Coherent), the W-VIEW GEMINI Image splitting optics (Hamamatsu), and an EMCCD camera (iXon Life 888, Andor Technology). Imaging sequences of a single position or multi-positions were acquired using Micro-Manager 2.0 (Edelstein et al., 2010).

Cells were transfected with the PI(3,4,5)P_3_ sensors and other plasmids using Lipofectamine 3000 (Invitrogen) and then plated on single-well, 4-well, or 8-well confocal dishes (Cellvis) 6-8 hours after transfection. SUM159 cells with relatively low levels of sensor expression were imaged the next day in phenol-free DMEM/F12 (Corning) containing 5% FBS and 20 mM HEPES. For serum starvation, SUM159 cells were washed with DMEM/F12 once and then cultured in DMEM/F12 overnight. T47D, HCT116, HeLa, and HT-1080 cells were imaged in FluoroBrite DMEM (Thermo Fisher Scientific) containing 10% FBS and 20 mM HEPES. Montages and intensity measurements were performed using Fiji (Schindelin et al., 2012). Kymographs were created using the Multi Kymograph plugin in Fiji.

### Quantification of sensor distribution at the plasma membrane

To quantify the membrane distribution of the Btk sensors around the cell periphery, the cell boundary was initially segmented based on the fluorescence of the PI(4,5)P_2_ sensor using either CellProfiler 4 (Stirling et al., 2021) (http://www.cellprofiler.org) or manual segmentation (also see Figure S4A). Subsequently, a 20-pixel eroded mask was generated using the ‘imerode’ function (disk-shaped structuring element) in MATLAB (MathWorks), where small objects (<2500 pixels) were removed. Based on the border of the eroded mask, the intensity profile along the eroded line was then measured using the built-in ‘Plot Profile’ function (line width of 40 pixels) in Fiji. The intensity profile was normalized to its average and interpolated to 1,000 data points to maintain a consistent profile length. To create an intensity heatmap, intensity profiles from 60 randomly selected cells from each experiment were plotted side by side. The color bar ranging from 0 to 4.5 was used for all intensity profile heatmaps. The standard deviation of the intensity profile from each cell was calculated to describe the overall extent of PI(3,4,5)P_3_ uneven distribution at the plasma membrane.

### Calculation of cell boundary velocity and sensor intensity

The boundary protrusion/retraction velocity and fluorescence intensity were determined using BoundaryTrack (https://github.com/fjmrt/BoundaryTrack) in MATLAB. Briefly, the boundary coordinates of an object were extracted and segmented into an arbitrary number of positions and then mapped clockwise between time frames to minimize the sum of square distance. To calculate the intensity of the PI(3,4,5)P_3_ sensor along the cell boundary, a 20-pixel eroded line based on the cell boundary was generated for each frame, and the intensity values were calculated (neighbor = 1) as described in BoundaryTrack.

### Focal adhesion analysis

The detection and segmentation of FAs were conducted using the online Focal Adhesion Analysis Server (FAAS) (https://faas.bme.unc.edu) or locally in MATLAB (https://github.com/mbergins/focal_adhesions) (Berginski and Gomez, 2013). The intensities of detected FAs and non-FA regions were measured and analyzed in Fiji and MATLAB. Montages of FAs in selected frames were generated using FAAS in MATLAB. Each segmented adhesion was color-coded based on its mean intensity from the original image. Overlays of all FAs, color-coded by their appearance time in the time series, were generated using the online FAAS.

Randomization of detected FAs at the plasma membrane was performed using the algorithm as described (Vega-Lugo et al., 2022). Briefly, the segmented FAs were randomly distributed inside an eroded cell mask region, sequentially from the largest to the smallest. Non-FA regions within the cell boundary were generated using the ‘bubblebath’ function (https://www.mathworks.com/matlabcentral/fileexchange/70348) in MATLAB. Circles with a specified radius range were randomly drawn, with no overlap, in an image of the same size as the original image. Circles outside the cell boundary and circles overlapping with or too close to (within 5 pixels) detected FAs were discarded.

### Single-molecule imaging and analysis of p110α recruitment at the plasma membrane

Gene-edited p110α-mNeonGreen^+/-^ cells transiently expressing mCherry-FAK were imaged every 200 ms for 601 frames (exposure time 100 ms for each channel) using the Nikon Ti2-E TIRF microscope with single-molecule sensitivity. The last 301 frames of each time series were used for further imaging analysis. Single p110α-mNeonGreen fluorescent spots were detected and tracked by u-track (https://github.com/DanuserLab/u-track) (Jaqaman et al., 2008) in MATLAB. Parameters with non-default values used: psfSigma: 1.25; mergeSplit: 0; minSearchRadius: 2; maxSearchRadius: 6; maximum allowed gap: 3. FA and Non-FA tracks were then extracted based on spot coordinates and FA detection using local FAAS as described above. Tracks longer than 2 (≥ 3) frames were used to calculate track initiation rate and lifetimes. Tracks longer than 5 (≥ 6) frames were further used for mean square displacement (MSD) analysis using the MSDanalyzer package (https://tinevez.github.io/msdanalyzer) (Tarantino et al., 2014) in MATLAB. The motion states (immobile, confined, free) classification of each track was based on the moment scaling spectrum from u-track results.

### Single-molecule imaging and analysis of AKT1 recruitment at the plasma membrane

Gene-edited mNeonGreen-AKT1^+/+^ cells transiently expressing mCherry-FAK were firstly imaged for 10 frames (both channels) using the Nikon Ti2-E TIRF microscope to capture the distribution of FAK at the plasma membrane. The same cells were subsequently imaged in the mNeonGreen channel every 50 ms for 601 frames. Single mNeonGreen-AKT1 fluorescent spots in the time series were detected and tracked by u-track in MATLAB. Parameters with non-default values used: psfSigma: 1.33; mergeSplit: 0; minSearchRadius: 2; maxSearchRadius: 4; maximum allowed gap: 4. FAs were detected by FAAS as described above. FAs were dilated by 0.8 μm to get regions around FAs. Tracks longer than 5 (≥ 6) frames were used to calculate track density and diffusion rate. MSD and diffusion coefficients (*D*) were calculated using MSDanalyzer package. Track density maps and diffusion maps were generated using the InferenceMAP software (https://research.pasteur.fr/en/software/inferencemap/) (El Beheiry et al., 2015).

### Live-cell imaging with spinning-disk confocal microscopy

The spinning-disk confocal microscopy system was built on the Nikon TiE microscope described above. The microscope was equipped with a CSU-X1 spinning-disk confocal unit (Yokogawa) and an EMCCD camera (iXon Ultra 897, Andor Technology), positioned on the left side port of the microscope.

### Quantification of PI(3,4,5)P_3_ levels by mass spectrometry

Quantification of PI(3,4,5)P_3_ was performed based on the LC-MRM method as previously described (Clark et al., 2011). Lipids were extracted under acidic conditions to increase recovery of phosphoinositides from biological tissues (Boss and Massel, 1985). In brief, cells were incubated with an extraction solvent containing chloroform: methanol (1:1)+ 0.5N HCl and 2 mM AlCl_3_ for 10 min at 1,500 rpm. Internal standard cocktail containing 17:0/20:4 PI3P, 17:0/20:4 PI4P, 17:0/20:4 PI5P, 17:0/20:4 PI(3,4)P_2_, 17:0/20:4 PI(3,5)P_2_, 17:0/20:4 PI(4,5)P_2_ and 17:0/20:4 PI(3,4,5)P_3_ (Avanti Polar Lipids) was added into individual samples during extraction. At the end of incubation, deionized water was added to induce phase separation and the samples were centrifuged. The lower organic phase containing lipids was extracted and transferred to a new tube. The extraction procedure was repeated for three rounds. The pooled extract was derivatized with 2M TMS-diazomethane in hexane according to a published protocol (Clark et al., 2011) and analyzed on an Exion AD-UPLC coupled to Sciex QTRAP 6500 Plus. Phosphoinositides were separated on a Phenomenex Kinetex C18 column (2.6 µm, 100×2.1 mm) using water: acetonitrile (1:1) containing 2 mM ammonium acetate and 0.1% (v/v) formic acid as mobile phase A, and acetonitrile: isopropanol (1:1) as mobile phase B. Endogenous phosphoinositides were quantitated by reference to the levels of internal standards added during lipid extraction.

### Transwell migration and invasion assays

Transwell migration and invasion assays were performed using chambers with a diameter of 6.5 mm and pore size of 8.0 μm (Corning). In brief, cells were cultured with DMSO, alpelisib (1 μM), defactinib (5 μM), or a combination of alpelisib and defactinib (1 μM and 5 μM) in 6-well plates for 48 hours. Subsequently, 200,000 cells were suspended in 200 μL of serum-free medium and seeded in the upper chamber. The lower chamber was filled with 650 μL of culture medium containing 20% FBS. After a 24-hour incubation at 37°C and 5% CO_2_, the filters were fixed with methanol for 2 min. Then, the filters were stained with crystal violet solution for 10 min. Non-migratory cells were gently removed from the upper side of the membrane using a cotton swab. For the invasion assay, the upper chamber membrane was pre-coated with 50 µL serum-free medium containing 2.5 mg/mL of Matrigel (BD). The subsequent steps were similar to the migration assay. Finally, the membranes were imaged by a Leica DMi8 microscope with a 10x objective.

### Cell viability and proliferation assays

The cell viability assay was conducted using the Cell Counting Kit 8 (CCK-8, TargetMol) according to the manufacturer’s instructions. Cell activity was measured 72 hours after incubating the cells with DMSO, alpelisib (1 μM), defactinib (5 μM), or a combination of alpelisib and defactinib (1 μM and 5 μM) in the culture medium. The growth curves of SUM159 cells were generated using the xCELLigence RTCA MP system (Agilent). Approximately 2,000 cells were seeded in one well of a 16-well E-plate (Agilent), and DMSO or different inhibitors were added 24 h later. The index of adherent cells in each well was measured every hour to generate the growth curves.

### Western blot analysis

Cells were solubilized at 4°C for 30 min in RIPA lysis buffer (50 mM Tris-HCl, pH 8.0, 150 mM NaCl, 1% NP-40, 0.5% sodium deoxycholate, 0.1% SDS) with a protease and phosphatase inhibitor cocktail (Thermo Fisher Scientific, 78442) and then pelleted at 13,400 g for 15 min at 4 °C. The supernatant was mixed with 5× sample buffer (GenScript, MB01015), heated to 100°C for 10 min, and then fractionated by SDS–PAGE and transferred to nitrocellulose membranes (Pall, 66485). The membranes were incubated in TBST buffer containing 5% skim milk for 3 hours at room temperature, followed by overnight incubation at 4 °C with specific primary antibodies. After three washes with TBST (5 min each), the membranes were incubated with the appropriate HRP-conjugated secondary antibody (Beyotime, A0208, or A0216, 1:1,000) at room temperature for 1 hour. The membranes were incubated with the SignalFire^TM^ ECL Reagent (Cell Signaling Technology, 6883S) and imaged using the MiniChemi 610 chemiluminescent imaging system (Sage Creation Science). Primary antibodies against AKT (Cell Signaling Technology, 4691, 1:1,000), pAKT (Cell Signaling Technology, 4060, 1:2,000), AKT1 (Cell Signaling Technology, 2938, 1:1,000), GAPDH (Proteintech, 60004-1-Ig, 1:50,000), pFAK (Invitrogen, 44-624G, 1:1,000), FAK (Proteintech, 66258-1-Ig, 1:3,000), PYK2 (Cell Signaling Technology, 3292S, 1:1,000), p110α (Proteintech, 67071-1-Ig, 1:2,000), p110β (Abcam, ab151549, 1:1,000), p85α (Proteintech, 60225-1-Ig, 1:5,000), p85β (ABclonal, A4860, 1:500), PTEN (Proteintech, 22034-1-AP, 1:2,000), and GFP (TransGen Biotech, HT801-01, 1:3,000) were used in this study.

### Co-immunoprecipitation (co-IP)

Cells transfected with the specified plasmids and pretreated with inhibitors were lysed in a lysis buffer supplemented with a cocktail of protease and phosphatase inhibitors (Thermo Fisher Scientific, 78442). After incubating on ice for 30 min, the cells were centrifuged at 20,000 g for 15 min at 4℃. Supernatants were collected and incubated with GFP-Nanoab-Agarose beads (Lablead, GNA-25-500) with gentle rotation for 6 h at 4℃. Then the beads were washed three times with washing buffer, and the bound proteins were eluted using 2× SDS sample buffer. The samples were then subjected to immunoblot analysis.

## QUANTIFICATION AND STATISTICAL ANALYSIS

Statistical analyses were performed using GraphPad Prism 9 (GraphPad Software). The two-sided Student’s t test was used to compare two groups. The ordinary one-way ANOVA, followed by Tukey’s multiple-comparisons test, was used to compare three or four groups. *: *P* < 0.05; **: *P* < 0.01; ****: *P* < 0.0001; ns: no significant difference. *P* < 0.05 was considered statistically significant. No statistical methods were used to predetermine sample sizes.

## References

Aytenfisu, T.Y., Campbell, H.M., Chakrabarti, M., Amzel, L.M., and Gabelli, S.B. (2022). Class I PI3K Biology. Curr Top Microbiol Immunol 436, 3–49.

Backer, J.M. (2010). The regulation of class IA PI 3-kinases by inter-subunit interactions. Curr Top Microbiol Immunol 346, 87–114.

Bai, X., Sun, P., Wang, X., Long, C., Liao, S., Dang, S., Zhuang, S., Du, Y., Zhang, X., Li, N., et al. (2023). Structure and dynamics of the EGFR/HER2 heterodimer. Cell Discov 9, 18.

Bailey, M.H., Tokheim, C., Porta-Pardo, E., Sengupta, S., Bertrand, D., Weerasinghe, A., Colaprico, A., Wendl, M.C., Kim, J., Reardon, B., et al. (2018). Comprehensive Characterization of Cancer Driver Genes and Mutations. Cell 174, 1034–1035.

Balla, T. (2013). Phosphoinositides: tiny lipids with giant impact on cell regulation. Physiol Rev 93, 1019–1137.

Batrouni, A.G., and Baskin, J.M. (2020). A MAP for PI3K activation on endosomes. Nat Cell Biol 22, 1292–1294.

Berginski, M.E., and Gomez, S.M. (2013). The Focal Adhesion Analysis Server: a web tool for analyzing focal adhesion dynamics. F1000Res 2, 68.

Bilanges, B., Posor, Y., and Vanhaesebroeck, B. (2019). PI3K isoforms in cell signalling and vesicle trafficking. Nat Rev Mol Cell Biol 20, 515–534.

Boss, W.F., and Massel, M.O. (1985). Polyphosphoinositides are present in plant tissue culture cells. Biochem Biophys Res Commun 132, 1018–1023.

Burke, J.E. (2018). Structural Basis for Regulation of Phosphoinositide Kinases and Their Involvement in Human Disease. Mol Cell 71, 653–673.

Burke, J.E., and Williams, R.L. (2015). Synergy in activating class I PI3Ks. Trends Biochem Sci 40, 88–100.

Case, L.B., Baird, M.A., Shtengel, G., Campbell, S.L., Hess, H.F., Davidson, M.W., and Waterman, C.M. (2015). Molecular mechanism of vinculin activation and nanoscale spatial organization in focal adhesions. Nature Cell Biology 17, 880–892.

Castel, P., Toska, E., Engelman, J.A., and Scaltriti, M. (2021). The present and future of PI3K inhibitors for cancer therapy. Nat Cancer 2, 587–597.

Chen, H.C., Appeddu, P.A., Isoda, H., and Guan, J.L. (1996). Phosphorylation of tyrosine 397 in focal adhesion kinase is required for binding phosphatidylinositol 3-kinase. J Biol Chem 271, 26329–26334.

Chen, H.C., and Guan, J.L. (1994). Association of focal adhesion kinase with its potential substrate phosphatidylinositol 3-kinase. Proc Natl Acad Sci U S A 91, 10148–10152.

Chung, J.K., Nocka, L.M., Decker, A., Wang, Q., Kadlecek, T.A., Weiss, A., Kuriyan, J., and Groves, J.T. (2019). Switch-like activation of Bruton’s tyrosine kinase by membrane-mediated dimerization. Proc Natl Acad Sci U S A 116, 10798–10803.

Clark, J., Anderson, K.E., Juvin, V., Smith, T.S., Karpe, F., Wakelam, M.J., Stephens, L.R., and Hawkins, P.T. (2011). Quantification of PtdInsP3 molecular species in cells and tissues by mass spectrometry. Nat Methods 8, 267–272.

Comer, F.I., and Parent, C.A. (2002). PI 3-kinases and PTEN: how opposites chemoattract. Cell 109, 541–544.

Dawson, J.C., Serrels, A., Stupack, D.G., Schlaepfer, D.D., and Frame, M.C. (2021). Targeting FAK in anticancer combination therapies. Nat Rev Cancer 21, 313–324.

Di Paolo, G., Pellegrini, L., Letinic, K., Cestra, G., Zoncu, R., Voronov, S., Chang, S., Guo, J., Wenk, M.R., and De Camilli, P. (2002). Recruitment and regulation of phosphatidylinositol phosphate kinase type 1 gamma by the FERM domain of talin. Nature 420, 85–89.

Ebner, M., Lucic, I., Leonard, T.A., and Yudushkin, I. (2017a). PI(3,4,5)P3 Engagement Restricts Akt Activity to Cellular Membranes. Mol Cell 65, 416–431 e416.

Ebner, M., Lucic, I., Leonard, T.A., and Yudushkin, I. (2017b). PI(3,4,5)P(3) Engagement Restricts Akt Activity to Cellular Membranes. Mol Cell 65, 416–431 e416.

Edelstein, A., Amodaj, N., Hoover, K., Vale, R., and Stuurman, N. (2010). Computer control of microscopes using µManager. Curr Protoc Mol Biol Chapter 14, Unit14.20.

El Beheiry, M., Dahan, M., and Masson, J.B. (2015). InferenceMAP: mapping of single-molecule dynamics with Bayesian inference. Nat Methods 12, 594–595.

Fox, M., Mott, H.R., and Owen, D. (2020). Class IA PI3K regulatory subunits: p110-independent roles and structures. Biochem Soc Trans 48, 1397–1417.

Franke, T.F., Kaplan, D.R., Cantley, L.C., and Toker, A. (1997). Direct regulation of the Akt proto-oncogene product by phosphatidylinositol-3,4-bisphosphate. Science 275, 665–668.

Fruman, D.A., Chiu, H., Hopkins, B.D., Bagrodia, S., Cantley, L.C., and Abraham, R.T. (2017). The PI3K Pathway in Human Disease. Cell 170, 605–635.

Funamoto, S., Meili, R., Lee, S., Parry, L., and Firtel, R.A. (2002). Spatial and temporal regulation of 3-phosphoinositides by PI 3-kinase and PTEN mediates chemotaxis. Cell 109, 611–623.

Gao, X., Lowry, P.R., Zhou, X., Depry, C., Wei, Z., Wong, G.W., and Zhang, J. (2011). PI3K/Akt signaling requires spatial compartmentalization in plasma membrane microdomains. Proc Natl Acad Sci U S A 108, 14509–14514.

Geiger, B., Spatz, J.P., and Bershadsky, A.D. (2009). Environmental sensing through focal adhesions. Nat Rev Mol Cell Biol 10, 21–33.

Gilby, D.C., Goodeve, A.C., Winship, P.R., Valk, P.J., Delwel, R., and Reilly, J.T. (2007). Gene structure, expression profiling and mutation analysis of the tumour suppressor SHIP1 in Caucasian acute myeloid leukaemia. Leukemia 21, 2390–2393.

Goulden, B.D., Pacheco, J., Dull, A., Zewe, J.P., Deiters, A., and Hammond, G.R.V. (2019). A high-avidity biosensor reveals plasma membrane PI(3,4)P2 is predominantly a class I PI3K signaling product. J Cell Biol 218, 1066–1079.

Guo, H., German, P., Bai, S., Barnes, S., Guo, W., Qi, X., Lou, H., Liang, J., Jonasch, E., Mills, G.B., et al. (2015). The PI3K/AKT Pathway and Renal Cell Carcinoma. J Genet Genomics 42, 343–353.

Hammond, G.R., Sim, Y., Lagnado, L., and Irvine, R.F. (2009). Reversible binding and rapid diffusion of proteins in complex with inositol lipids serves to coordinate free movement with spatial information. J Cell Biol 184, 297–308.

Hollestelle, A., Elstrodt, F., Nagel, J.H., Kallemeijn, W.W., and Schutte, M. (2007). Phosphatidylinositol-3-OH kinase or RAS pathway mutations in human breast cancer cell lines. Mol Cancer Res 5, 195–201.

Huang, C.H., Mandelker, D., Schmidt-Kittler, O., Samuels, Y., Velculescu, V.E., Kinzler, K.W., Vogelstein, B., Gabelli, S.B., and Amzel, L.M. (2007). The structure of a human p110alpha/p85alpha complex elucidates the effects of oncogenic PI3Kalpha mutations. Science 318, 1744–1748.

Iijima, M., and Devreotes, P. (2002). Tumor suppressor PTEN mediates sensing of chemoattractant gradients. Cell 109, 599–610.

Isakoff, S.J., Engelman, J.A., Irie, H.Y., Luo, J., Brachmann, S.M., Pearline, R.V., Cantley, L.C., and Brugge, J.S. (2005). Breast cancer-associated PIK3CA mutations are oncogenic in mammary epithelial cells. Cancer Res 65, 10992–11000.

Jaqaman, K., Loerke, D., Mettlen, M., Kuwata, H., Grinstein, S., Schmid, S.L., and Danuser, G. (2008). Robust single-particle tracking in live-cell time-lapse sequences. Nat Methods 5, 695–702.

Kang, S., Bader, A.G., and Vogt, P.K. (2005). Phosphatidylinositol 3-kinase mutations identified in human cancer are oncogenic. Proc Natl Acad Sci U S A 102, 802–807.

Kuo, J.C. (2014). Focal adhesions function as a mechanosensor. Prog Mol Biol Transl Sci 126, 55–73.

Lawrence, M.S., Stojanov, P., Mermel, C.H., Robinson, J.T., Garraway, L.A., Golub, T.R., Meyerson, M., Gabriel, S.B., Lander, E.S., and Getz, G. (2014). Discovery and saturation analysis of cancer genes across 21 tumour types. Nature 505, 495–501.

Layton, M.J., Rynkiewicz, N.K., Ivetac, I., Horan, K.A., Mitchell, C.A., and Phillips, W.A. (2014). Assessing the subcellular distribution of oncogenic phosphoinositide 3-kinase using microinjection into live cells. Biosci Rep 34.

Lee, Y.R., Chen, M., and Pandolfi, P.P. (2018). The functions and regulation of the PTEN tumour suppressor: new modes and prospects. Nat Rev Mol Cell Biol 19, 547–562.

Lietha, D., Cai, X., Ceccarelli, D.F., Li, Y., Schaller, M.D., and Eck, M.J. (2007). Structural basis for the autoinhibition of focal adhesion kinase. Cell 129, 1177–1187.

Ling, K., Doughman, R.L., Firestone, A.J., Bunce, M.W., and Anderson, R.A. (2002). Type I gamma phosphatidylinositol phosphate kinase targets and regulates focal adhesions. Nature 420, 89–93.

Liu, S.L., Wang, Z.G., Hu, Y., Xin, Y., Singaram, I., Gorai, S., Zhou, X., Shim, Y., Min, J.H., Gong, L.W., et al. (2018). Quantitative Lipid Imaging Reveals a New Signaling Function of Phosphatidylinositol-3,4-Bisphophate: Isoform- and Site-Specific Activation of Akt. Mol Cell 71, 1092–1104 e1095.

Liu, X., Yang, S., Hart, J.R., Xu, Y., Zou, X., Zhang, H., Zhou, Q., Xia, T., Zhang, Y., Yang, D., et al. (2021). Cryo-EM structures of PI3Kalpha reveal conformational changes during inhibition and activation. Proc Natl Acad Sci U S A 118.

Liu, X., Zhou, Q., Hart, J.R., Xu, Y., Yang, S., Yang, D., Vogt, P.K., and Wang, M.W. (2022). Cryo-EM structures of cancer-specific helical and kinase domain mutations of PI3Kalpha. Proc Natl Acad Sci U S A 119, e2215621119.

Lucic, I., Rathinaswamy, M.K., Truebestein, L., Hamelin, D.J., Burke, J.E., and Leonard, T.A. (2018). Conformational sampling of membranes by Akt controls its activation and inactivation. Proc Natl Acad Sci U S A 115, E3940–E3949.

Luo, J., Field, S.J., Lee, J.Y., Engelman, J.A., and Cantley, L.C. (2005). The p85 regulatory subunit of phosphoinositide 3-kinase down-regulates IRS-1 signaling via the formation of a sequestration complex. J Cell Biol 170, 455–464.

Luo, M., and Guan, J.L. (2010). Focal adhesion kinase: A prominent determinant in breast cancer initiation, progression and metastasis. Cancer Lett 289, 127–139.

Ma, A., Richardson, A., Schaefer, E.M., and Parsons, J.T. (2001). Serine phosphorylation of focal adhesion kinase in interphase and mitosis: a possible role in modulating binding to p130(Cas). Mol Biol Cell 12, 1–12.

Manna, D., Albanese, A., Park, W.S., and Cho, W. (2007). Mechanistic basis of differential cellular responses of phosphatidylinositol 3,4-bisphosphate- and phosphatidylinositol 3,4,5-trisphosphate-binding pleckstrin homology domains. J Biol Chem 282, 32093–32105.

Manning, B.D., and Toker, A. (2017). AKT/PKB Signaling: Navigating the Network. Cell 169, 381–405.

Mijanovic, L., and Weber, I. (2022). Adhesion of Dictyostelium Amoebae to Surfaces: A Brief History of Attachments. Front Cell Dev Biol 10.

Naguib, A., Bencze, G., Cho, H., Zheng, W., Tocilj, A., Elkayam, E., Faehnle, C.R., Jaber, N., Pratt, C.P., Chen, M., et al. (2015). PTEN functions by recruitment to cytoplasmic vesicles. Mol Cell 58, 255–268.

Nakatsu, F., Perera, R.M., Lucast, L., Zoncu, R., Domin, J., Gertler, F.B., Toomre, D., and De Camilli, P. (2010). The inositol 5-phosphatase SHIP2 regulates endocytic clathrin-coated pit dynamics. J Cell Biol 190, 307–315.

Norris, D.M., Yang, P., Krycer, J.R., Fazakerley, D.J., James, D.E., and Burchfield, J.G. (2017). An improved Akt reporter reveals intra- and inter-cellular heterogeneity and oscillations in signal transduction. J Cell Sci 130, 2757–2766.

Okoh, M.P., and Vihinen, M. (1999). Pleckstrin homology domains of tec family protein kinases. Biochem Biophys Res Commun 265, 151–157.

Pinilla-Macua, I., Grassart, A., Duvvuri, U., Watkins, S.C., and Sorkin, A. (2017). EGF receptor signaling, phosphorylation, ubiquitylation and endocytosis in tumors in vivo. Elife 6.

Ran, F.A., Hsu, P.D., Wright, J., Agarwala, V., Scott, D.A., and Zhang, F. (2013). Genome engineering using the CRISPR-Cas9 system. Nat Protoc 8, 2281–2308.

Rathinaswamy, M.K., and Burke, J.E. (2020). Class I phosphoinositide 3-kinase (PI3K) regulatory subunits and their roles in signaling and disease. Adv Biol Regul 75, 100657.

Rich, T., Zhao, F., Cruciani, R.A., Cella, D., Manola, J., and Fisch, M.J. (2017). Association of fatigue and depression with circulating levels of proinflammatory cytokines and epidermal growth factor receptor ligands: a correlative study of a placebo-controlled fatigue trial. Cancer Manag Res 9, 1–10.

Riehle, R.D., Cornea, S., and Degterev, A. (2013). Role of phosphatidylinositol 3,4,5-trisphosphate in cell signaling. Adv Exp Med Biol 991, 105–139.

Salamon, R.S., and Backer, J.M. (2013). Phosphatidylinositol-3,4,5-trisphosphate: tool of choice for class I PI 3-kinases. Bioessays 35, 602–611.

Samuels, Y., Diaz, L.A., Jr., Schmidt-Kittler, O., Cummins, J.M., Delong, L., Cheong, I., Rago, C., Huso, D.L., Lengauer, C., Kinzler, K.W., et al. (2005). Mutant PIK3CA promotes cell growth and invasion of human cancer cells. Cancer Cell 7, 561–573.

Samuels, Y., Wang, Z., Bardelli, A., Silliman, N., Ptak, J., Szabo, S., Yan, H., Gazdar, A., Powell, S.M., Riggins, G.J., et al. (2004). High frequency of mutations of the PIK3CA gene in human cancers. Science 304, 554.

Schindelin, J., Arganda-Carreras, I., Frise, E., Kaynig, V., Longair, M., Pietzsch, T., Preibisch, S., Rueden, C., Saalfeld, S., Schmid, B., et al. (2012). Fiji: an open-source platform for biological-image analysis. Nat Methods 9, 676–682.

Shaner, N.C., Lambert, G.G., Chammas, A., Ni, Y., Cranfill, P.J., Baird, M.A., Sell, B.R., Allen, J.R., Day, R.N., Israelsson, M., et al. (2013). A bright monomeric green fluorescent protein derived from Branchiostoma lanceolatum. Nat Methods 10, 407–409.

Singh, P., Dar, M.S., Singh, G., Jamwal, G., Sharma, P.R., Ahmad, M., and Dar, M.J. (2016). Dynamics of GFP-Fusion p110alpha and p110beta Isoforms of PI3K Signaling Pathway in Normal and Cancer Cells. J Cell Biochem 117, 2864–2874.

Stirling, D.R., Swain-Bowden, M.J., Lucas, A.M., Carpenter, A.E., Cimini, B.A., and Goodman, A. (2021). CellProfiler 4: improvements in speed, utility and usability. BMC Bioinformatics 22, 433.

Su, Y., Li, R., Ning, X., Lin, Z., Zhao, X., Zhou, J., Liu, J., Jin, Y., and Yin, Y. (2019). Discovery of 2,4-diarylaminopyrimidine derivatives bearing dithiocarbamate moiety as novel FAK inhibitors with antitumor and anti-angiogenesis activities. Eur J Med Chem 177, 32–46.

Sugiyama, M.G., Fairn, G.D., and Antonescu, C.N. (2019). Akt-ing Up Just About Everywhere: Compartment-Specific Akt Activation and Function in Receptor Tyrosine Kinase Signaling. Front Cell Dev Biol 7, 70.

Sun, H., Lesche, R., Li, D.M., Liliental, J., Zhang, H., Gao, J., Gavrilova, N., Mueller, B., Liu, X., and Wu, H. (1999). PTEN modulates cell cycle progression and cell survival by regulating phosphatidylinositol 3,4,5,-trisphosphate and Akt/protein kinase B signaling pathway. Proc Natl Acad Sci U S A 96, 6199–6204.

Tarantino, N., Tinevez, J.Y., Crowell, E.F., Boisson, B., Henriques, R., Mhlanga, M., Agou, F., Israel, A., and Laplantine, E. (2014). TNF and IL-1 exhibit distinct ubiquitin requirements for inducing NEMO-IKK supramolecular structures. J Cell Biol 204, 231–245.

Thapa, N., Chen, M., Horn, H.T., Choi, S., Wen, T., and Anderson, R.A. (2020). Phosphatidylinositol-3-OH kinase signalling is spatially organized at endosomal compartments by microtubule-associated protein 4. Nat Cell Biol 22, 1357–1370.

Vadas, O., Burke, J.E., Zhang, X., Berndt, A., and Williams, R.L. (2011). Structural basis for activation and inhibition of class I phosphoinositide 3-kinases. Sci Signal 4, re2.

Vanhaesebroeck, B., Perry, M.W.D., Brown, J.R., Andre, F., and Okkenhaug, K. (2021). PI3K inhibitors are finally coming of age. Nat Rev Drug Discov 20, 741–769.

Varnai, P., Rother, K.I., and Balla, T. (1999). Phosphatidylinositol 3-kinase-dependent membrane association of the Bruton’s tyrosine kinase pleckstrin homology domain visualized in single living cells. J Biol Chem 274, 10983–10989.

Vasan, N., Razavi, P., Johnson, J.L., Shao, H., Shah, H., Antoine, A., Ladewig, E., Gorelick, A., Lin, T.Y., Toska, E., et al. (2019). Double PIK3CA mutations in cis increase oncogenicity and sensitivity to PI3Kalpha inhibitors. Science 366, 714–723.

Vega-Lugo, J., da Rocha-Azevedo, B., Dasgupta, A., and Jaqaman, K. (2022). Analysis of conditional colocalization relationships and hierarchies in three-color microscopy images. J Cell Biol 221.

Walpole, G.F.W., Pacheco, J., Chauhan, N., Clark, J., Anderson, K.E., Abbas, Y.M., Brabant-Kirwan, D., Montano-Rendon, F., Liu, Z., Zhu, H., et al. (2022). Kinase-independent synthesis of 3-phosphorylated phosphoinositides by a phosphotransferase. Nat Cell Biol 24, 708–722.

Wang, Q., Pechersky, Y., Sagawa, S., Pan, A.C., and Shaw, D.E. (2019). Structural mechanism for Bruton’s tyrosine kinase activation at the cell membrane. Proc Natl Acad Sci U S A 116, 9390–9399.

Watton, S.J., and Downward, J. (1999). Akt/PKB localisation and 3’ phosphoinositide generation at sites of epithelial cell-matrix and cell-cell interaction. Curr Biol 9, 433–436.

Yamakita, Y., Totsukawa, G., Yamashiro, S., Fry, D., Zhang, X., Hanks, S.K., and Matsumura, F. (1999). Dissociation of FAK/p130(CAS)/c-Src complex during mitosis: role of mitosis-specific serine phosphorylation of FAK. J Cell Biol 144, 315–324.

Yip, S.C., El-Sibai, M., Hill, K.M., Wu, H., Fu, Z., Condeelis, J.S., and Backer, J.M. (2004). Over-expression of the p110beta but not p110alpha isoform of PI 3-kinase inhibits motility in breast cancer cells. Cell Motil Cytoskeleton 59, 180–188.

Zhan, H., Bhattacharya, S., Cai, H., Iglesias, P.A., Huang, C.H., and Devreotes, P.N. (2020). An Excitable Ras/PI3K/ERK Signaling Network Controls Migration and Oncogenic Transformation in Epithelial Cells. Dev Cell 54, 608–623 e605.

Zhao, X., and Guan, J.L. (2011). Focal adhesion kinase and its signaling pathways in cell migration and angiogenesis. Adv Drug Deliver Rev 63, 610–615.

