## Supplemental information for "Spatial organization of PI3K-PI(3,4,5)P3-AKT signaling by focal adhesions"

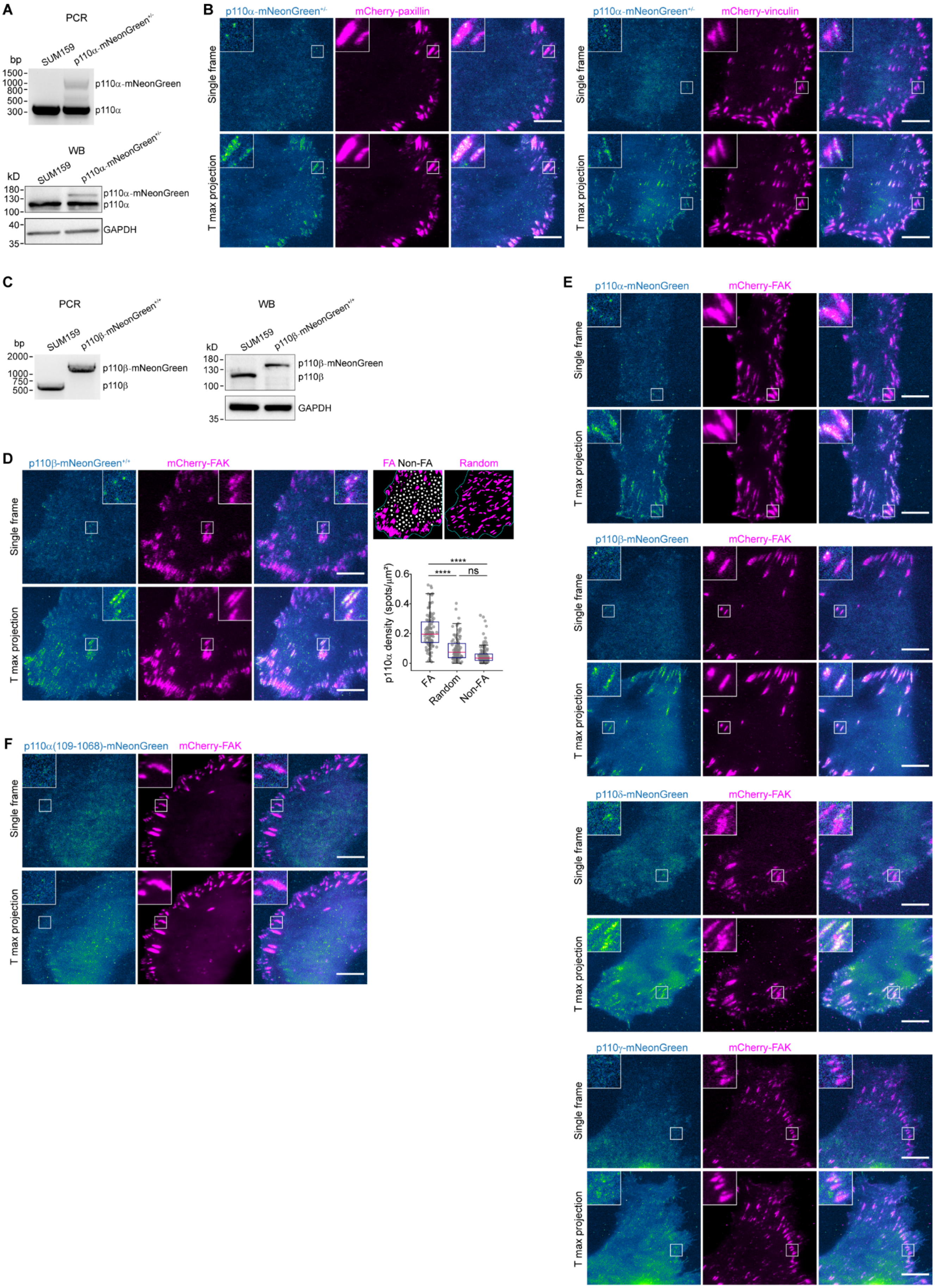

A
PCR
SUM159
p110α-mNeonGreen+/-
bp
1500
1000
800
500
300
p110α-mNeonGreen
p110α
WB
kD
180
130
100
40
35
GAPDH
B
p110α-mNeonGreen+/-
mCherry-paxillin
mCherry-vinculin
Single frame
T max projection
C
p110β-mNeonGreen+/+
2000
750
p110β-mNeonGreen
p110β
D
mCherry-FAK
FA Non-FA
Random
p110α density (spots/μm²)
FA
Random
Non-FA
****
ns
E
p110α-mNeonGreen
p110β-mNeonGreen
p110δ-mNeonGreen
p110γ-mNeonGreen
F
p110α(109-1068)-mNeonGreen

**Figure S1. The catalytic subunits of class IA PI3K are dynamically recruited to FAs, related to Figure 1**

(A) Single allelic integration of the mNeonGreen sequence into the *PIK3CA* genomic locus (with wild-type H1047) in clonal gene-edited p110α-mNeonGreen$^{+/-}$ SUM159 cells, as confirmed by genomic PCR analysis (top) and western blot analysis with antibodies for p110α and GAPDH (bottom).

(B) p110α-mNeonGreen$^{+/-}$ cells transiently expressing mCherry-paxillin or mCherry-vinculin were imaged at 0.2-s intervals for 601 frames. Images of a single frame and maximum-intensity projection of a representative time series are shown.

(C) Dual allelic integration of the mNeonGreen sequence into the *PIK3CB* genomic locus in clonal gene-edited p110β-mNeonGreen$^{+/+}$ SUM159 cells, as confirmed by genomic PCR analysis (left) and western blot analysis with antibodies for p110β and GAPDH (right).

(D) Left: p110β-mNeonGreen$^{+/+}$ cells transiently expressing mCherry-FAK were imaged at 0.2-s intervals for 601 frames. Images of a single frame and maximum-intensity projection of a representative time series are shown. Right: Images showing FA, Non-FA, and Random regions of the cell. Box plots show the relative density of p110β-mNeonGreen molecules recruited to different regions (box: median with the 25th and 75th percentiles; whiskers: 1.5-fold the interquartile range).

(E) Cells transiently expressing mCherry-FAK with p110α-mNeonGreen, p110β-mNeonGreen, p110δ-mNeonGreen, or p110γ-mNeonGreen were imaged at 0.2-s intervals for 601 frames. Images of a single frame and maximum-intensity projection of a representative time series are shown.

(F) Cells transiently expressing mCherry-FAK and p110α(109-1068)-mNeonGreen were imaged at 0.2-s intervals for 601 frames. Images of a single frame and maximum-intensity projection of a representative time series are shown.

Cells were imaged at the bottom surface by TIRF microscopy in (B) and (D-F). Statistical analysis was performed using ordinary one-way ANOVA with Tukey's multiple comparisons test; $****P < 0.0001$; ns, not significant. Scale bars, 10 μm.

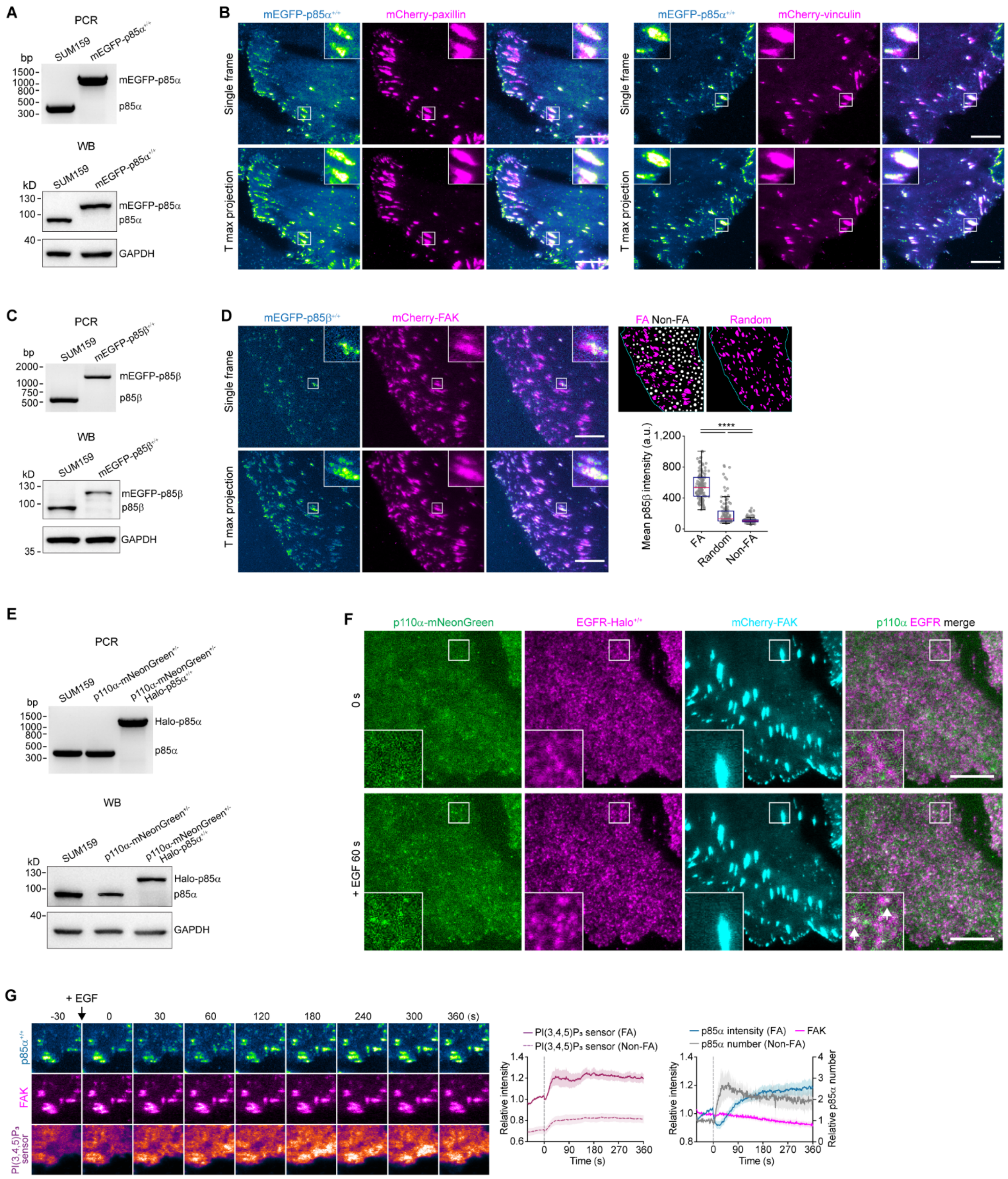

A
PCR
bp
SUM159
mEGFP-p85α+/+
mEGFP-p85α
p85α
WB
kD
GAPDH
B
mEGFP-p85α+/+
mCherry-paxillin
mCherry-vinculin
Single frame
T max projection
C
mEGFP-p85β+/+
mEGFP-p85β
p85β
D
mCherry-FAK
FA Non-FA
Random
Mean p85β intensity (a.u.)
****
FA
Random
Non-FA
E
p110α-mNeonGreen+/-
p110α-mNeonGreen+/- Halo-p85α+/+
Halo-p85α
F
p110α-mNeonGreen
EGFR-Halo+/+
mCherry-FAK
p110α EGFR merge
0 s
+ EGF 60 s
G
+ EGF
-30 0 30 60 120 180 240 300 360 (s)
p85α+/+
FAK
PI(3,4,5)P3 sensor
PI(3,4,5)P3 sensor (FA)
PI(3,4,5)P3 sensor (Non-FA)
Relative intensity
Time (s)
p85α intensity (FA)
FAK
p85α number (Non-FA)
Relative p85α number

**Figure S2. The regulatory subunits of class IA PI3K are dynamically recruited to FAs, related to Figure 1**

(A) Biallelic integration of the mEGFP sequence into the *PIK3R1* genomic locus of SUM159 cells to generate the gene-edited clonal cell line mEGFP-p85α$^{+/+}$, as confirmed by genomic PCR analysis (top) and western blot analysis with antibodies for p85α and GAPDH (bottom).

(B) mEGFP-p85α$^{+/+}$ cells transiently expressing mCherry-paxillin (left) or mCherry-vinculin (right) were imaged at 0.2-s intervals for 601 frames. Images of a single frame and maximum-intensity projection of a representative time series are shown.

(C) Biallelic integration of the mEGFP sequence into the *PIK3R2* genomic locus of SUM159 cells to generate the gene-edited clonal cell line mEGFP-p85β$^{+/+}$, as confirmed by genomic PCR analysis (top) and western blot analysis with antibodies for p85β and GAPDH (bottom).

(D) mEGFP-p85β$^{+/+}$ cells transiently expressing mCherry-FAK were imaged at 0.2-s intervals for 601 frames. Images of a single frame and maximum-intensity projection of a representative time series are shown. Right: Images showing FA, Non-FA, and Random regions of the cell. Box plots show the relative intensity of mEGFP-p85β at different regions.

(E) Biallelic integration of the HaloTag sequence into the *PIK3R1* genomic locus of SUM159 p110α-mNeonGreen$^{+/-}$ cells to generate the gene-edited clonal cell line p110α-mNeonGreen$^{+/-}$ Halo-p85α$^{+/+}$, as confirmed by genomic PCR analysis (top) and western blot analysis with antibodies for p85α and GAPDH (bottom).

(F) Gene-edited EGFR-Halo$^{+/+}$ (labeled with $JFX_{650}$-HaloTag ligand) SUM159 cells transiently expressing p110α-mNeonGreen were imaged at 10-s intervals, with EGF added (set as 0 s) during continuous imaging. Representative images showing the recruitment of p110α-mNeonGreen to the plasma membrane and colocalization with EGFR (arrows) after EGF treatment.

(G) mEGFP-p85α$^{+/+}$ cells transiently expressing mCherry-FAK and the PI(3,4,5)$P_3$ sensor Halo-2xBtk(2-166) (labeled with $JFX_{650}$-HaloTag ligand) were imaged at 10-s intervals, with EGF added (set as 0 s) during continuous imaging. Left: Montage showing images at the indicated frames of a representative time series. Middle: Plots showing the relative intensity of the PI(3,4,5)$P_3$ sensor in FA and Non-FA regions (normalized by the mean intensity in FA regions before EGF treatment). Right: Plots showing the relative intensity of mEGFP-p85α and mCherry-FAK in FAs, and the relative number of mEGFP-p85α in Non-FA regions (mean ± SEM; n = 5 cells).

Cells were imaged at the bottom surface by TIRF microscopy in (B), (D), (F), and (G). Statistical analysis was performed using ordinary one-way ANOVA with Tukey's multiple comparisons test; ****$P < 0.0001$. Scale bars, 10 μm.

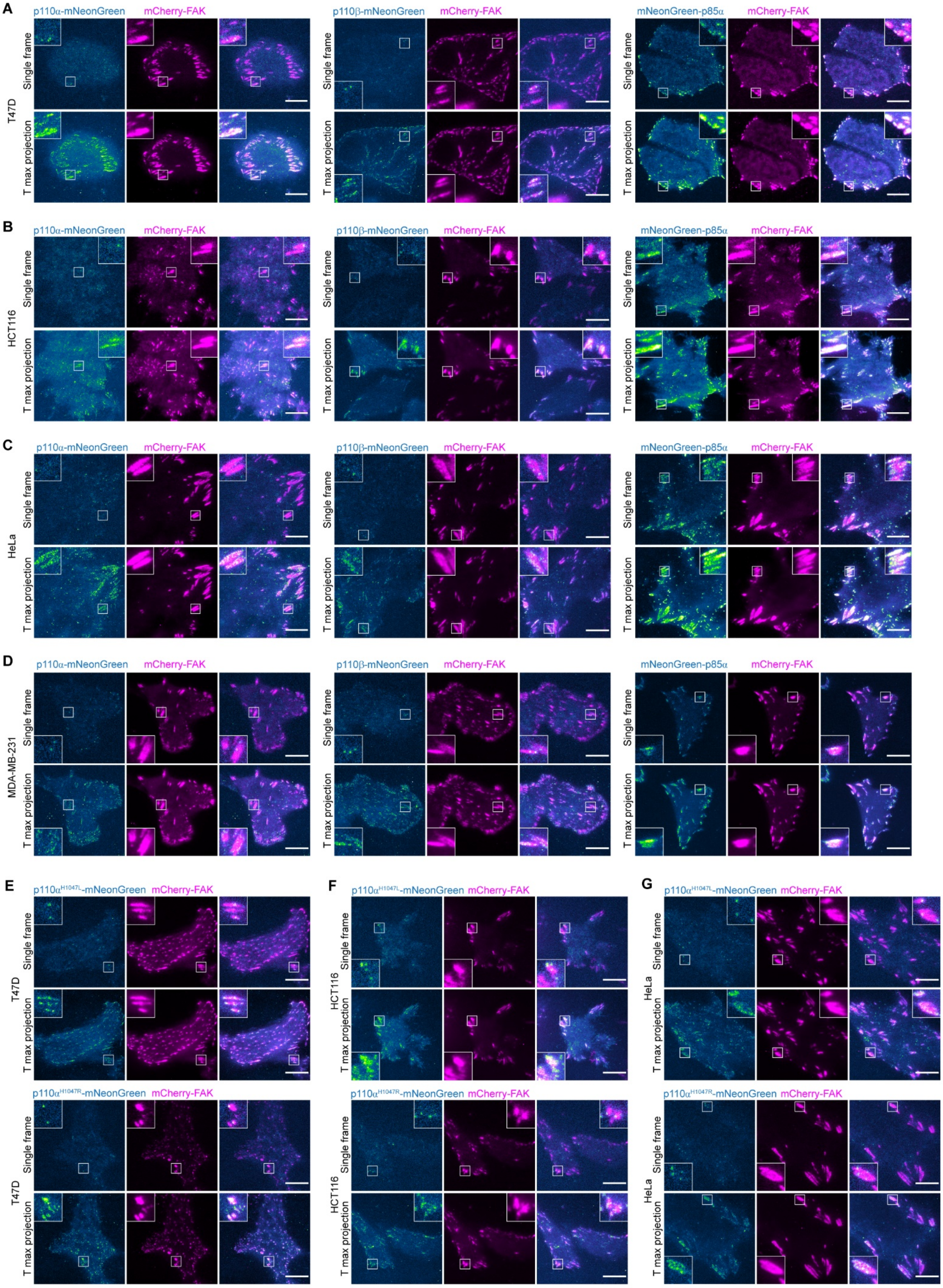
A
p110α-mNeonGreen
mCherry-FAK
p110β-mNeonGreen
mCherry-FAK
mNeonGreen-p85α
mCherry-FAK
T47D
Single frame
T max projection
B
HCT116
C
HeLa
D
MDA-MB-231
E
p110α^H1047L-mNeonGreen
mCherry-FAK
T47D
p110α^H1047R-mNeonGreen
F
HCT116
G
HeLa

**Figure S3. The catalytic and regulatory subunits of class IA PI3K are dynamically recruited to FAs in different cancer cell lines, related to Figure 1**

(A-D), Bottom surfaces of T47D (A), HCT116 (B), HeLa (C), or MDA-MB-231 (D) cells transiently expressing mCherry-FAK with p110α-mNeonGreen (left), p110β-mNeonGreen (middle), or mNeonGreen-p85α (right). Imaging was performed by TIRF microscopy. Cells in the left and middle panels were imaged at 0.2-s intervals for 601 frames and cells in the right panels were imaged at 2-s intervals for 151 frames. Images of a single frame and maximum-intensity projection of a representative time series are shown.

(E-G) Bottom surfaces of T47D (E), HCT116 (F), or HeLa (G) cells transiently expressing mCherry-FAK with p110α$^{H1047L}$-mNeonGreen (top) or p110α$^{H1047R}$-mNeonGreen (bottom) were imaged at 0.2-s intervals for 601 frames by TIRF microscopy. Images of a single frame and maximum-intensity projection of a representative time series are shown. Scale bars, 10 μm.

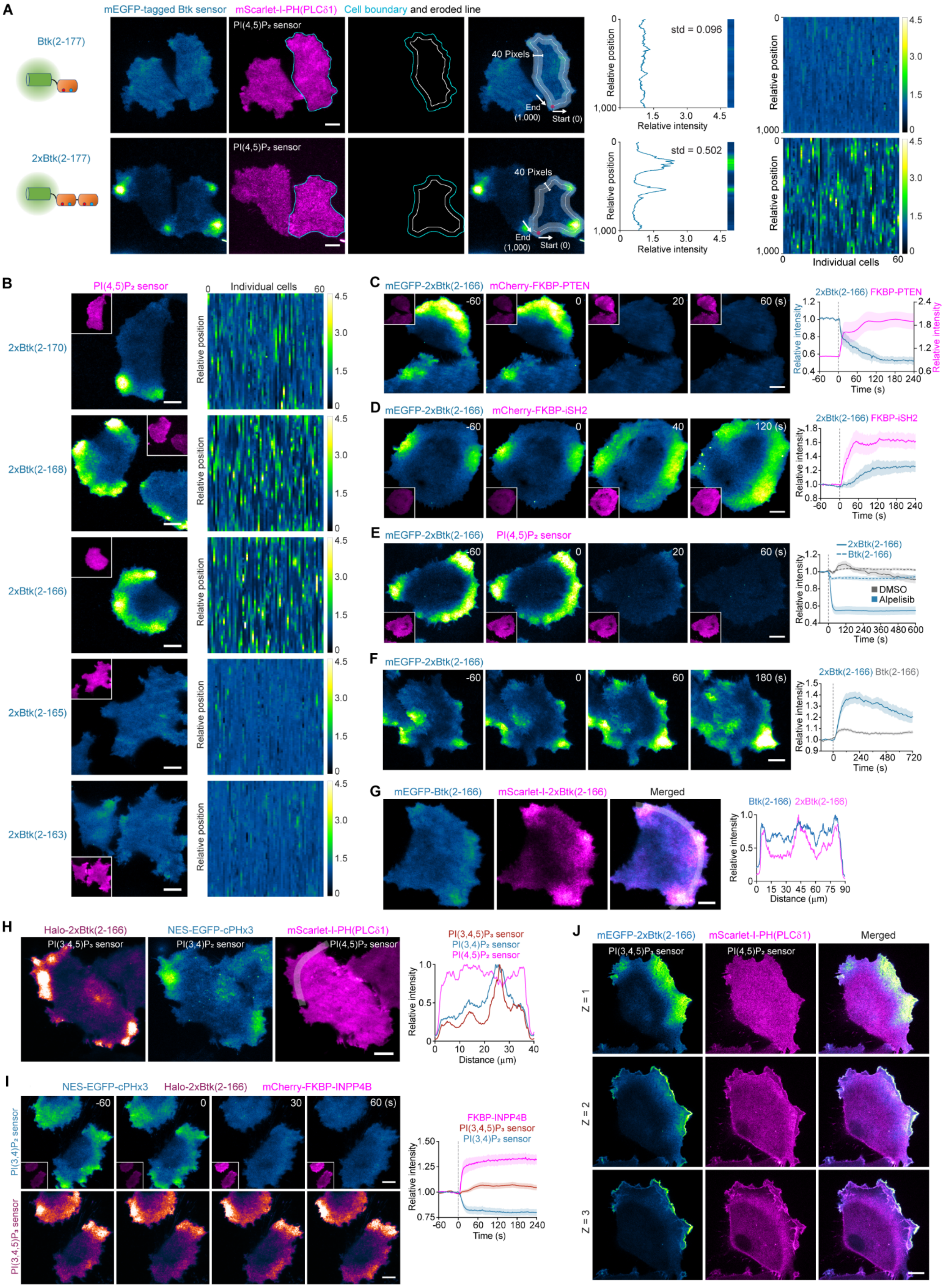
A
mEGFP-tagged Btk sensor
mScarlet-I-PH(PLCδ1)
Cell boundary and eroded line
Btk(2-177)
2xBtk(2-177)
PI(4,5)P₂ sensor
40 Pixels
End (1,000)
Start (0)
std = 0.096
std = 0.502
Relative position
Relative intensity
Individual cells
B
PI(4,5)P₂ sensor
2xBtk(2-170)
2xBtk(2-168)
2xBtk(2-166)
2xBtk(2-165)
2xBtk(2-163)
C
mEGFP-2xBtk(2-166)
mCherry-FKBP-PTEN
2xBtk(2-166) FKBP-PTEN
Time (s)
D
mCherry-FKBP-iSH2
2xBtk(2-166) FKBP-iSH2
E
2xBtk(2-166)
Btk(2-166)
DMSO
Alpelisib
F
2xBtk(2-166) Btk(2-166)
G
mEGFP-Btk(2-166)
mScarlet-I-2xBtk(2-166)
Merged
Btk(2-166) 2xBtk(2-166)
Distance (μm)
H
Halo-2xBtk(2-166)
NES-EGFP-cPHx3
PI(3,4,5)P₃ sensor
PI(3,4)P₂ sensor
PI(4,5)P₂ sensor
I
mCherry-FKBP-INPP4B
FKBP-INPP4B
J
mEGFP-2xBtk(2-166)
Z = 1
Z = 2
Z = 3

**Figure S4. The 2xBtk(2-166) domain exhibits high selectivity and sensitivity in binding PI(3,4,5)$P_3$ at the plasma membrane, related to Figure 2**

(A) Cells transiently expressing the PI(4,5)$P_2$ sensor mScarlet-I-PH(PLCδ1) together with low levels of mEGFP-Btk(2-177) or mEGFP-2xBtk(2-177) were imaged by TIRF microscopy. Schematic domain structures of the Btk sensors and the distribution of Btk sensors with mScarlet-I-PH(PLCδ1) at the plasma membrane are shown in the left panels. To analyze the distribution of Btk sensors around the cell periphery, the cell outline was segmented based on the fluorescence of mScarlet-I-PH(PLCδ1) (cyan lines) and eroded for 20 pixels (white lines). The fluorescence intensity of Btk sensors along the eroded line (width of 40 pixels, starting from the bottom position of the cell) was measured. Plots show the measured intensity profile along the line (starting position set as 0, end position set as 1,000) normalized by the average intensity of the measured region. The standard deviation (std) of each plot is shown. Each plot is also presented as a color-coded bar. The rightmost panels show the stacked color-coded bar graphs from multiple cells (intensity profile heatmap, n = 60 cells each).

(B) Left: Representative images of cells transiently expressing the PI(4,5)$P_2$ sensor mScarlet-I-PH(PLCδ1) (inserts) together with the indicated mEGFP-tagged Btk sensors. Right: Intensity profile heatmaps of mEGFP-tagged Btk sensors (n = 60 cells each).

(C) Cells co-expressing LYN11-FRB-ECFP, mCherry-FKBP-PTEN, and the PI(3,4,5)$P_3$ sensor mEGFP-2xBtk(2-166) were imaged at 2-s intervals. Rapamycin was added (set as 0 s) during continuous imaging to trigger acute depletion of PI(3,4,5)$P_3$ by recruiting mCherry-FKBP-PTEN (inserts) from the cytosol to the plasma membrane. Plots show the relative intensity of mCherry-FKBP-PTEN and PI(3,4,5)$P_3$ sensor before and after rapamycin treatment (n = 9 cells).

(D) Cells co-expressing LYN11-FRB-ECFP, mCherry-FKBP-iSH2, and the PI(3,4,5)$P_3$ sensor mEGFP-2xBtk(2-166) were imaged at 2-s intervals. Rapamycin was added (set as 0 s) during continuous imaging to trigger acute production of PI(3,4,5)$P_3$ by recruiting mCherry-FKBP-iSH2 (inserts) from the cytosol to the plasma membrane. Plots show the relative intensity of mCherry-FKBP-iSH2 and PI(3,4,5)$P_3$ sensor before and after rapamycin treatment (n = 6 cells).

(E) Left: Cells co-expressing the PI(4,5)$P_2$ sensor mScarlet-I-PH(PLCδ1) with mEGFP-2xBtk(2-166) or mEGFP-Btk(2-166) were imaged at 10-s intervals. The p110α inhibitor alpelisib was added (set as 0 s) during continuous imaging to trigger acute depletion of PI(3,4,5)$P_3$. Right: Plots showing the relative intensity of mEGFP-2xBtk(2-166) and mEGFP-Btk(2-166) in the alpelisib (n = 12 and 14 cells) or DMSO (n = 14 and 12 cells) treated cells.

(F) Cells expressing mEGFP-2xBtk(2-166) or mEGFP-Btk(2-166) were imaged at 10-s intervals. EGF was added (set as 0 s) during continuous imaging to trigger acute generation of PI(3,4,5)$P_3$. Plots show the relative intensity of mEGFP-2xBtk(2-166) and mEGFP-Btk(2-166) in the cells treated with EGF (n = 28 and 25 cells).

(G) Left: Representative images of cells transiently expressing mEGFP-Btk(2-166) and mScarlet-I-2xBtk(2-166). Right: Plots showing the relative intensity profile along the line.

(H) Left: Representative images of cells transiently expressing the PI(3,4,5)$P_3$ sensor Halo-2xBtk(2-166) (labeled with $JFX_{650}$-HaloTag ligand), the PI(4,5)$P_2$ sensor mScarlet-I-PH(PLCδ1), and the PI(3,4)$P_2$ sensor NES-EGFP-cPHx3. Right: Plots showing the relative intensity profile along the line on the PI(4,5)$P_2$ sensor.

(I) Cells co-expressing LYN11-FRB-ECFP, mCherry-FKBP-INPP4B, the PI(3,4)$P_2$ sensor NES-EGFP-cPHx3, and the PI(3,4,5)$P_3$ sensor Halo-2xBtk(2-166) (labeled with $JFX_{650}$-HaloTag ligand) were imaged at 2-s intervals. Rapamycin was added (set as 0 s) during continuous imaging to trigger acute depletion of plasma membrane PI(3,4)$P_2$ by recruiting mCherry-FKBP-INPP4B (inserts) from the cytosol to the plasma membrane. Plots show the relative fluorescent intensity of mCherry-FKBP-INPP4B, $PI(3,4)P_2$ sensor, and $PI(3,4,5)P_3$ sensor before and after rapamycin treatment (mean ± SEM; n = 16 cells).

(J) SUM159 cells transiently expressing the $PI(4,5)P_2$ sensor mScarlet-I-PH(PLCδ1) with the $PI(3,4,5)P_3$ sensor mEGFP-2xBtk(2-166) were imaged by spinning-disk confocal microscopy. Representative images show the distribution of $PI(3,4,5)P_3$ and $PI(4,5)P_2$ sensors at optical sections (spaced 0.5 μm) from the bottom surface (Z = 1) to the middle of the cells.

Cells were imaged at the bottom surface by TIRF microscopy in (A-I). Data are shown as mean ± SEM in (C-F) and (I). Scale bars, 10 μm.

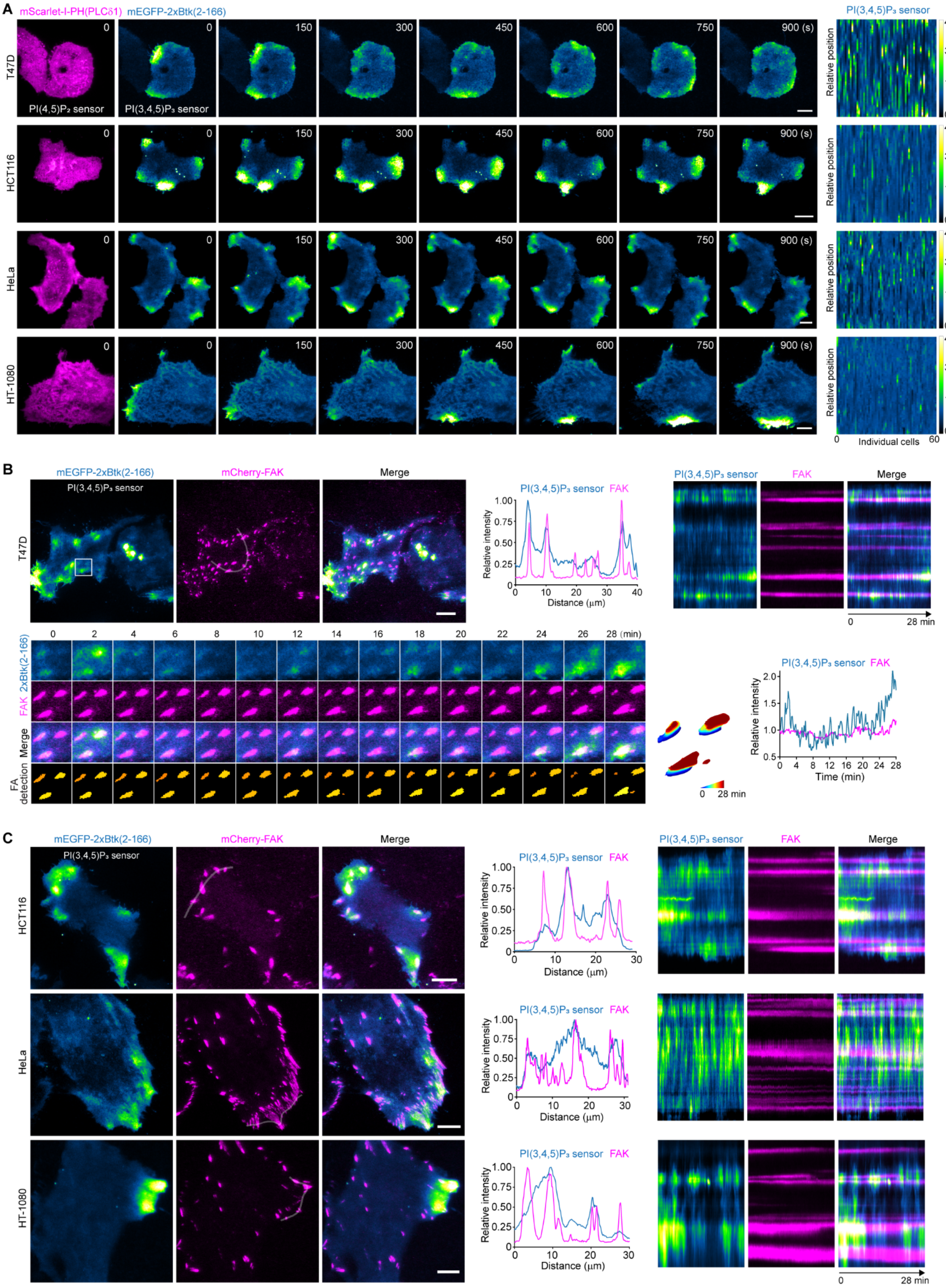
A
mScarlet-I-PH(PLCδ1) mEGFP-2xBtk(2-166)
PI(4,5)P₂ sensor
PI(3,4,5)P₃ sensor
T47D
HCT116
HeLa
HT-1080
0 150 300 450 600 750 900 (s)
PI(3,4,5)P₃ sensor
Relative position
Individual cells
B
mEGFP-2xBtk(2-166)
mCherry-FAK
Merge
Relative intensity
Distance (μm)
FAK
0 28 min
2xBtk(2-166)
FA detection
28 (min)
Time (min)
C

**Figure S5. Dynamic local enrichment of PI(3,4,5)$P_3$ around FAs at the plasma membrane of different cancer cell lines, related to Figure 2**

(A) Bottom surfaces of T47D, HCT116, HeLa, or HT-1080 cells transiently expressing the PI(3,4,5)$P_3$ sensor mEGFP-2xBtk(2-166) and the PI(4,5)$P_2$ sensor mScarlet-I-PH(PLCδ1) were imaged at 10-s intervals by TIRF microscopy. Left: Montages showing images of PI(3,4,5)$P_3$ and PI(4,5)$P_2$ sensors at the indicated times of representative time series. Right: Intensity profile heatmap of the PI(3,4,5)$P_3$ sensor (n = 60 cells).

(B) Bottom surfaces of T47D cells transiently expressing the PI(3,4,5)$P_3$ sensor mEGFP-2xBtk(2-166) with mCherry-FAK were imaged at 10-s intervals. From left to right in the top panels: Images of PI(3,4,5)$P_3$ sensor and mCherry-FAK in a single frame of a representative time series; plots showing the relative intensity profile along the line on mCherry-FAK; kymographs generated along the line showing dynamic oscillation of PI(3,4,5)$P_3$ sensor around FAs. Bottom: Montage showing images of PI(3,4,5)$P_3$ sensor and mCherry-FAK at the indicated times of the time series, with detected FAs plotted at the bottom; plots showing the relative intensity of PI(3,4,5)$P_3$ sensor and mCherry-FAK over time.

(C) Bottom surfaces of HCT116, HeLa, or HT-1080 cells transiently expressing the PI(3,4,5)$P_3$ sensor mEGFP-2xBtk(2-166) and mCherry-FAK were imaged at 10-s intervals. Left: Images of PI(3,4,5)$P_3$ sensor and mCherry-FAK on a single frame of representative time series. Middle: Plots showing the relative intensity profile along the lines on mCherry-FAK. Right: Kymographs generated along the lines showing dynamic oscillation of PI(3,4,5)$P_3$ sensor around FAs.

Cells were imaged at the bottom surface by TIRF microscopy in (A-C). Scale bars, 10 μm.

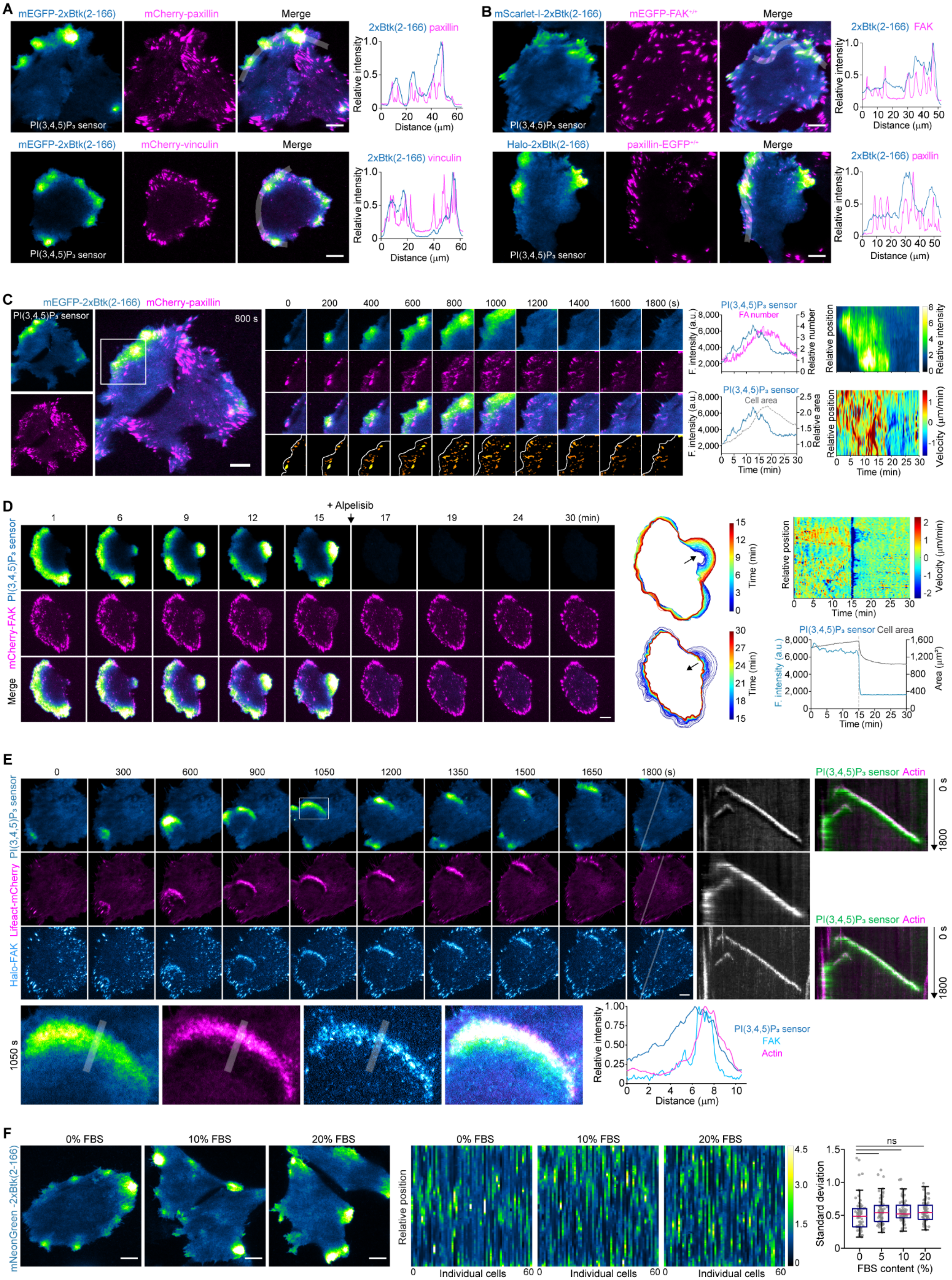

A
mEGFP-2xBtk(2-166)
mCherry-paxillin
Merge
PI(3,4,5)P₃ sensor
2xBtk(2-166) paxillin
Relative intensity
Distance (μm)
mCherry-vinculin
2xBtk(2-166) vinculin
B
mScarlet-I-2xBtk(2-166)
mEGFP-FAK+/+
2xBtk(2-166) FAK
Halo-2xBtk(2-166)
paxillin-EGFP+/+
2xBtk(2-166) paxillin
C
mEGFP-2xBtk(2-166) mCherry-paxillin
800 s
0 200 400 600 800 1000 1200 1400 1600 1800 (s)
PI(3,4,5)P₃ sensor
FA number
F. intensity (a.u.)
Relative number
Relative position
Relative intensity
Cell area
Relative area
Velocity (μm/min)
Time (min)
D
+ Alpelisib
1 6 9 12 15 17 19 24 30 (min)
PI(3,4,5)P₃ sensor
mCherry-FAK
Merge
Time (min)
Relative position
Velocity (μm/min)
PI(3,4,5)P₃ sensor Cell area
F. intensity (a.u.)
Area (μm²)
Time (min)
E
0 300 600 900 1050 1200 1350 1500 1650 1800 (s)
PI(3,4,5)P₃ sensor
Lifeact-mCherry
Halo-FAK
PI(3,4,5)P₃ sensor Actin
0 s
1800
1050 s
Relative intensity
Distance (μm)
PI(3,4,5)P₃ sensor
FAK
Actin
F
mNeonGreen-2xBtk(2-166)
0% FBS
10% FBS
20% FBS
Relative position
Individual cells
ns
Standard deviation
FBS content (%)

**Figure S6. Dynamic local enrichment of PI(3,4,5)$P_3$ around FAs at the plasma membrane of SUM159 cells, related to Figure 2**

(A) Representative images of SUM159 cells transiently expressing the PI(3,4,5)$P_3$ sensor mEGFP-2xBtk(2-166) with mCherry-paxillin (top) or mCherry-vinculin (bottom). Plots show the relative intensity profile along the line on the merged image.

(B) Representative images of SUM159 cells gene-edited for mEGFP-$FAK^{+/+}$ and transiently expressing the PI(3,4,5)$P_3$ sensor mScarlet-I-2xBtk(2-166) (top), or SUM159 cells gene-edited for paxillin-$EGFP^{+/+}$ and transiently expressing the PI(3,4,5)$P_3$ sensor Halo-2xBtk(2-166) (labeled with $JFX_{650}$-HaloTag ligand; bottom). Plots show the relative intensity profile along the line on the merged image.

(C) Cells transiently expressing the PI(3,4,5)$P_3$ sensor mEGFP-2xBtk(2-166) and mCherry-paxillin were imaged at 10-s intervals. Left: Images of PI(3,4,5)$P_3$ sensor and mCherry-paxillin from a single frame of a representative time series. Middle: Montage showing selected frames of the boxed regions; plots showing the intensity of PI(3,4,5)$P_3$ sensor around the leading edge along with the relative numbers of FAs (top) or the area (bottom) of the growing protrusions over time. Right: Heatmap showing the relative intensity of PI(3,4,5)$P_3$ sensor (top) and the cell boundary extension/retraction velocity (bottom) along the leading edge over time.

(D) Cells transiently expressing the PI(3,4,5)$P_3$ sensor mEGFP-2xBtk(2-166) and mCherry-FAK were imaged at 10-s intervals, with alpelisib added at 15 min during continuous imaging. Left: Montage showing images of the indicated frames of the cell before and after alpelisib treatment in a representative time series. Middle: Overlays of the cell outlines between successive frames of the cell before (0-15 min) and after alpelisib treatment (15-30 min). Right top: Cell boundary extension/retraction velocity along the leading edge over time. Right bottom: Plots showing the intensity of PI(3,4,5)$P_3$ sensor and cell area over time.

(E) SUM159 cells transiently expressing the PI(3,4,5)$P_3$ sensor mEGFP-2xBtk(2-166), Halo-FAK (labeled with $JFX_{650}$-HaloTag ligand), and F-actin marker Lifeact-mCherry were imaged at 30-s intervals by TIRF microscopy. Top: Representative montage showing the distribution of PI(3,4,5)$P_3$ sensor, Halo-FAK, and Lifeact-mCherry at the plasma membrane of the cell. Kymographs generated along the line show the distribution of PI(3,4,5)$P_3$ sensor and FAK waves along with propagation of the F-actin wave. Bottom: Magnification of the boxed region on frame 1050 s. Plots show the relative intensity profile along the lines on the magnified images.

(F) Cells stably expressing the PI(3,4,5)$P_3$ sensor mNeonGreen-2xBtk(2-166) were cultured overnight in DMEM/F12 containing 0%, 5%, 10%, or 20% of FBS. Left and middle: Representative image and intensity profile heatmap (n = 60 cells each) of PI(3,4,5)$P_3$ sensor at the plasma membrane of cells in different culture conditions. Right: Box plots showing the standard deviation of PI(3,4,5)$P_3$ sensor distribution at the plasma membrane in each culture condition (box: median with the 25th and 75th percentiles; whiskers: 1.5-fold the interquartile range; n = 63, 71, 70, and 66 cells). Statistical analysis was performed using ordinary one-way ANOVA with Tukey's multiple comparisons test; ns, not significant.

Cells were imaged at the bottom surface by TIRF microscopy in (A-F). Scale bars, 10 μm.

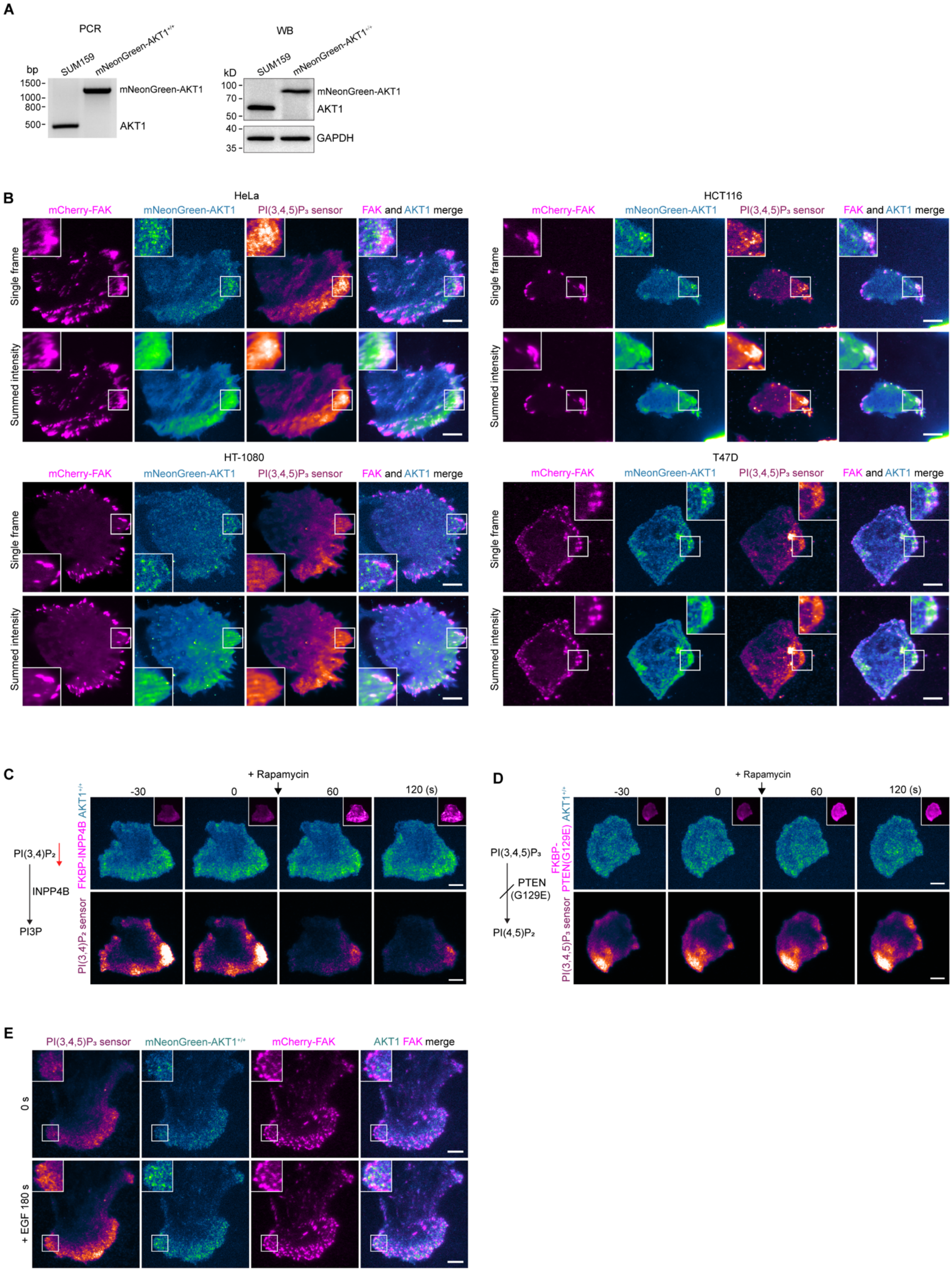
A
PCR
bp
SUM159
mNeonGreen-AKT1+/+
1500-
1000-
800-
500-
mNeonGreen-AKT1
AKT1
WB
kD
SUM159
mNeonGreen-AKT1+/+
100-
70-
50-
40-
35-
mNeonGreen-AKT1
AKT1
GAPDH
B
HeLa
HCT116
HT-1080
T47D
mCherry-FAK
mNeonGreen-AKT1
PI(3,4,5)P3 sensor
FAK and AKT1 merge
Single frame
Summed intensity
C
+ Rapamycin
-30
0
60
120 (s)
PI(3,4)P2
INPP4B
PI3P
FKBP-INPP4B AKT1+/+
PI(3,4)P2 sensor
D
+ Rapamycin
-30
0
60
120 (s)
PI(3,4,5)P3
PTEN (G129E)
PI(4,5)P2
FKBP-PTEN(G129E) AKT1+/+
PI(3,4,5)P3 sensor
E
PI(3,4,5)P3 sensor
mNeonGreen-AKT1+/+
mCherry-FAK
AKT1 FAK merge
0 s
+ EGF 180 s

**Figure S7. Compartmentalized recruitment of AKT1 around FAs in different cancer cell lines, related to Figure 3**

(A) Biallelic integration of the mNeonGreen sequence into the *AKT1* genomic locus of SUM159 cells to generate the gene-edited clonal cell line mNeonGreen-AKT1$^{+/+}$, as confirmed by genomic PCR analysis and western blot analysis with antibodies for AKT1 and GAPDH.

(B) Bottom surfaces of HeLa, HCT116, HT-1080, or T47D cells transiently expressing mNeonGreen-AKT1, mCherry-FAK and the PI(3,4,5)$P_3$ sensor Halo-2xBtk(2-166) (labeled with $JFX_{650}$-HaloTag ligand) were imaged at 0.3-s intervals. Shown are images of a single frame and summed-intensity projection of a representative time series (121 frames).

(C) mNeonGreen-AKT1$^{+/+}$ SUM159 cells transiently co-expressing the PI(3,4)$P_2$ sensor NES-Halo-cPHx3 (labeled with $JFX_{650}$-HaloTag ligand), LYN11-FRB-ECFP and mCherry-FKBP-INPP4B were imaged at 5-s intervals. Rapamycin was added (set as 0 s) during continuous imaging. Top: Montage showing the removal of PI(3,4)$P_2$ sensor but not mNeonGreen-AKT1 from the plasma membrane upon recruitment of mCherry-FKBP-INPP4B (inserts) from the cytosol to the plasma membrane.

(D) mNeonGreen-AKT1$^{+/+}$ SUM159 cells transiently co-expressing the PI(3,4,5)$P_3$ sensor Halo-2xBtk(2-166) (labeled with $JFX_{650}$-HaloTag ligand), LYN11-FRB-ECFP and mCherry-FKBP-PTEN(G129E) were imaged at 5-s intervals. Rapamycin was added (set as 0 s) during continuous imaging. Top: Montage showing the unaffected recruitment of mNeonGreen-AKT1 and PI(3,4,5)$P_3$ sensor upon recruitment of mCherry-FKBP-PTEN(G129E) (inserts) from the cytosol to the plasma membrane.

(E) mNeonGreen-AKT1$^{+/+}$ SUM159 cells transiently expressing mCherry-FAK and the PI(3,4,5)$P_3$ sensor Halo-2xBtk(2-166) (labeled with $JFX_{650}$-HaloTag ligand) were imaged at 5-s intervals. EGF (10 ng/mL) was added (set as 0 s) during continuous imaging. Images at 0 s and 180 s after EGF treatment from a representative time series are shown.

Cells were imaged at the bottom surface by TIRF microscopy in (B-E). Scale bars, 10 μm.

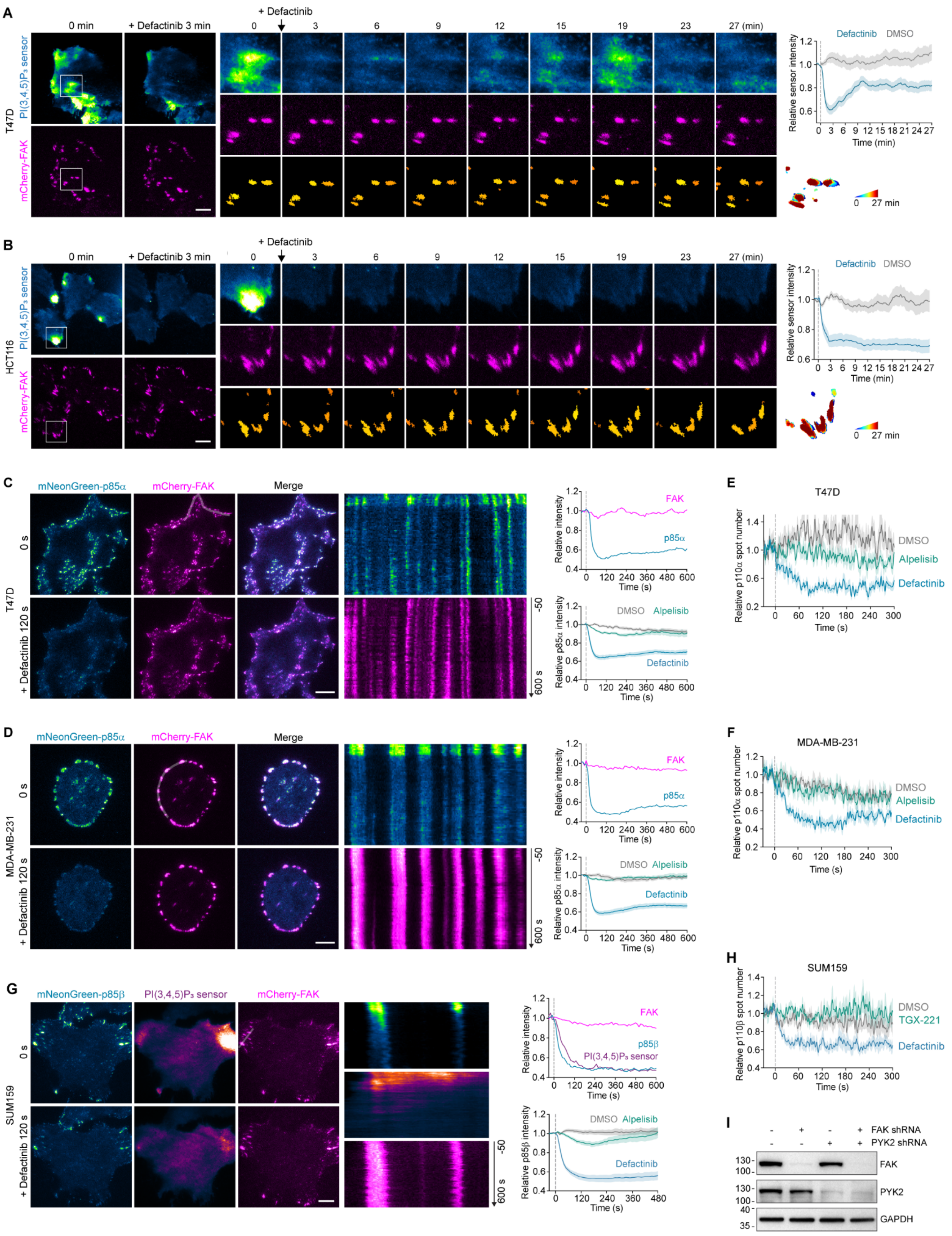
A
0 min
+ Defactinib 3 min
+ Defactinib
0
3
6
9
12
15
19
23
27 (min)
T47D
PI(3,4,5)P₃ sensor
mCherry-FAK
Defactinib
DMSO
Relative sensor intensity
Time (min)
0 27 min
B
HCT116
C
mNeonGreen-p85α
mCherry-FAK
Merge
0 s
+ Defactinib 120 s
Relative intensity
FAK
p85α
Time (s)
Relative p85α intensity
DMSO
Alpelisib
Defactinib
-50
600 s
E
T47D
Relative p110α spot number
D
MDA-MB-231
F
G
mNeonGreen-p85β
SUM159
p85β
Relative p85β intensity
H
Relative p110β spot number
TGX-221
I
FAK shRNA
PYK2 shRNA
FAK
PYK2
GAPDH

**Figure S8. Regulation of PI(3,4,5)$P_3$ generation and class IA PI3K recruitment by the activated FAK in different cell lines, related to Figure 4**

(A and B) Bottom surfaces of T47D (A) or HCT116 (B) cells transiently expressing the PI(3,4,5)$P_3$ sensor mEGFP-2xBtk(2-166) and mCherry-FAK were imaged at 10-s intervals, with defactinib added (set as 0 min) during continuous imaging. Left: Images at 0 min and 3 min after defactinib treatment in a representative time series. Middle: Montages showing the indicated frames of the boxed regions, with detected FAs plotted at the bottom. Right: Plots showing the relative intensity of PI(3,4,5)$P_3$ sensor on FAs of defactinib- or DMSO-treated T47D cells (n = 18 and 16 cells) or HCT116 cells (n = 19 and 12 cells).

(C and D) T47D (C) or MDA-MB-231 (D) cells transiently expressing mNeonGreen-p85α and mCherry-FAK were imaged at 10-s intervals, with DMSO, defactinib, or alpelisib added (set as 0 s) during continuous imaging. Left: Images at 0 s and 120 s after defactinib treatment in a representative time series. Middle: Kymographs generated along the line on mCherry-FAK showing dynamics of mNeonGreen-p85α and mCherry-FAK. Right top: Plots showing the relative intensity of mNeonGreen-p85α and mCherry-FAK on FAs from the cell. Right bottom: Plots showing the average intensity of mNeonGreen-p85α in cells treated with DMSO, defactinib, or alpelisib (n = 12, 10, and 12 T47D cells; n = 12, 18, and 19 MDA-MB-231 cells).

(E and F) T47D (E) or MDA-MB-231 (F) cells transiently expressing p110α-mNeonGreen and mCherry-FAK were imaged at 1-s intervals, with DMSO, defactinib, or alpelisib added (set as 0 s) during continuous imaging. Plots show the relative numbers of p110α-mNeonGreen molecules recruited to the plasma membrane in cells treated with DMSO, defactinib, or alpelisib (n = 8, 8, and 8 T47D cells; n = 8, 8, and 8 MDA-MB-231 cells).

(G) SUM159 cells transiently expressing mNeonGreen-p85β, the PI(3,4,5)$P_3$ sensor Halo-2xBtk(2-166) (labeled with $JFX_{650}$-HaloTag ligand) and mCherry-FAK were imaged at 10-s intervals, with defactinib added (set as 0 s) during continuous imaging. Left: Images at 0 s and 120 s after defactinib treatment in a representative time series. Middle: Kymographs generated along the line showing dynamics of mNeonGreen-p85β, PI(3,4,5)$P_3$ sensor, and mCherry-FAK. Right top: Plots showing the relative intensity of mNeonGreen-p85β, PI(3,4,5)$P_3$ sensor, and mCherry-FAK on FAs from the cell. Right bottom: Plots showing the average intensity of mNeonGreen-p85β on FAs in cells treated with DMSO, defactinib, or alpelisib (n = 16, 14, and 12 cells).

(H) SUM159 cells transiently expressing p110β-mNeonGreen, mCherry-FAK, and the PI(3,4,5)$P_3$ sensor Halo-2xBtk(2-166) (labeled with $JFX_{650}$-HaloTag ligand) were imaged at 1-s intervals, with DMSO, defactinib, or p110β inhibitor TGX-221 added (set as 0 s) during continuous imaging. Plots show the relative numbers of p110β-mNeonGreen molecules recruited to the plasma membrane in cells treated with DMSO, defactinib, or TGX-221 (n = 7, 8, and 6 cells).

(I) mEGFP-p85$\alpha^{+/+}$ cells were treated with control shRNA, or shRNA targeting FAK or PYK2. The expression of FAK and PYK2 in the cells was analyzed by western blot with the indicated antibodies.

Cells were imaged at the bottom surface by TIRF microscopy in (A-H). Data are shown as mean ± SEM in (A-H). Scale bars, 10 μm.

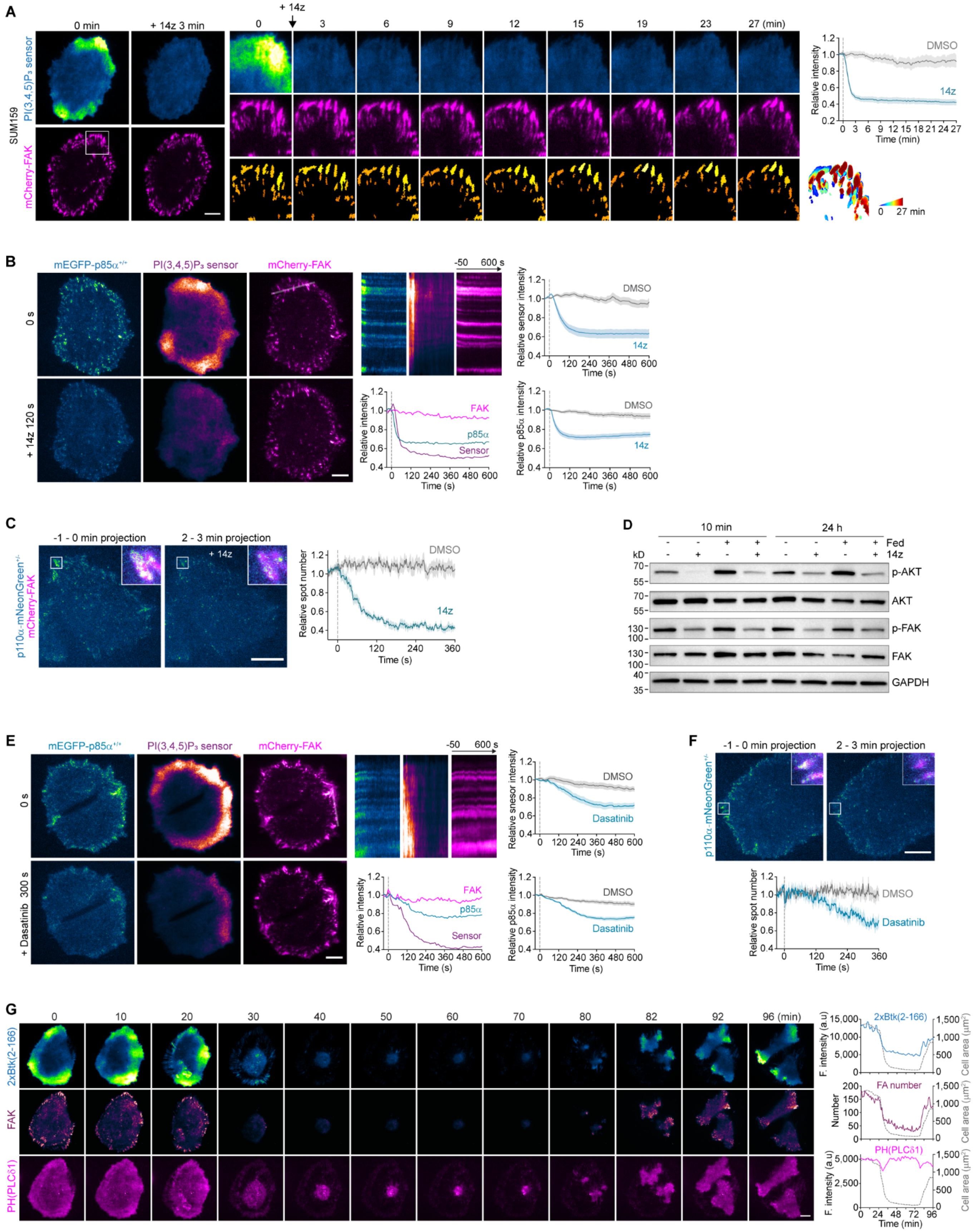
A
0 min
+ 14z 3 min
+ 14z
0 3 6 9 12 15 19 23 27 (min)
SUM159
PI(3,4,5)P₃ sensor
mCherry-FAK
Relative intensity
DMSO
14z
Time (min)
0 27 min
B
mEGFP-p85α+/+
PI(3,4,5)P₃ sensor
mCherry-FAK
0 s
+ 14z 120 s
-50 600 s
Relative sensor intensity
Relative intensity
FAK
p85α
Sensor
Relative p85α intensity
Time (s)
C
-1 - 0 min projection
2 - 3 min projection
+ 14z
p110α-mNeonGreen+/-
mCherry-FAK
Relative spot number
D
10 min
24 h
Fed
14z
kD
p-AKT
AKT
p-FAK
FAK
GAPDH
E
0 s
+ Dasatinib 300 s
Relative snesor intensity
Dasatinib
F
G
2xBtk(2-166)
FAK
PH(PLCδ1)
0 10 20 30 40 50 60 70 80 82 92 96 (min)
F. intensity (a.u)
Cell area (μm²)
FA number
Number
Time (min)

**Figure S9. Regulation of class IA PI3K recruitment and activation by the activated FAK, related to Figure 4**

(A) Bottom surfaces of SUM159 cells transiently expressing the PI(3,4,5)$P_3$ sensor mEGFP-2xBtk(2-166) and mCherry-FAK were imaged at 10-s intervals, with FAK inhibitor 14z added (set as 0 min) during continuous imaging. Left: Images at 0 min and 3 min after 14z treatment in a representative time series. Middle: Montages showing the indicated frames of the boxed regions, with detected FAs plotted at the bottom. Right: Plots showing the average intensity of PI(3,4,5)$P_3$ sensor on FAs from cells treated with DMSO or 14z (n = 9 and 16 cells).

(B) mEGFP-p85α$^{+/+}$ cells transiently expressing mCherry-FAK and the PI(3,4,5)$P_3$ sensor Halo-2xBtk(2-166) (labeled with $JFX_{650}$-HaloTag ligand) were imaged at 10-s intervals, with 14z added (set as 0 s) during continuous imaging. Left: Images at 0 s and 120 s after 14z treatment from a representative time series. Middle: Kymographs generated along the line on mCherry-FAK showing the recruitment dynamics of mEGFP-p85α and PI(3,4,5)$P_3$ sensor to FAs; plots showing the relative intensity of mEGFP-p85α, PI(3,4,5)$P_3$ sensor, and mCherry-FAK on FAs over time. Right: Plots showing the average intensity of PI(3,4,5)$P_3$ sensor (top) or mEGFP-p85α (bottom) on FAs in cells treated with DMSO or 14z (n = 10 and 16 cells).

(C) p110α-mNeonGreen$^{+/-}$ cells transiently expressing mCherry-FAK and the PI(3,4,5)$P_3$ sensor Halo-2xBtk(2-166) were imaged at 1-s intervals, with DMSO or 14z added (set as 0 s) during continuous imaging. The maximum-intensity projections of the 60 frames before and 1 minute after 14z treatment in the time series are shown, with the p110α and FAK overlaid image of the boxed region shown in the corner. Plots show the relative numbers of p110α-mNeonGreen molecules recruited to the plasma membrane in cells treated with DMSO or 14z (n = 8 and 10 cells).

(D) Cells were treated with DMSO or 14z for 10 min after culturing in complete medium (Fed) or starvation medium for 24 hours, or the cells were treated with DMSO or 14z for 24 hours in complete medium or starvation medium. The total and phosphorylated AKT and FAK levels were analyzed by western blot.

(E) mEGFP-p85α$^{+/+}$ cells transiently expressing mCherry-FAK and the PI(3,4,5)$P_3$ sensor Halo-2xBtk(2-166) (labeled with $JFX_{650}$-HaloTag ligand) were imaged at 10-s intervals, with Src inhibitor dasatinib added (set as 0 s) during continuous imaging. Left: Images at 0 s and 300 s after dasatinib treatment from a representative time series. Middle: Kymographs generated along the line on mCherry-FAK showing the recruitment dynamics of mEGFP-p85α and PI(3,4,5)$P_3$ sensor; plots showing the relative intensity of mEGFP-p85α, PI(3,4,5)$P_3$ sensor, and mCherry-FAK on FAs over time. Right: Plots showing the average intensity of PI(3,4,5)$P_3$ sensor (top) or mEGFP-p85α (bottom) on FAs in cells treated with DMSO or dasatinib (n = 14 and 15 cells).

(F) p110α-mNeonGreen$^{+/-}$ cells transiently expressing mCherry-FAK and the PI(3,4,5)$P_3$ sensor Halo-2xBtk(2-166) were imaged at 1-s intervals, with DMSO or dasatinib added (set as 0 s) during continuous imaging. The maximum-intensity projections of the 60 frames before and 1 minute after dasatinib treatment in the time series are shown. The boxed region is enlarged at the top right, with the p110α and FAK images overlaid. Plots show the relative numbers of p110α-mNeonGreen molecules recruited to the plasma membrane in cells treated with DMSO or dasatinib (n = 8 and 8 cells).

(G) SUM159 cells transiently expressing the PI(3,4,5)$P_3$ sensor mScarlet-I-2xBtk(2-166), the PI(4,5)$P_2$ sensor mEGFP-PH(PLCδ1), and Halo-FAK (labeled with $JFX_{650}$-HaloTag ligand) were imaged at 60 s intervals by TIRF microscopy. Left: Montage showing the distribution of mScarlet-I-2xBtk(2-166), mEGFP-PH(PLCδ1), and Halo-FAK at the plasma membrane of the cells during mitosis. Right: Time courses showing the fluorescence intensity of mScarlet-I-2xBtk(2-166) and mEGFP-PH(PLCδ1), the number of FAs, and the total areas of the cells during mitosis.

Cells were imaged at the bottom surface by TIRF microscopy in (A-C), and (E-G). Data are shown as mean ± SEM in (A-C), (E), and (F). Scale bars, 10 μm.

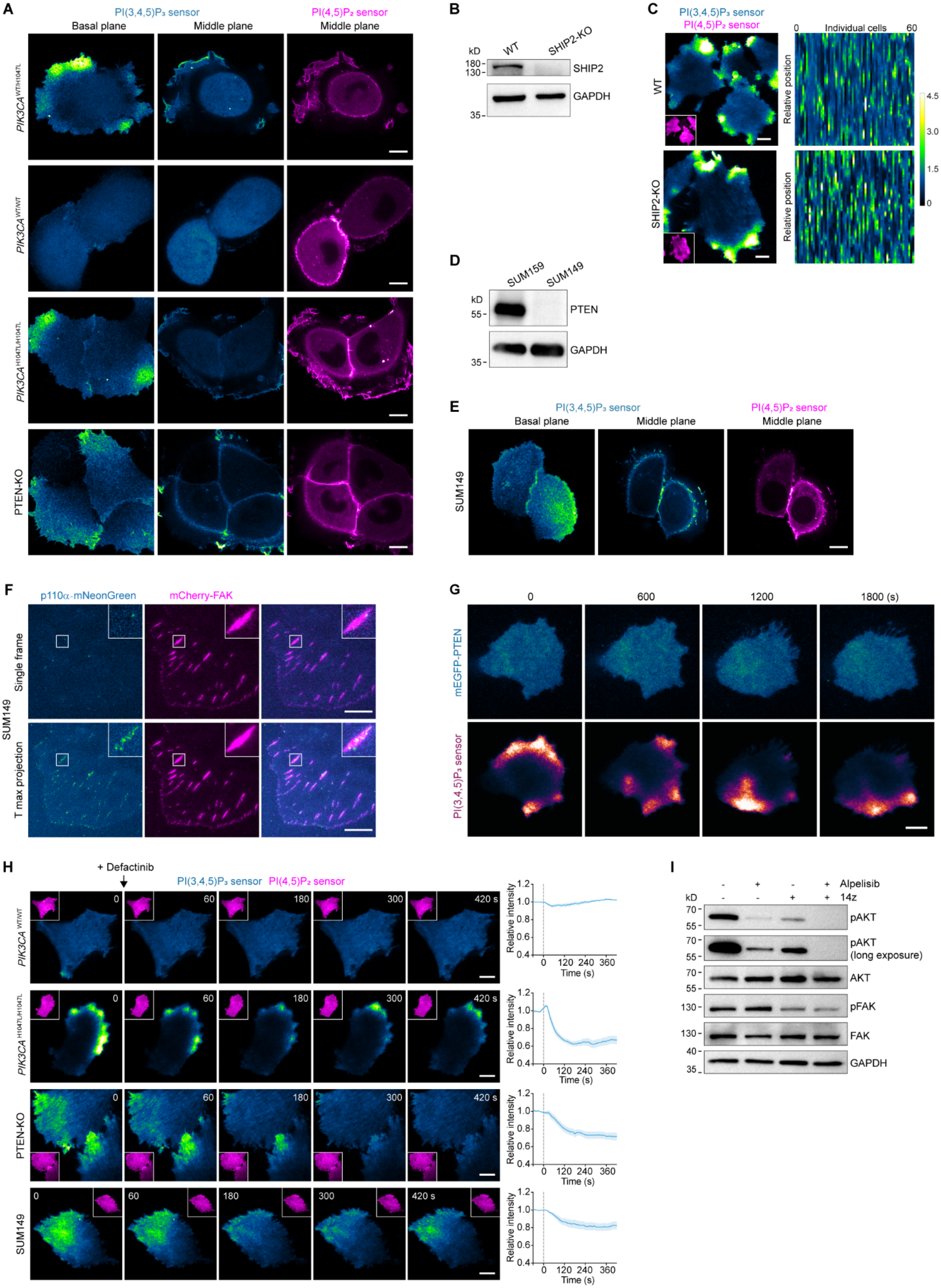
A
PI(3,4,5)P₃ sensor
PI(4,5)P₂ sensor
Basal plane
Middle plane
Middle plane
PIK3CA WT/H1047L
PIK3CA WT/WT
PIK3CA H1047L/H1047L
PTEN-KO
B
kD
WT
SHIP2-KO
180
130
SHIP2
GAPDH
35
C
PI(3,4,5)P₃ sensor
PI(4,5)P₂ sensor
0
Individual cells
60
WT
SHIP2-KO
Relative position
4.5
3.0
1.5
0
D
SUM159
SUM149
kD
55
PTEN
GAPDH
35
E
PI(3,4,5)P₃ sensor
PI(4,5)P₂ sensor
Basal plane
Middle plane
Middle plane
SUM149
F
p110α-mNeonGreen
mCherry-FAK
Single frame
T max projection
SUM149
G
0
600
1200
1800 (s)
mEGFP-PTEN
PI(3,4,5)P₃ sensor
H
+ Defactinib
PI(3,4,5)P₃ sensor PI(4,5)P₂ sensor
PIK3CA WT/WT
PIK3CA H1047L/H1047L
PTEN-KO
SUM149
0
60
180
300
420 s
Relative intensity
Time (s)
0 120 240 360
I
Alpelisib
14z
kD
70
55
pAKT
pAKT (long exposure)
AKT
130
pFAK
FAK
40
35
GAPDH

**Figure S10. Regulation of $PI(3,4,5)P_3$ distribution at the plasma membrane by class I PI3K and PTEN, related to Figures 5 and 6**

(A) $\mathit{PIK3CA}^{WT/H1047L}$, $\mathit{PIK3CA}^{WT/WT}$, $\mathit{PIK3CA}^{H1047L/H1047L}$, or PTEN-KO SUM159 cells transiently expressing the $PI(3,4,5)P_3$ sensor mEGFP-2xBtk(2-166) and the $PI(4,5)P_2$ sensor mScarlet-I-PH(PLCδ1) were imaged by spinning-disk confocal microscopy. Representative images show the distribution of both sensors at the basal and middle planes of the cells.

(B) Expression of SHIP2 was eliminated in SUM159 cells by CRISPR/Cas9-targeted knockout of SHIP2 (SHIP2-KO), as confirmed by western blot using antibodies against SHIP2 and GAPDH.

(C) WT and SHIP2-KO SUM159 cells transiently expressing the $PI(3,4,5)P_3$ sensor mEGFP-2xBtk(2-166) and the $PI(4,5)P_2$ sensor mScarlet-I-PH(PLCδ1) were imaged by TIRF microscopy. Left: Representative images of mEGFP-2xBtk(2-166) and mScarlet-I-PH(PLCδ1) (inserts). Right: Intensity profile heatmaps show the distribution of $PI(3,4,5)P_3$ sensor at the plasma membrane of WT and SHIP2-KO cells (n = 60 cells each).

(D) SUM149 cells lack PTEN expression, as confirmed by western blot using antibodies against PTEN and GAPDH.

(E) SUM149 cells transiently expressing the $PI(3,4,5)P_3$ sensor mEGFP-2xBtk(2-166) and the $PI(4,5)P_2$ sensor mScarlet-I-PH(PLCδ1) were imaged by spinning-disk confocal microscopy. Representative images show the distribution of both sensors at the basal and middle planes of the cells.

(F) SUM149 cells transiently expressing p110α-mNeonGreen and mCherry-FAK were imaged at 0.2-s intervals for 601 frames. Images of a single frame and maximum-intensity projection of a representative time series are shown.

(G) SUM159 cells transiently expressing mEGFP-PTEN and the $PI(3,4,5)P_3$ sensor mScarlet-I-2xBtk(2-166) were imaged at 30-s intervals. The montage shows images at the indicated times of a representative time series.

(H) SUM159 cells ($\mathit{PIK3CA}^{WT/WT}$, $\mathit{PIK3CA}^{\mathrm{H1047L/H1047L}}$, and PTEN-KO) and SUM149 cells were transiently transfected with the $PI(3,4,5)P_3$ sensor mEGFP-2xBtk(2-166) and the $PI(4,5)P_2$ sensor mScarlet-I-PH(PLCδ1) (inserts) and then imaged at 10-s intervals by TIRF microscopy, with defactinib added (set as 0 s) during continuous imaging. Montages show images at the indicated times of a representative time series. Plots show the average intensity of $PI(3,4,5)P_3$ sensor in defactinib-treated cells (n = 6, 8, 8, and 8 cells).

(I) Cells were treated with DMSO, alpelisib, 14z, or both alpelisib and 14z for 24 hours. The total and phosphorylated AKT and FAK levels were analyzed by western blot with the indicated antibodies.

Data are shown as mean ± SEM in (H). Scale bars, 10 μm.
